# Elaborate expansion of syntenic V1R hotspots correlates with high species diversity in nocturnal mouse and dwarf lemurs

**DOI:** 10.1101/637348

**Authors:** Kelsie E. Hunnicutt, George P. Tiley, Rachel C. Williams, Peter A. Larsen, Marina B. Blanco, Rodin M. Rasoloarison, C. Ryan Campbell, Kevin Zhu, David W. Weisrock, Hiroaki Matsunami, Anne D. Yoder

**Affiliations:** Department of Biology, Duke University, Durham, NC 27708, USA; Duke Lemur Center, Duke University, Durham, NC 27705; Behavioral Ecology and Sociobiology Unit, German Primate Centre, 37077 Göttingen, Germany; Département de Biologie Animale, Université d’Antananarivo, BP 906, Antananarivo 101, Madagascar; Department of Molecular Genetics and Microbiology, Duke University Medical Center, Durham, NC 27710; Department of Biology, University of Kentucky, Lexington, KY 40506; Department of Neurobiology, Duke Institute for Brain Sciences, Duke University Medical Center, Durham, NC 27710; Department of Biological Sciences, University of Denver, Denver, CO 80208; Department of Veterinary and Biomedical Sciences, University of Minnesota, Saint Paul, MN 55108

**Author notes:** Equal contributors (Kelsie E. Hunnicutt, George P. Tiley). Author for Correspondence: Anne D. Yoder, Department of Biology, Duke University, Durham, USA.

**Keywords:** speciation, pheromone receptors, synteny, gene family evolution, nocturnality, vomeronasal

## Abstract

Sensory gene families are of special interest, both for what they can tell us about molecular evolution, and for what they imply as mediators of social communication. The vomeronasal type-1 receptors (V1Rs) have often been hypothesized as playing a fundamental role in driving or maintaining species boundaries given their likely function as mediators of intraspecific mate choice, particularly in nocturnal mammals. Here, we employ a comparative genomic approach for revealing patterns of V1R evolution within primates, with a special focus on the small-bodied nocturnal mouse and dwarf lemurs of Madagascar (genera *Microcebus* and *Cheirogaleus*, respectively). By doubling the existing genomic resources for strepsirrhine primates (i.e., the lemurs and lorises), we find that the highly-speciose and morphologically-cryptic mouse lemurs have experienced an elaborate proliferation of V1Rs that we argue is functionally related to their capacity for rapid lineage diversification. Contrary to a previous study that found equivalent degrees of V1R diversity in diurnal and nocturnal lemurs, our study finds a strong correlation between nocturnality and V1R elaboration, with nocturnal lemurs showing elaborate V1R repertoires and diurnal lemurs showing less diverse repertoires. Recognized subfamilies among V1Rs show unique signatures of diversifying positive selection, as might be expected if they have each evolved to respond to specific stimuli. Further, a detailed syntenic comparison of mouse lemurs with mouse (genus *Mus*) and other mammalian outgroups shows that orthologous mammalian subfamilies, predicted to be of ancient origin, tend to cluster in a densely populated region across syntenic chromosomes that we refer to as V1R “hotspots.”

## Introduction

The evolutionary dynamics of sensory gene families are of fundamental interest as a model for how molecular evolutionary processes can shape the content and structure of genomes and for their ability to characterize the life history and ecological traits of organisms. Vomeronasal type-1 receptor genes (V1Rs) comprise one such gene family and have been the subject of increasing interest in both the molecular genetics (e.g., Adipietro et al. 2012) and evolutionary genetics (e.g., Yohe and Brand 2018) communities. Vomerolfaction is a form of chemosensation that mediates semiochemical detection and occurs in the vomeronasal organ (VNO) of mammals (Leinders-Zufall et al. 2000). V1Rs are expressed on vomeronasal sensory neurons in the VNO and have been demonstrated to detect pheromones in mice (Boschat et al. 2002; Haga-Yamanaka et al. 2014). For example, impaired vomeronasal function in mice, either through a knockout of V1Rs or removal of the VNO, alters appropriate chemosensory behaviors such as conspecific avoidance of sick animals, interspecies defensive cues, male sexual behavior, and maternal aggression (Del Punta et al. 2002; Papes et al. 2010; Boillat et al. 2015). Thus, the evolution of V1Rs can have direct consequences for both the emitter and the receiver of pheromone signals, with ample evidence indicating that molecular evolution of V1Rs is associated with the speciation process (Lane et al. 2002; Kurzweil et al. 2009; Hohenbrink et al. 2012; Nikaido et al. 2014).

V1Rs are ideally suited for study within the context of “sensory drive” wherein mate preferences in communication systems diverge in the face of novel environmental opportunity (Boughman 2002). Communication mechanisms for mate recognition have been recognized as an important component for driving rapid reproductive isolation (Mendelson 2003; Dopman et al. 2010; Servedio and Boughman 2017; Brand et al. 2019) and with the reinforcement of species boundaries (Servedio and Noor 2003). Sensory drive can affect diverging populations in two ways by targeting the pheromone receptors and/or their signaling molecules. As examples, adaptation of a likely V1R signal in mice, androgen binding protein, is associated with assortative mating between *Mus musculus* subspecies (Karn et al. 2010; Chung et al. 2017; Hurst et al. 2017) just as fixation of nonsynonymous polymorphisms among V1Rs is associated with the speciation of *Mus spretus* and *Mus musculus* (Kurzweil et al. 2009). Moreover, it has been shown that differential expression of vomeronasal and olfactory receptor genes, including V1Rs, is associated with assortative mating in a pair of house mouse subspecies (Loire et al. 2017) and is likely reinforcing the subspecies along their hybrid zone.

The V1R gene family has experienced many duplications and losses in the evolutionary history of mammals, and the availability of duplicate copies can allow for divergence among sequences, gene expression, and ultimately function (e.g. Lynch and Conery 2000; Des Marais and Rausher 2008). Though not directly addressed in this study, it is worth noting that changes in gene expression often occur rapidly after gene duplication events (Makova and Li 2003; Keller and Yi 2014; Guschanski et al. 2017) and are often accompanied by shifts in rates of molecular evolution (Chen et al. 2010; Yang and Gaut 2011). Although the mechanisms that explain variable rates of molecular evolution, specifically the nonsynonymous to synonymous substitution rate ratio (*dN*/*dS*), are complex, there is some interdependence on expression levels (O’Toole et al. 2018) and genome architecture (Dai et al. 2014; Xie et al. 2016). The V1R gene family demonstrates structural complexity (Ohara et al. 2009; Yohe et al. 2018), and gene family expansions and directional selection acting on duplicate copies may be important for the maintenance of species boundaries where vomerolfaction is linked with assortative mating (Luo et al. 2003; Isogai et al. 2011; Fu et al. 2015).

Here, we present a comparative genomic study of V1R evolution within the lemuriform primates, primarily focusing on the mouse lemurs of Madagascar (genus *Microcebus*). Mouse lemurs are perhaps the most species-rich clade of living primates (Hotaling et al. 2016), and are well-known for high levels of interspecific genetic divergence though with nearly uniform morphological phenotypes. They have thus come to be regarded as a classic example of a cryptic species radiation, perhaps related to their nocturnal lifestyle (Yoder et al. 2016). Mouse lemurs, and the closely-related dwarf lemurs, have elaborate olfactory communication behaviors that are associated with adaptive strategies such as predator recognition (Sündermann et al. 2008), fecundity (Drea 2015), and even biased sex ratios (Perret 1996; Perret and Colas 1997). V1Rs take on particular interest in mouse lemurs as we hypothesize that their observed role in both speciation and in the maintenance of species boundaries within *Mus* may also apply to this speciose clade of primates (Smadja et al. 2015; Loire et al. 2017). We hypothesize that among primates, mouse lemurs will show signatures of sensory drive via genomic elaboration of the V1R complex and evidence of positive selection acting on V1R genes.

There are numerous lines of evidence to lead us to this hypothesis: 1) Previous studies have indicated that V1Rs within the lemuriform clade have evolved under pervasive positive selection 5/22/19 3:35:00 PM, 2) that the majority of gene copies are intact (Young et al. 2010; Larsen et al. 2014), and 3) that the differential expression of a large number of vomeronasal receptors in both the VNO and main olfactory epithelium of mouse lemurs are associated with different behaviors and chemical signals (Hohenbrink et al. 2014). In fact, along with murids, opossums, and platypus, mouse lemurs have been reported to have among the largest V1R repertoires found in mammals (Young et al. 2010). Even so, numerous obstacles such as complexities of chemical background, chemical signals, and the genetic basis of chemosensation complicate both ecological and experimental approaches for differentiating between cause and effect in the speciation process (Yohe and Brand 2018). This is particularly problematic for studies of mouse lemurs given their remote geographic distribution, nocturnal life history, and endangered status. Thus, we take a comparative genomic approach for reconstructing the evolutionary dynamics of the V1R gene family within the small-bodied and nocturnal mouse and dwarf lemurs (family, Cheirogaleidae).

### A Comparative Genomic Approach

V1R loci are highly repetitive and they, along with their surrounding regions, are notoriously challenging for genome assembly. Though previous studies have used targeted sequencing or short-read sequencing to examine the evolutionary dynamics of V1R expansions in a limited number of species (Young et al. 2010; Hohenbrink et al. 2012; Yoder et al. 2014), strepsirrhine primates have until recently remained woefully underrepresented in genomic databases (Perry et al. 2012; Meyer et al. 2015; Larsen et al. 2017; Hawkins et al. 2018). Here, we take advantage of the chromosome-level assembly of the gray mouse lemur, *Microcebus murinus*, along with short-read sequencing in related species, to characterize the V1R repertoires for lemuriform primates. Recent improvements using long-read sequencing of the mouse lemur genome (Larsen et al. 2017) improve our ability to characterize the V1R repertoire (Larsen et al. 2014) and allow for comparisons of the genomic architecture of V1R-containing regions in expanded and contracted V1R repertoires across mammals.

In this study, we have sequenced and assembled seven new cheirogaleid genomes, with a particular focus on the mouse lemurs. Further, to explore intraspecific copy number variation and evaluate the effects of assembly error on V1R repertoire counts, we resequenced and *de novo* assembled genomes from eight *M*. *murinus* individuals from a captive breeding colony. Our study thus serves as a timely companion to two recent overviews of comparative genomic studies for illuminating the evolutionary and life-history dynamics of chemosensory gene family evolution in vertebrates (Bear et al. 2016; Hughes et al. 2018). A comparative genomic approach allows us to explore classic predictions of gene-family evolution, namely, that genomic drift can operate at very fine scales to produce high intraspecific copy number variation (Nozawa et al. 2007) and that gene-family evolution is often marked by a strong birth-death process over phylogenetic time scales (Nei et al. 1997; Csűrös and Miklos 2009; Hughes et al. 2018). The latter question is of particular interest for V1R evolution given that adaptive pressures on these genes makes them highly vulnerable to pseudogenization in cases of relaxed selection, thus yielding the observed correlations between levels of V1R ornamentation and diverse adaptive regimes. An overview of primates shows that those with elaborate representation of subfamilies have a strong reliance on chemosensory communication whereas those with depauperate V1R representation rely on alternative mechanisms for inter- and intra-specific communication (Yoder and Larsen 2014).

These new genomic resources have also allowed us to address a number of questions regarding rates of molecular evolution in V1Rs. Divergent gene function following gene duplication predicts that some signature of positive selection should be evident in the gene sequences (Zhang et al. 1998), but it remains unknown if selection has acted pervasively over time or has occurred in episodic bursts prior to the diversification of mouse lemurs. We might anticipate episodic positive selection to be the primary mechanism if purifying selection has been operating at more recent time scales to preserve gene function among duplicate copies. For strepsirrhine primates (i.e., the lemurs and lorises), pervasive positive selection has been detected at the interspecific level (Hohenbrink et al. 2012; Yoder et al. 2014), while strong purifying selection has been found within populations. Here we disentangle pervasive versus episodic positive selection among V1Rs and show that both gene duplication and rates of molecular evolution have been active in shaping expanded V1R repertoires among the dwarf and mouse lemurs. Moreover, through comparison with *Mus* and other mammals, we show that orthologous subfamilies tend to cluster in a densely populated region on syntenic chromosomes that we refer to as V1R “hotspots.”

## Results and Discussion

### Novel genome assemblies of several strepsirrhine primates

We *de novo* assembled seven novel strepsirrhine genomes: *Microcebus griseorufus, M. ravelobensis, M. mittermeieri, M. tavaratra, Mirza zaza, Cheirogaleus sibreei* and *C. medius*. These efforts have doubled the number of publicly available genomes for the Strepsirrhini with a specific focus on the dwarf and mouse lemur clade. Excluding *C. medius*, the seven genomes were sequenced to an average depth of coverage between 26x and 45x with scaffold N50s of 17-76kb (Supplementary Table S1). The *C. medius* reference genome was assembled using Dovetail Genomics to an average depth of coverage of 110x and a scaffold N50 of approximately 50Mb (Williams et al. 2019). We evaluated assembly completeness using the Benchmarking Universal Single-Copy Orthologs tool, BUSCO (Simão et al. 2015), which assesses genomes for the presence of complete near-universal single-copy orthologs (Supplementary Figure S1). The assemblies recovered between 77.2% and 92.3% of the mammalian BUSCO gene set. We also resequenced eight *M. murinus* individuals, with one duplicate individual (Campbell et al. 2019), and here have *de novo* assembled genomes for each individual with 21x-29x effective coverage using the 10x Genomics Supernova pipeline. The additional scaffolding information provided by the 10x Genomics linked-reads resulted in scaffold N50s of 0.6-1.2 Mb. BUSCO analyses revealed that the resequenced assemblies recovered between 89.9% and 95.5% of the mammalian gene set. A denser sampling of genomes within Cheirogaleidae not only provides an opportunity for illuminating patterns of V1R gene family evolution but also promotes greater understanding of the molecular evolution of primate and strepsirrhine-specific genomes. Genome resequencing of *M. murinus* individuals has allowed investigation of intraspecific V1R copy number variation as well as questions regarding microevolutionary processes and gene family evolution (Park et al. 2011).

The monophyletic genus *Microcebus* contains 24 named species (Hotaling et al. 2016), and our results clearly demonstrate that the clade has a uniquely complex V1R repertoire compared to other primates thus far characterized (Figure 1A and B). Contrary to a previous study suggesting that V1R expansion is ubiquitous across the lemuriform clade (Yoder et al. 2014), increased sampling reveals that expansion has been profound in the nocturnal dwarf and mouse lemurs. This is consistent with the original hypothesis that local V1R expansions may play a role in forming or maintaining speciation boundaries within *Cheirogaleus* and *Microcebus* as might be predicted given their nocturnal lifestyle. Phylogenetic analyses revealed that expanded V1R repertoires in mouse lemurs demonstrate a remarkably higher rate of duplicate gene retention in comparison to other primates (Figure 1A and B; Table 1). The common ancestor of mouse lemurs is not associated with novel subfamily birth though the diversity and number of V1R gene copies is striking (Figure 1A; Table 1). Although genomes generated exclusively from short-read data are vulnerable to collapsing loci in assemblies (Larsen et al. 2014), our inference of increased V1R retention in *M. murinus* relative to non-cheirogaleid primates was robust to assembly strategies and data sources (Supplementary Table S2). Further, the resequenced *M. murinus* individuals reveal low intraspecific variation in copy number (Figure 2), suggesting that the observed differences in repertoire size between mouse lemurs and other non-nocturnal lemurs is not an artifact of individual sampling or assembly error (Figure 3).

**Figure 1.**
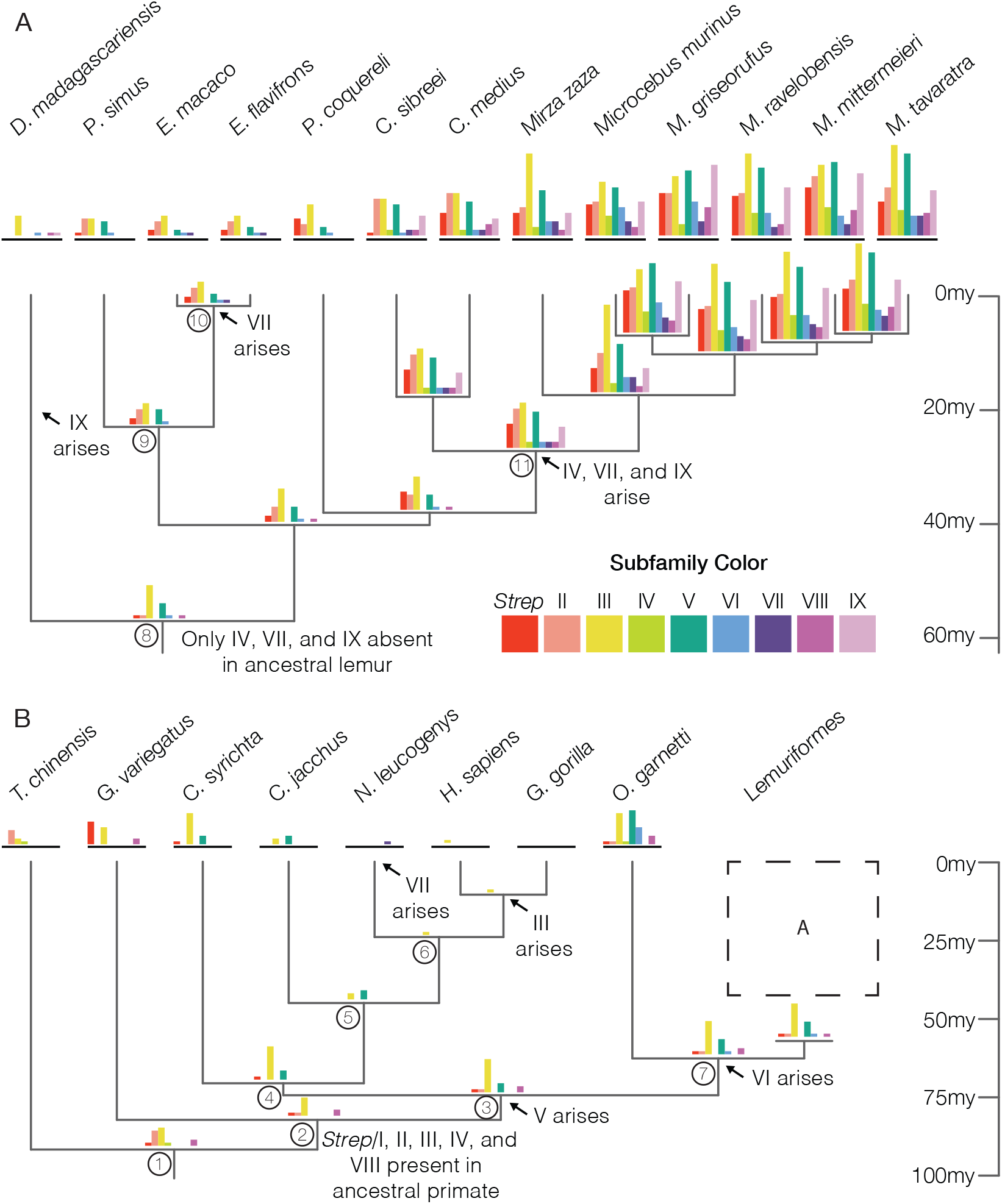
V1R subfamily membership across the primate phylogeny. Membership was assessed for available strepsirrhine genomes (A) and for select primate outgroups (B) and estimated using the Count software for ancestral lineages. Bar graphs show absolute gene count for each subfamily. Predicted gene subfamily origins are annotated with arrows. Tree adapted from dos Reis 2018 (dos Reis et al. 2018). Circled node numbers correspond with Table 1.

**Figure 2.**
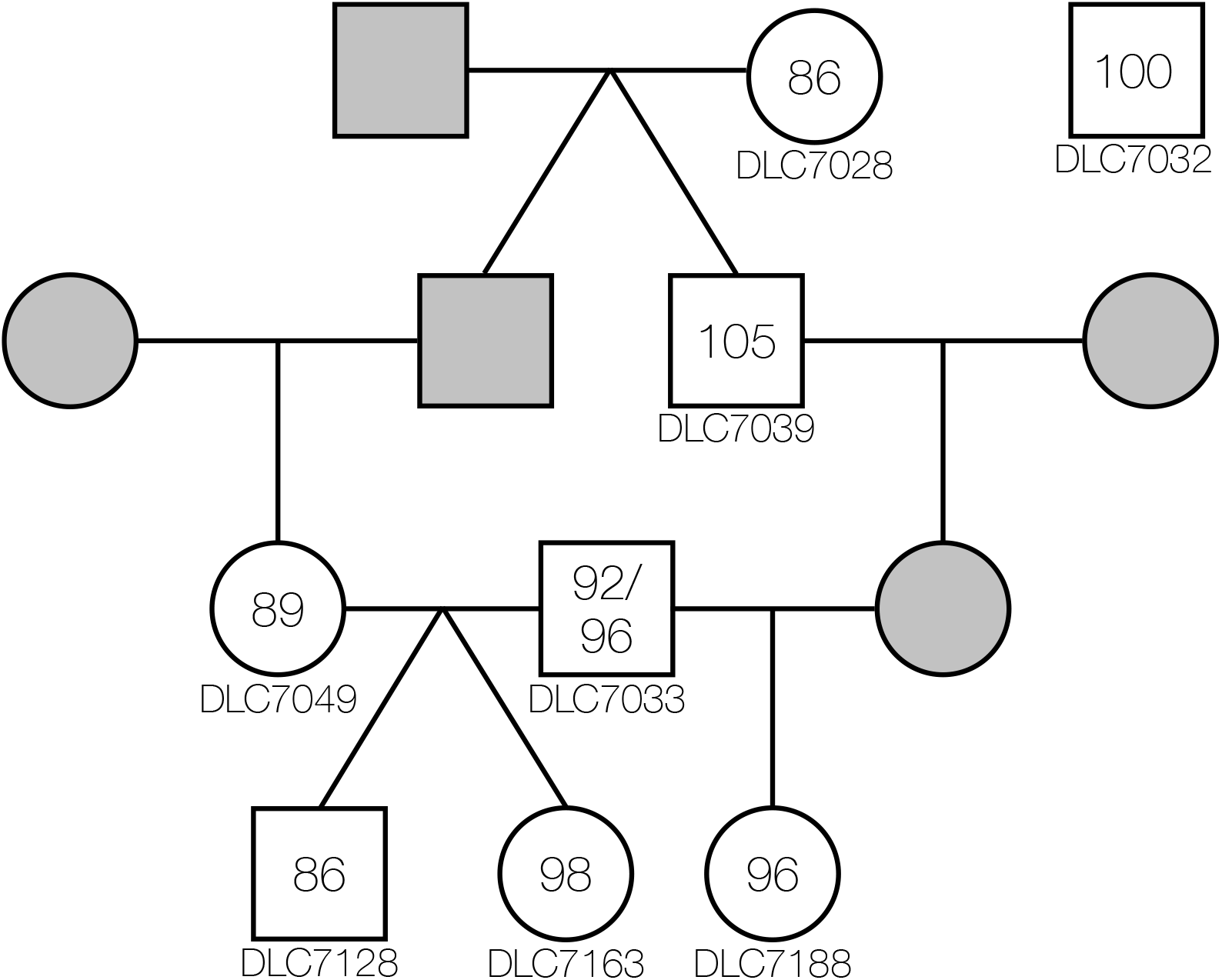
Intraspecific variation in V1R repertoire size estimates across eight closely-related *Microcebus murinus* individuals. Genomes were *de novo* assembled and mined for loci with significant V1R homology and an ORF longer than 801bp. Individual DLC7033 was sequenced twice and repertoire size estimates are reported for both assemblies. Squares represent males and circles represent females. Horizontal lines indicate mate pairs (mother and father) and vertical or slanted lines indicate parent to offspring relationshiop. Numbers inside the symbols represent repertoire size estimates. Individuals represented by grey symbols were not sequenced.

**Figure 3.**
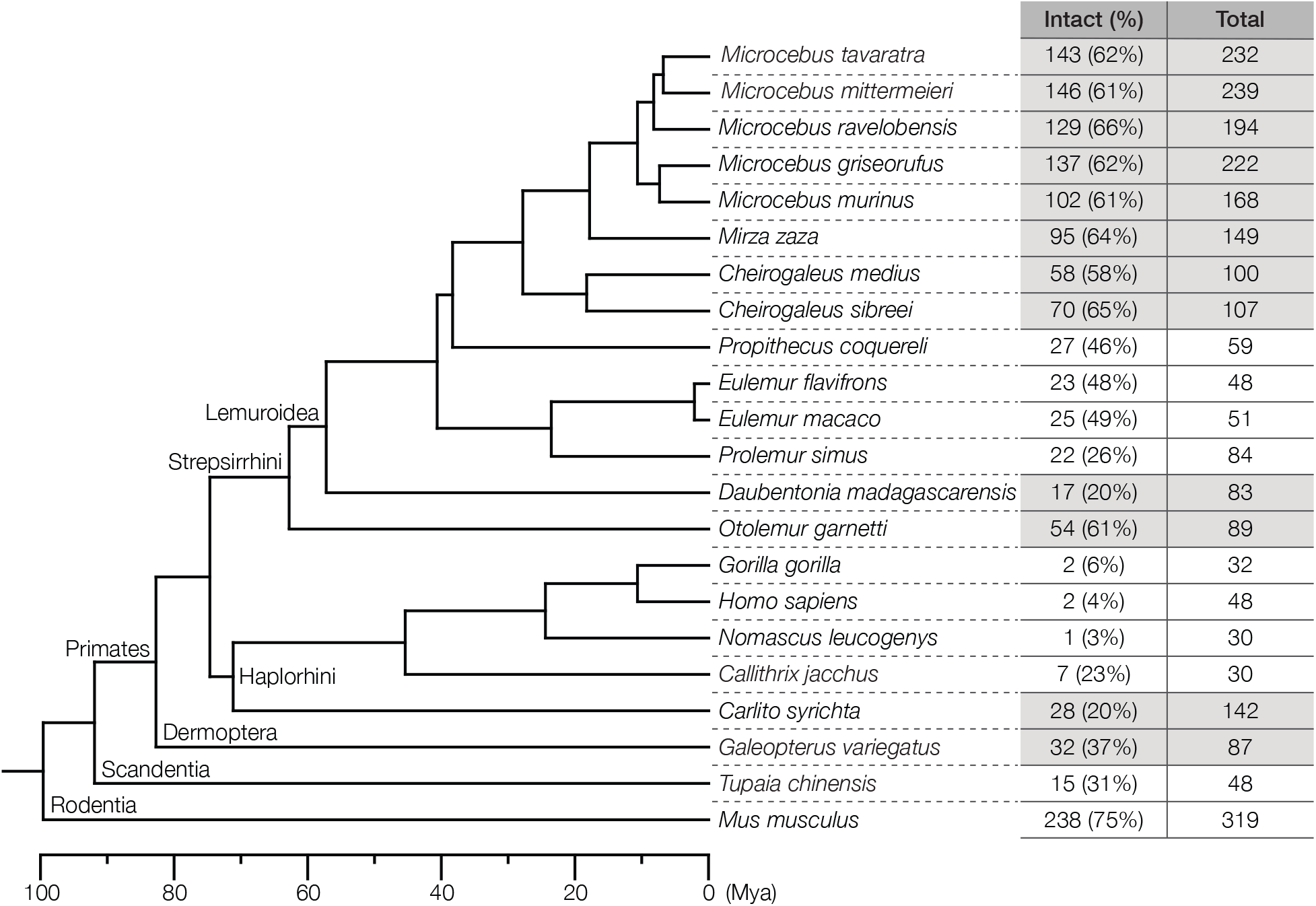
V1R repertoire size estimates across the strepsirrhine phylogeny. Sequences with V1R homology were mined from available strepsirrhine and select outgroup genomes. Total V1Rs consist of all genomic regions with V1R homology that are ≥ 600bp in length. Intact genes are defined by vomeronasal homology and a ≥ 801bp ORF. Nocturnal species are highlighted in gray. Tree adapted from dos Reis 2018 (dos Reis et al. 2018).

**Table 1.**
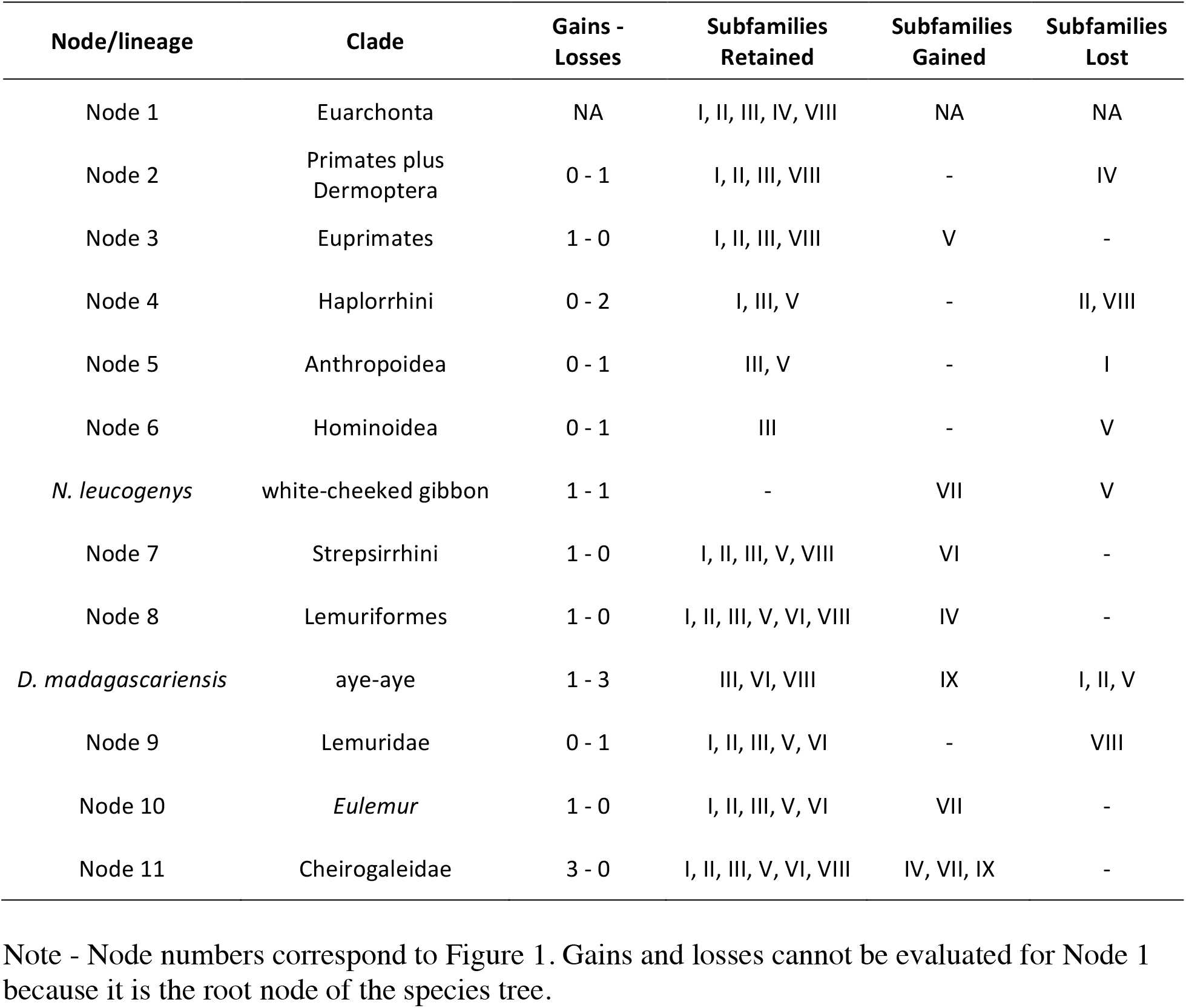
Inference of V1R birth-death process within primates.

| Node/lineage | Clade | Gains - Losses | Subfamilies Retained | Subfamilies Gained | Subfamilies Lost |
| --- | --- | --- | --- | --- | --- |
| Node 1 | Euarchonta | NA | I, II, III, IV, VIII | NA | NA |
| Node 2 | Primates plus Dermoptera | 0 - 1 | I, II, III, VIII | - | IV |
| Node 3 | Euprimates | 1 - 0 | I, II, III, VIII | V | - |
| Node 4 | Haplorrhini | 0 - 2 | I, III, V | - | II, VIII |
| Node 5 | Anthropoidea | 0 - 1 | III, V | - | I |
| Node 6 | Hominoidea | 0 - 1 | III | - | V |
| *N. leucogenys* | white-cheeked gibbon | 1 - 1 | - | VII | V |
| Node 7 | Strepsirrhini | 1 - 0 | I, II, III, V, VIII | VI | - |
| Node 8 | Lemuriformes | 1 - 0 | I, II, III, V, VI, VIII | IV | - |
| *D. madagascariensis* | aye-aye | 1 - 3 | III, VI, VIII | IX | I, II, V |
| Node 9 | Lemuridae | 0 - 1 | I, II, III, V, VI | - | VIII |
| Node 10 | *Eulemur* | 1 - 0 | I, II, III, V, VI | VII | - |
| Node 11 | Cheirogaleidae | 3 - 0 | I, II, III, V, VI, VIII | IV, VII, IX | - |

Note - Node numbers correspond to Figure 1. Gains and losses cannot be evaluated for Node 1 because it is the root node of the species tree.

The expansion dynamics of V1Rs within Cheirogaleidae do not support a simple linear correlation between species richness and repertoire size. Although all cheirogaleid repertoires had full primate subfamily membership, there was variation in subfamily proportions between species, which is consistent with our hypothesis that species-specific V1R repertoires and chemosensation may be important for species diversity of cheirogaleids in comparison to diurnal strepsirrhines. Dwarf lemurs, genus *Cheirogaleus*, are hypothesized to have as many as 18 species (Lei et al. 2014) though have the smallest V1R repertoires within the cheirogaleids examined here. Conversely, the genus *Mirza*, with only two recognized species, has a repertoire size that is nearly equal to that of *Microcebus murinus*. It is notable, however, that the *Mirza* genome’s expanded repertoire is primarily enriched for subfamily III (Figure 1A). The differential subfamily enrichment among species suggests that despite the similarity in size to *Microcebus* repertoires, the V1R repertoire of *Mirza* has experienced independent selective pressures on gene retention and may ultimately fulfill a different functional role compared to *Microcebus*.

### V1R repertoire estimation across primates

We estimated V1R repertoire size evolution across strepsirrhine primates as well as for several well-annotated primate and mammalian genomes for outgroup comparison. Notably, repertoire estimates of extant primates are comparable to previous studies that used trace archive fragments and earlier draft genome versions (Figure 3; Supplemental Table S2; Young et al. 2010; Moriya-Ito et al. 2018). The expanded V1R repertoire within the gray mouse lemur is not ubiquitous across the Strepsirrhini, however. Rather, repertoire size expanded gradually from a reduced set in the strepsirrhine common ancestor to its peak in the mouse lemur clade. This expansion is characterized by a reduced repertoire in the early diverging aye-aye lineage (genus *Daubentonia*), moderate repertoires among diurnal lemurs, and an expansion that likely occurred in the common ancestor of Cheirogaleidae (Figure 3). If the origins of many V1R copies in mouse lemur date to the Cheirogaleidae common ancestor, this means that at least some of those duplicates have remained functional and intact since their origins 30 million years ago, as would be consistent with divergence time estimates for the cheirogaleid radiation (Yang and Yoder 2003; dos Reis et al. 2018).

Within Cheirogaleidae, repertoire sizes ranged from a low of 58 intact V1Rs in *C. medius* to highs between 102-143 intact V1Rs in the genus *Microcebus*. The mouse lemurs have universally large V1R repertoires (102-146 intact genes) with notable intragenus variation. Prior to this study, *M. murinus* had been identified as having one of the largest V1R repertoires within mammals (Young et al. 2010). Additional sampling from *Microcebus* reveals, however, that among the five mouse lemur species here characterized, *M. murinus* actually has the smallest repertoire with only 102 intact V1Rs. We also estimated the percent of intact V1Rs contained within the total repertoire for each species. Most haplorrhine primates (Anthropoidea plus *Tarsius*) species have repertoires with low percentages of intact receptors (<37% intact). Within Lemuroidae, the diurnal lemurs also have small and pseudogenized repertoires (26% to 49% intact) containing only 22-27 intact V1Rs. In contrast, among nocturnal species excluding aye-aye, we observe intact repertoires between 58% to 66% within Cheirogaleidae, and 61% for the nocturnal lorisiform *Otolemur garnetti*.

These comparisons do not, however, provide definitive evidence that expanded V1R repertoires in primates are strictly associated with nocturnal life history (Wang et al. 2010; Moriya-Ito et al. 2018). Although *Otolemur garnetti* shows a proportion of intact V1R copies similar to dwarf and mouse lemurs (Figure 3), subfamilies VII and IX are absent from *O. garnetti* (Figure 1B). By comparison, the genomes of both the aye-aye and the tarsier (Schmitz et al. 2016) contain low numbers of intact V1R gene copies, which appears to contradict the hypothesis that a nocturnal life history alone is sufficient for explaining V1R elaboration in mouse lemurs. Though it is true that both aye-aye and tarsier have more V1R copies than the diurnal primates compared here, they also show a high proportion of putative pseudogenes and an absence of some V1R subfamilies found in Cheirogaleidae (Figure 1A and B).

Our phylogenetic approach reveals a pattern of gene family evolution compatible with active gene birth and death (Nei et al. 1997; Csűrös and Miklos 2009; Hughes et al. 2018) with an independent V1R expansion isolated to Cheirogaleidae with three subfamily gains rather than a single more ancient expansion followed by losses in diurnal lineages (Table 1). Although the gain and loss dynamics of V1Rs over time is complex with uncertainty in the origins of specific subfamilies, variation in subfamily membership among species suggests that nocturnal primates possess more diverse repertoires than their diurnal counterparts (Figure 1A and B). These results are suggestive of an association between nocturnal life histories and V1R repertoire evolution, as well as the importance of chemosensation generally among nocturnal primates. Our findings are not conclusive, however, as the pattern observed in aye-aye deviates from this expectation, though it must be noted that the quality of the aye-aye genome assembly is considerably poorer than the others compared with the lowest contig/scaffold N50 and most incomplete BUSCO results (Supplementary Figure S1 and Table S1). An improved genome for aye-aye, a notably solitary primate (Sterling and Richard 1995), as well as genomes for species within the diurnal nest-dwelling genus, *Varecia*, will allow for more formal tests of how life history traits are correlated with V1R copy number.

### Subfamily membership and ancestral repertoire reconstruction

For each genome analyzed, we classified repertoire subfamily membership based on homology inferred from a maximum likelihood (ML) tree (Figure 4) and previously-described subfamily designations (Hohenbrink et al. 2012). During alignment, sequences that introduced excessive gaps to transmembrane regions were iteratively removed, resulting in alignments of increasing conservatism (see Materials and Methods “V1R repertoire estimation and ancestral count reconstruction”). We tested whether these varying alignments affected our estimates of subfamily composition and found little impact. Regardless of the number of sequences removed from the alignment, the relative proportions of subfamily membership within each species remained constant (Supplementary Figure S2). Although topological errors may contribute to uncertainty in gene count reconstructions, the ML tree shows 70% or greater bootstrap support for 63% of nodes (Figure 4), with little additional improvement possible due to the limitations of a single-exon gene family (Supplementary Table S3). Our results suggest that both the ancestral primate and the ancestral lemur had repertoires more limited in size and diversity than many living strepsirrhine primates, further supporting the controversial hypothesis that the ancestral primate was diurnal rather than nocturnal (Tan et al. 2005; Borges et al. 2018).

**Figure 4.**
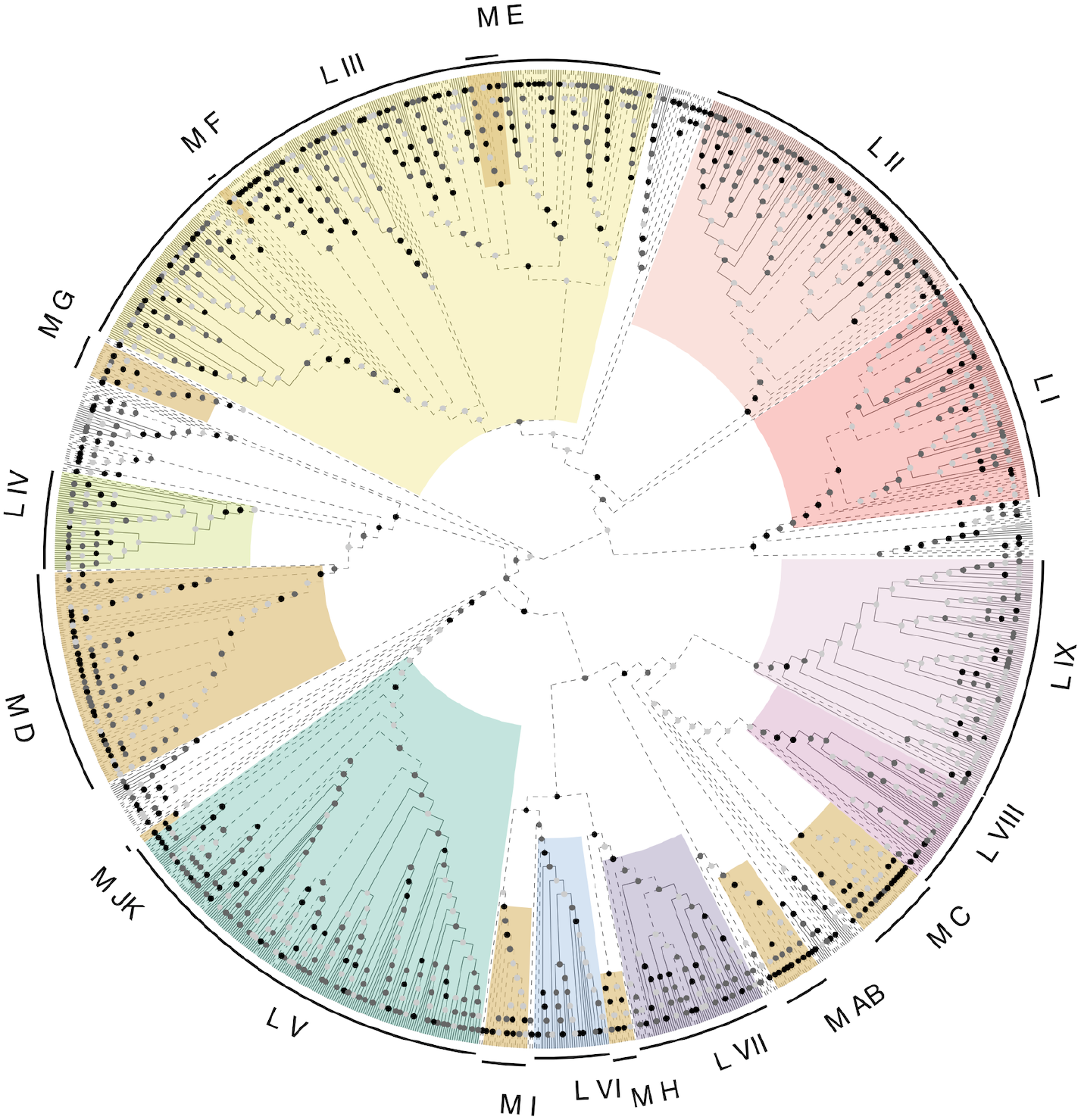
ML topology of V1R repertoire. V1R subfamilies in primates are highlighted and circumscribed based on Hohenbrink et al. (2012). There are nine described subfamilies in lemurs, L *Strep*/I through L IX, although not all lemur sequences fall into these subfamilies. Clades of V1R subfamilies with known function in mice are shown in burnt orange (M AB through M JK). Circles at nodes represent bootstrap support. Black nodes have 100% bootstrap support, dark grey nodes are supported with 70% or more bipartitions from bootstrap trees, and light grey nodes are weak or unsupported with less than 70% of bipartitions across bootstrap replicates. The topology is arbitrarily rooted for visualization. Solid lines represent dwarf and mouse lemur V1Rs (or branches subtending clades of dwarf and mouse lemur V1Rs). Dashed lines represent V1R lineages that are not within Cheirogaleidae.

Subfamily membership varies among the other extant strepsirrhines examined (Figure 1A and B). While *Otolemur garnetti* contains a very diverse repertoire, it lacks subfamily VII and IX membership. The diurnal lemurs lack receptors belonging to a few subfamilies, most consistently IV, VIII, and IX. The basal lineage within the lemuriform radiation, *Daubentonia madagascariensis*, lacks membership for most subfamilies, including *Strep*/I, II, IV, V, and VII. Subfamily I, referred to as “V1R*strep*” in Yoder et al. (2014), is used synonymously here for distinction from the mouse subfamily “I”. The repertoires of cheirogaleids are highly enriched for subfamily III, V, and IX membership, while the diurnal lemurs are enriched for subfamilies *Strep*/I, II, and III. In haplorrhine primates, repertoires contain only one or a few subfamilies. Ancestral state reconstruction with asymmetric parsimony (Csűrös and Miklos 2009; Csűrös 2010) revealed that the stem primate possessed only a subset of now extant V1R subfamilies, *Strep*/I, II, III, IV and VIII (Figure 1B). Subfamily IX has undergone a notable expansion in Cheirogaleidae, but the aye-aye repertoire also contains members from subfamily IX thus, subfamily IX is the only subfamily exclusive to nocturnal strepsirrhines, despite its absence in *Otolemur garnetti* (Figure 1A and B).

### Copy number variation in intraspecific *Microcebus murinus* repertoires

We resequenced eight *M. murinus* individuals of known pedigree from the colony at the Duke Lemur Center in Durham, North Carolina. Using these genomes, we estimated intraspecific variation in V1R repertoire size (Figure 2). For the eight resequenced *M. murinus* individuals, we observed low levels of intraspecific V1R repertoire size variation relative to size variation between taxonomic families with individual repertoires ranging from 86 to 105 intact V1R loci. Though one might expect that levels of intraspecific variation in V1R repertoire size in a captive population may be reduced relative to wild populations of *M. murinus*, the colony at the Duke Lemur Center shows signs of admixture from two distinct evolutionary lineages, *M. murinus* and *M*. *ganzhorni* (Larsen et al., 2017), presently recognized as distinct species (Hotaling et al., 2016). Therefore, the intraspecific variation observed here may actually be exaggerated, rather than reduced, which increases our confidence in the robustness of repertoire size estimates among species through sampling of single individuals. To test for the potentially confounding effects of sequencing and assembly error, one individual, DLC7033, was sequenced twice as a technical replicate. The duplicate genome assemblies respectively contained 92 or 96 intact loci indicating that sequencing and assembly error likely play a measurable role in generating variation among observed repertoire counts, though the effect appears to be modest. Thus, taking the results of the pedigree analysis as largely accurate, this emphasizes the highly dynamic nature of V1R repertoire size evolution, even over generational timescales.

### Complex history of diversifying positive selection in the dwarf and mouse lemurs

Our results agree with previous studies in finding that selection has acted pervasively across the V1R gene family over time (Hohenbrink et. al. 2012). Pervasive positive selection was revealed for all subfamilies identified in this study, even when analyzing the genus *Microcebus* alone (Supplementary Tables S4 and S5) and additional genome sequences for dwarf and mouse lemurs have likely increased the power of the sites tests. For example, positive selection was not evident for subfamily VII in a previous study limited to only *Microcebus murinus* (Hohenbrink et al. 2012). Furthermore, some subfamilies have unique profiles of sites under selection (Figure 5). Although lineage-specific rate variation is a confounding factor in V1R gene family evolution (Yoder et al. 2014), our analyses, spanning a range of taxon sampling schemes, show that our ability to characterize the V1R selection profiles are robust to such rate variation (Figure 5).

**Figure 5.**
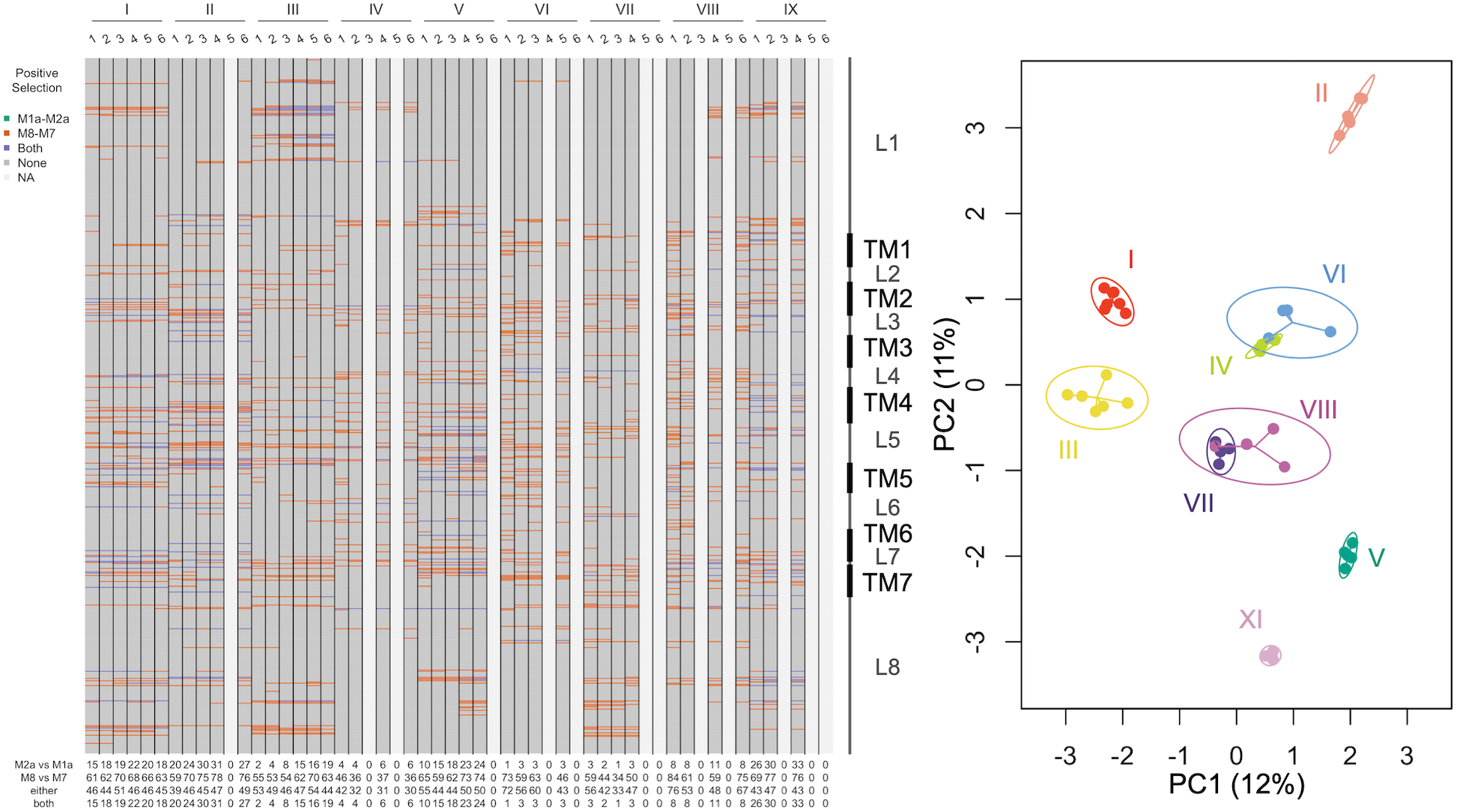
Sites under selection across the V1R alignment. Subfamilies are given at the top along with numbers that correspond to taxonomic filters. The first aligned codon starts at the top and the aligned codon position 588 at the bottom, with boundaries of transmembrane domains to the right. Sites under selection are colored. Missing columns means that the filter was redundant. Numbers along the bottom are counts of sites under selection detected by both model comparisons and their overlap. The boundaries of loop domains (L) and transmembrane domains (T) are shown along the aligned V1R repertoire. A PCA of sites under selection treated all codons as a binary character, determined by whether the site was under selection or not. Circle are 95% CIs for centroids of subfamily variation by taxonomic filters.

We performed two different model comparisons to differentiate between hypotheses of neutrality versus selection, and for the latter, for differentiating between the effects of rate constancy versus heterogeneity among sites. The M7 and M8 model comparisons always recovered more sites under selection than the more conserved M1a and M2a comparisons, but individual sites under selection detected by Bayes empirical Bayes with M2a were subsets of those detected by M8. Both model comparisons use likelihood ratio tests (LRTs) to detect positive selection and assume *dN/dS* is constant across branches, but the M2a and M1a comparison (Zhang et al. 2005) uses three finite mixtures of *dN/dS* while the M8 and M7 comparison (Yang et al. 2000) accounts for heterogeneity in *dN/dS* among nearly neutral sites with a beta distribution. Tests of pervasive positive selection were also performed on data realigned by subfamily, and similar estimates of proportions of sites under positive selection suggested that our site models were not misled by alignment errors (Supplementary Tables S6 and S7). Most individual sites under positive selection are unique to different subfamilies (Supplementary Figure S3) and reflect biases in selective pressures across different loop and transmembrane domains (Supplemental Figure S4). However, some selection profiles were more differentiated than others, such as *Strep*/I, II, V, and IX (Figure 5). The divergent selection profiles among subfamilies leads us to interpret positive selection acting on V1R genes in primates to be largely diversifying. Differentiated selection profiles among subfamilies are explained by biases among transmembrane and loop domains (Supplementary Material; Supplementary Figure S5; Supplementary Tables S8-S10).

Previous studies have indicated that extracellular loops have been primary targets of positive selection in V1Rs (Hohenbrink et al. 2012), and our results agree with these findings. Positive selection acting on extracellular loops two and three from Hohenbrink et al. (2012), identified here simply as loops three and five respectively, is evident (Supplementary Figure S5). These specific domains are probable regions where V1Rs bind to semiochemicals (Hohenbrink et al. 2012). Our results also show the transmembrane domains themselves, whether directly or by linkage, have also been under variable levels of positive selection (Supplementary Material; Supplementary Table S8). We find limited evidence for an enriched number of sites under positive selection in transmembrane domains four and five, and a depletion in transmembrane domain three, which have been previously predicted to form the ligand binding pocket of V1Rs (Kobilka et al. 1988; Pilpel and Lancet 1999; Palczewski et al. 2000; Yoder et al. 2014). These results prompt us to hypothesize that pervasive diversifying positive selection has accompanied selection for divergent function among V1R subfamilies, although additional evidence is needed for hypothesis testing.

Branch-site models detected evidence of episodic positive selection in the evolution of all V1R subfamilies except for lemur VIII (Supplementary Figure S6; Supplementary Table S11). Tests of episodic positive selection across the V1R subfamilies in the house mouse have been of little interest (Karn et al. 2010) and our tests of episodic selection here are generally not associated with the expansion of V1Rs in dwarf and mouse lemurs. However, subfamily IX would be a candidate for further investigation, given that it was the only subfamily to show notable levels of episodic positive selection, and is the only subfamily specific to nocturnal strepsirrhines. Further, many of the sites identified to be under positive selection correspond to the previously identified ligand binding domains (Supplementary Figure S6, Supplementary Table S12). Exploration of alternative topologies revealed that branches showing episodic positive selection were likely not due to topological errors (Supplementary Material; Supplementary Figure S6).

### Comparative evolution of V1R repertoires and genome architecture across Mammalia

Here we take advantage of the recently published chromosome-level assembly for *M. murinus* and other chromosome-level mammalian assemblies in an effort to identify genomic features that are generally associated with V1R expansion. The molecular environment of V1Rs is predicted to play a role in their regulation and has previously been studied only in mouse, rat, and pig (Lane et al. 2002; Stewart and Lane 2007; Kambere and Lane 2009; Michaloski et al. 2011; Dinka and Le 2017). We compared the expanded V1R repertoires of mouse and mouse lemur with the putatively contracted V1R repertoires of horse, cow, dog, and human. As predicted from previous studies (Kambere and Lane 2007; Kambere and Lane 2009), enrichment for repetitive LINE elements is associated with expansion of V1Rs in mammals (Supplementary Figure S7). We find that mouse lemur V1Rs primarily cluster by subfamily at chromosomal locations across the genome as is also characteristic of the V1R repertoire in mouse. Only mouse lemur subfamily VIII does not form a cluster but is instead uniquely dispersed across three different chromosomes (Figure 6). We also analyzed the locations of all regions demonstrating V1R homology to determine if there are any potential pseudogenized subfamilies or clusters in the genome and found no evidence for pseudogenized clusters of V1Rs in mouse lemur (Supplemental Figure S12). Both LINE enrichment and physical clustering of V1R loci have been predicted to be associated with proper regulation of V1Rs (Lane et al. 2002; Kambere and Lane 2007) and may be characteristic of expanded V1R repertoires in general.

**Figure 6.**
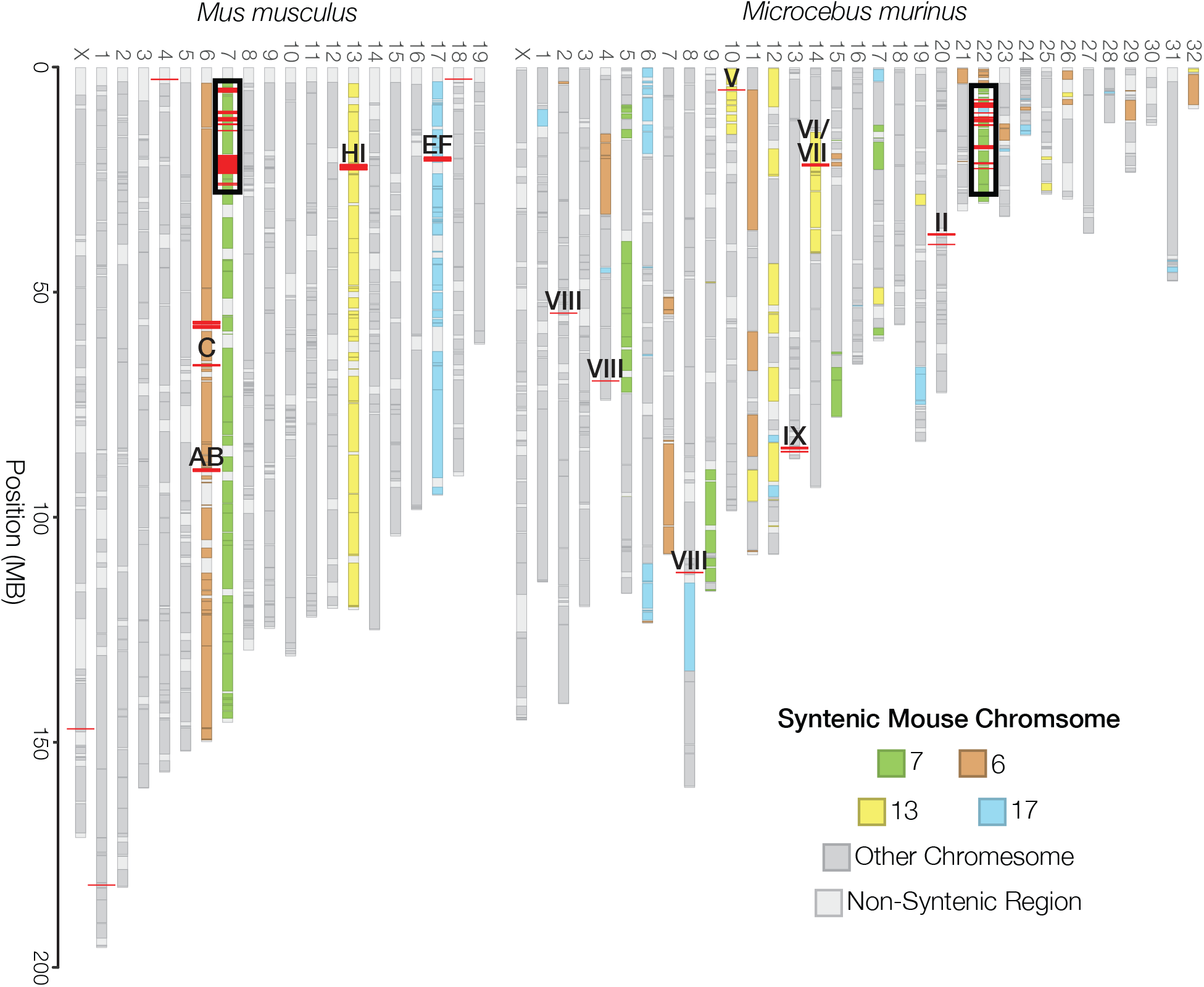
Chromosomal synteny between mouse and mouse lemur V1R-containing regions. Synteny between *Mus musculus* and *Microcebus murinus* was estimated using the SynChro software. Chromosomes are colored relative to V1R-containing mouse chromosomes. V1R loci are indicated with red lines and are labelled by subfamily identity. Regions outlined in black are enriched for V1R loci and are examined in further detail in Figure 7B.

To investigate whether homologous subfamilies have retained chromosomal synteny in species with expanded repertoires and across mammals broadly, we evaluated chromosomal synteny for each species relative to mouse using the SynChro software (Drillon et al. 2014; Figure 6, Figure 7A and B, Supplementary Figures S8-S11). In mouse and mouse lemur, most homologous V1R subfamilies retain chromosomal synteny (Figure 6, Figure 7A and B). Mouse subfamily D is most closely related to mouse lemur subfamily IV, and both subfamilies share mouse chromosome 7 synteny. Subfamilies J/K and V as well as subfamilies G and *Strep*/I also share mouse chromosome 7 synteny. Lemur subfamily III is syntenic with mouse E and F on mouse chromosomes 6 and 7. Lemur subfamilies VI and VII are syntenic with mouse subfamilies H and I on chromosome 13. Lemur subfamilies not sharing synteny with any mouse subfamily include subfamilies II, VIII, and IX. The expanded subfamilies in Cheirogaleidae, IV, VII, and IX, all map to different chromosomal regions of the *M. murinus* genome and were not linked on an ancestral syntenic block based on comparisons between *M. murinus* and mouse. Therefore subfamily expansions have occurred independently and not as tandem duplications of a single genomic region.

**Figure 7.**
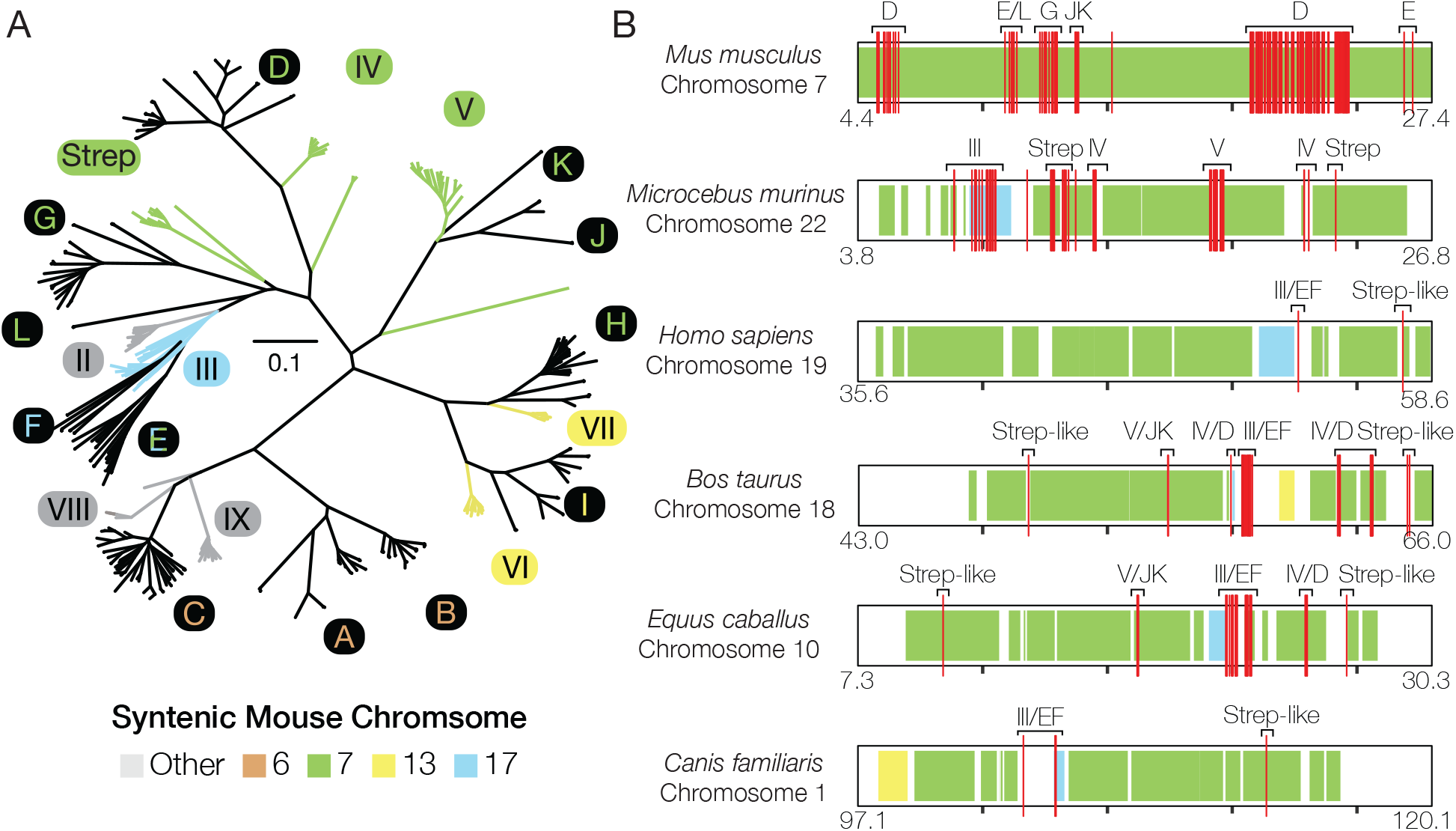
Highly orthologous loci on “hotspot” V1R chromosome. (A) RAxML tree of *Mus musculus* V1R cDNA sequences with intact V1R sequences from the *Microcebus murinus* genome. Mouse subfamilies are encircled in black and labelled by chromosomal location. Mouse lemur subfamilies are labelled in black and encircled in the color corresponding to the syntenic mouse chromosomal location. (B) Chromosomal “hotspot” regions enriched in V1R loci from several mammalian taxa. Orthologous regions are shaded by syntenic mouse chromosome. V1Rs loci are labelled by phylogenetic relationship to mouse/lemur subfamilies. Starting and end genomic positions are given for each species, and all regions are 23Mb long with tick marks representing 5 Mb intervals.

Interestingly, when comparing all mammalian species examined, our results reveal that in each species, one chromosome contains a very dense block of highly homologous subfamilies on a backbone of mouse chromosome 7 synteny, referred to here as “V1R hotspots” (Figure 7B). These hotspots usually contain receptors of the EF/III, D/IV, JK/V, and Strep/G subfamilies, and cluster order is maintained with a few species-specific subfamily deletions. The chromosomal synteny of the “hotspots” is rarely interrupted, and if interrupted, it is almost exclusively interrupted by a stretch of synteny from another mouse chromosome containing V1Rs. These interleaving regions in hotspots are usually chromosome 13 or 17, indicating that genomic regions where V1Rs cluster are also subject to increased gene duplication rates. Interestingly, the only putative intact members of the contracted human V1R repertoire are also contained within this “hotspot” location and share homology with hotspot subfamilies.

Previous studies of Laurasiatheria have predicted that the V1R repertoires of cow, horse, and dog consist mostly of highly orthologous loci with evolutionary conserved functions (Yohe et al. 2018). While conserved function remains to be shown experimentally, retained synteny of these Laurasiatherian V1Rs within hotspots across Mammalia supports the hypothesized ancient origin of these subfamilies and reinforces the idea that V1Rs in these subfamilies are orthologous in function (Ohara et al. 2009; Yohe et al. 2018). Mouse lemur V1Rs show striking structural similarities to the functionally diverse repertoire of mouse and considering the independent gains in copy number and novel subfamily evolution, coupled with variable rates of molecular evolution and selective pressures, V1Rs in mouse lemurs may serve as an ideal system for elucidating pheromone evolution in primates. Similar patterns of deep synteny have been described for ∼80 My of odorant receptor evolution in bees (Brand and Ramirez 2017). Considered in this context, our results suggest that chemosensory gene family evolution may follow similar molecular “rules” in organisms with vastly different natural histories, even when evolved independently from different ancestral gene families, as would be the case comparing mammals to insects.

## Conclusions

We revealed that an expansion of the V1R gene family is shared across the dwarf and mouse lemurs, and that duplicate V1R gene copies have been evolving under strong selective pressures. Divergent patterns of molecular evolution among V1R subfamilies and diversity in subfamily membership and abundance suggests that V1Rs may serve as a test case for studying the evolution of sensory drive in primates. Pheromone detection among nocturnal primates, especially the morphologically cryptic mouse lemurs, may be more important for driving and maintaining species boundaries than previously appreciated. Syntenic analyses with improved genomic resources revealed strikingly similar genetic architecture between the expanded V1R repertoires of mouse and mouse lemur, and that some V1R subfamilies have been maintained in V1R “hotspots” across ∼184 million years of mammalian evolution (dos Reis Mario et al. 2012). Characterizing additional features of V1R hotspots across species will be important for future studies translating experimental genetic studies in mice to primates such as mouse lemur.

## Materials and Methods

### Sampling and DNA extraction

To improve the resolution of the V1R repertoire expansion in lemurs, we sequenced the genomes of *Microcebus griseorufus*, *M. mittermeieri*, *M. ravelobensis, M. tavaratra*, and *Mizra zaza*. Tissue biopsies were taken from wild individuals in Madagascar from 1997-2015 and from captive individuals at the Duke Lemur Center (Supplemental Table S13). To investigate within species variation in V1R repertoires, we also resequenced eight individuals from the Duke Lemur Center *Microcebus murinus* colony. Blood and tissue samples were collected in 2016 in accordance with IACUC guidelines. For the novel strepsirrhine genomes, DNA was extracted following manufacturer instructions using the Qiagen DNeasy Blood and Tissue kit, while DNA from the *Microcebus murinus* resequenced individuals was extracted using the Qiagen MagAttract Kit (Qiagen, Germantown, MD, USA).

### Genome Sequencing and Assembly

The genomes of *Microcebus griseorufus*, *M. mittermeieri*, *M. tavaratra*, and *Mizra zaza* were sequenced at the Baylor College of Medicine as approximately 400bp insert libraries on a single lane of an Illumina HiSeq 3000 with paired-end 150bp reads. We sequenced the *Microcebus ravelobensis* genome from two libraries, one with an average insert size of 570bp on an Illumina HiSeq 2000 at Florida State University and the other with a 500bp insert library on 5.5% of both lanes of an Illumina NovaSeq at the Duke University GCB Sequencing Core. We also generated two additional cheirogaleid assemblies for *Cheirogaleus sibreei* and *C. medius* (Williams et al. 2019)*. Cheirogaleus sibreei* was sequenced from a 300bp insert library on the Illumina HiSeq 4000 at the Duke University GCB Sequencing Core with paired-end 150bp reads. A reference genome was generated and assembled for *Cheirogaleus medius* using Dovetail Genomics. All other genomes were assembled using MaSuRCA v3.2.2 (Zimin et al. 2013). We assumed an insert size standard deviation of 15% and used automatic kmer selection. However, we did not use MaSuRCA’s scaffolds for annotation and downstream analyses. Scaffolds were obtained from SSPACE (Boetzer et al. 2010), which also attempted to correct assembly errors and extend contigs from MaSuRCA. *De novo* assembly statistics are available in the supplementary material (Supplementary Table S1) as well as annotation details (Supplementary Table S14) and SRA identifiers (Supplementary Table S15).

The eight *Microcebus murinus* individuals were resequenced from high molecular weight DNA prepared using the 10X Genomics Chromium platform. Briefly, high-molecular weight molecules of DNA are partitioned into gel beads with unique barcodes then prepared for Illumina sequencing (Weisenfeld et al. 2017). The resulting short-read libraries are barcoded such that individual “linked reads” can be traced back to their molecule of origin assisting the genomic scaffolding process. The libraries were size selected to approximately 550bp and sequenced on the Illumina HiSeq 4000 system at the Duke University GCB Sequencing Core. We then used the 10X Genomics Supernova assembly software to *de novo* assemble the resequenced genomes (version 2.0.1, 10x Genomics, San Francisco, CA, USA). One replicate individual was sequenced twice, and genomes were assembled *de novo* from each individual library.

BUSCO version 3.0.2 and Assemblathon2 scripts were used to assess genome quality statistics (Supplementary Figure S1; Supplementary Table S16; Simão et al. 2015). Additional genomes analyzed in this study were downloaded from the NCBI genome database and include all available Strepsirrhine genomes and additional high-quality primate and mammalian genomes for phylogenetic coverage (Supplementary Table S16).

### V1R repertoire estimation and ancestral count reconstruction

To assess total V1R repertoires in each species, tblastn searches (e-value cut-off = 0.001) were conducted with the blast+ software suite (version ncbi-blast-2.6.0+; Altschul et al. 1990) using available mouse and mouse lemur V1R query protein sequences downloaded from NCBI GenBank against the genomes analyzed in this study (Camacho et al. 2009). Duplicate protein sequences were removed from the query database using CD-HIT version 4.6 (Li and Godzik 2006). Bedtools merge (version 2.27.1) was used to merge overlapping hits within a genome, and bedtools slop and getFasta were used to extract receptor candidate regions longer than 600bp with 50 bp of upstream and downstream surrounding sequence (Quinlan and Hall 2010). For a full list of V1R containing regions analyzed see Supplementary File X).

To remove potential pseudogenes from further analyses, we used Geneious version 9.0.5 to predict open reading frames (ORFs) and considered sequences intact if they contained an ORF longer than 801bp. We then used MAFFT version 7.187 with the E-INS-i algorithm to align intact sequences from all species using the iterative approach described in Yoder 2014 (Katoh and Standley 2013; Yoder et al. 2014). The MAFFT algorithm is recommended for approaches analyzing ancestral sequence reconstruction (Vialle et al. 2018). A gene phylogeny was constructed using RAxML version 7.2.8 using the GTRGAMMAI nucleotide model with the rapid bootstrapping and search for best ML scoring tree algorithm with 500 bootstraps (Stamatakis 2014). We then assigned primate sequences to the subfamilies *Strep*/I-IX designated in Hohenbrink 2012 (Hohenbrink et al. 2012). The number of intact V1Rs, percentage of intact V1Rs, and the total V1R count were calculated for each species as well as subfamily membership. We then used Count version 10.04 with the Wagner parsimony algorithm and a gain penalty of 2 to infer total ancestral vomeronasal repertoire size and ancestral subfamily membership (Csűrös and Miklos 2009; Csűrös 2010).

### Establishing synteny of V1Rs across mammalian species

Genomes with chromosome level scaffolding information (*Mus musculus, Microcebus murinus, Homo sapiens, Equus caballus, Bos taurus*, and *Canis familiaris*) were used to assess chromosomal synteny of vomeronasal subfamilies among mammalian species. SynChro (Drillon et al. 2014) version SynChro_osx (January 2015) was used to reconstruct synteny blocks between each genome with *Mus musculus* as reference with a delta parameter of 2 using GenBank annotation files from Ensembl release 93 (Figure 6; Supplemental Figures S8-S11; Drillon et al. 2014). Orthologous block information was compared with vomeronasal receptor location for each species (Figure 7A and B).

### Detecting evidence of positive selection

Evidence for positive selection in V1R repertoires was evaluated with PAML 4.8e (Yang 2007). We used two different tests to detect both individual sites under pervasive positive selection throughout the tree (sites models) and individual branches that show an episodic burst of positive selection (branch-site models). For sites models, we applied two tests to each of the nine subfamily trees and alignments: 1) Comparison of the null hypothesis that all sites are a mixture of purifying and neutral rates of molecular evolution (M1a) and the alternative that allows for a third class of sites under positive selection (M2a; Zhang et al. 2005). 2) A null hypothesis that allows for a mixture of ten discretized beta-distributed site classes (M7), while the alternative hypothesis allows an extra component under positive selection (M8;Yang et al. 2000). Each of the recognized lemur subfamilies were analyzed separately. The ggtree R package (Yu et al. 2017) was used to extract subtrees for each subfamily and alignments were parsed with Perl scripts. Because signatures of positive selection may be time-dependent (Peterson and Masel 2009; Pegueroles et al. 2013), we explored variation in sites under positive selection using six different taxonomic filters: 1) *Microcebus*, 2) Cheirogaleidae, 3) Lemuriformes, 4) Strepsirrhini, 5) Primates, and 6) Euarchontoglires. For each site test, we assumed the LRT was 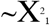 and individual sites were detected using the Bayes empirical Bayes procedure where the posterior probability of selection for each site was determined using the MLE *dN/dS* for the positive selection rate class (Yang et al. 2005). Individual sites were considered to have sufficient evidence for positive selection if the posterior probability was greater than 0.95. Enrichment of sites under selection in transmembrane domains used simple chi-square tests and Fisher exact tests in R (R Core Team 2018) for individual transmembrane and loop domains. Transmembrane domains were predicted using *M. murinus* sequences from subfamilies *Strep/*I through IX with TMHMM (Krogh et al. 2001) through the TMHMM server (http://www.cbs.dtu.dk/services/TMHMM/; last accessed 29 January 2019). Since V1R genes are expected to have seven transmembrane domains (Dulac and Axel 1995), only predicted structures with seven transmembrane domains were used to determine transmembrane domain boundaries in our alignment of the entire V1R repertoire. Predictions that had fewer or more than seven transmembrane domains are assumed to be due to inaccuracies of TMHMM (Melén et al. 2003) and not real domain losses or gains.

Of important note, the entire V1R repertoire was prohibitively large for ML optimization over the entire tree; we applied tests for selection to individual subfamilies to circumvent this limitation. This strategy also provided a way to evaluate contributions of alignment and topological errors to evidence of positive selection. First, we evaluated if the ML topology estimated from the entire repertoire was a plausible hypothesis using AU tests (Shimodaira 2004). First, we estimated the ML topology and branch lengths for each subfamily using the parsed alignments (i.e. the data was not re-aligned) using the same RAxML model and search strategy as the first analysis. We then re-aligned translated amino acid data with MAFFT and estimated phylogeny once more. Site log-likelihoods were then optimized for the three topologies with RAxML and AU p-values computed with CONSEL using the default multiscale bootstrapping strategy (Shimodaira and Hasegawa 2001). Bootstrap trees were also collected for the re-aligned data, but bipartitions were drawn onto the topologies parsed from the entire V1R repertoire tree. The ratio of bootstrap support values was used to identify potential topological errors; bipartitions in the original topology that are absent when the sequences for each subfamily were re-aligned. Site tests were run for both the parsed and re-aligned data to check for consistency in the inference of sites under positive selection across alignments.

Branch-site tests (Zhang et al. 2005) were performed for each branch for each subfamily, except subfamily III, which was still computationally limiting. Each test assumed the LRT was 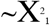, but we applied the Benjamini-Hochberg multiple testing correction (Benjamini and Hochberg 1995). With this correction, we do expect some false positives, but the family-wise error rate should be below 5% (Anisimova and Yang 2007) while not underpowering tests towards the tips of the trees (dos Reis and Yang 2011). Tests were only performed on the parsed topology without removing any species, but branches with evidence of episodic positive selection and the bootstrap ratios with re-aligned data were mapped to nodes of the subtree topologies using ggtree to help identify cases where topological errors might lead to false signatures of positive selection (Mendes and Hahn 2016). Individual sites with evidence of episodic positive selection were evaluated using the Bayes empirical Bayes procedure (Yang et al. 2005).

### Data Access

Newly sequenced genome data will be made available through NCBI upon publication. Complete record information is given in Supplementary Material (Supplementary Table S15).

## Acknowledgements

We thank the Malagasy authorities for permission to conduct this research and Duke Lemur Center staff, especially Erin Ehmke, Bobby Schopler, and Cathy Williams, for providing the *Microcebus murinus* and *Mirza zaza* tissue samples. We are grateful to our colleagues at Baylor College of Medicine, Jeff Rogers and Kim Worley, for many insightful discussions of mouse lemur genomics. Phillip Brand and Jeff Thorne provided critical review of the manuscript leading to its significant improvement. We thank Simon Gregory’s lab for preparing the 10x Genomics libraries. We are grateful for the support of Duke Research Computing and the Duke Data Commons (NIH 1S10OD018164-01) and appreciate the donation of free sequencing for *Microcebus ravelobensis* provided by the Duke GCB Sequencing Core. ADY gratefully acknowledges support from the John Simon Guggenheim Foundation and the Alexander von Humboldt Foundation during the writing phase of this project. The study was funded by a National Science Foundation Grant DEB-1354610 to ADY and DWW and Duke University startup funds to ADY. This is Duke Lemur Center publication no. XXX.

