## Supplemental Information for "Elaborate expansion of syntenic V1R hotspots correlates with high species diversity in nocturnal mouse and dwarf lemurs"

**Affiliations**:

[8]Current address: Department of Biological Sciences, University of Denver, Denver, CO 80208

†Equal contributors

### Supplementary Methods

#### Genome Annotation

Although *de novo* gene annotations were not used in this study, gene annotations are provided with genomes available through NCBI. RNAseq data (Peng et al. 2014) was used to train gene prediction models with the grey mouse lemur genome. RNAseq libraries from multiple tissues and replicates (Table S15) were used to assemble a single transcriptome with TRINITY v2.2.0 (Haas et al. 2013). Assembled transcripts and homologous protein evidence from closely-related species were used as direct evidence for gene annotations as well as for training the *ab initio* gene predictor SNAP (Korf 2004) with MAKER v2.31.9 (Cantarel et al. 2008) for the *Microcebus murinus* 3.0 assembly (https://www.ncbi.nlm.nih.gov/genome/?term=Microcebus+murinus; last accessed 8 November 2017). For an independently trained gene predictor, we optimized the AUGUSTUS v3.3 (Stanke et al. 2006) human parameters with the *M. murinus* 3.0 assembly using BUSCO v3.0.2 (Simão et al. 2015). Final annotations for all five genomes were estimated with a single MAKER run using a combination of transcriptome and homologous protein alignment as well as gene predictions. Genomes of related species used for homologous protein evidence are as follow:

- Norway Rat 6.0 (https://www.ncbi.nlm.nih.gov/genome/?term=rattus; last accessed 8 November 2017)
- House Mouse GRCm38.6 (https://www.ncbi.nlm.nih.gov/genome/?term=Mus+musculus; last accessed 8 November 2017)
- Chimpanzee 3.0 (https://www.ncbi.nlm.nih.gov/genome/?term=Pan+troglodytes; last accessed 8 November 2017)
- Gorilla 4.0 (https://www.ncbi.nlm.nih.gov/genome/?term=Gorilla; last accessed 8 November 2017)
- Sumatran orangutan 2.0.2 (https://www.ncbi.nlm.nih.gov/genome/?term=Pongo+abelii; last accessed 8 November 2017)
- Human GRCh38.p11 (https://www.ncbi.nlm.nih.gov/genome/?term=Homo+sapiens; last accessed 8 November 2017)

The transcriptome was assembled with 15 paired-end 100bp Illumina libraries representing 9 tissues. Reads were trimmed with trimmomatic. A single transcriptome was then assembled by pooling all libraries, using *in silico* normalization of each library to avoid unnecessarily large RAM requirements. We then estimated the fragments per kilobase million (FPKM) per locus using RSEM (Li and Dewey 2011) as called from TRINITY.

For training our *in silico* gene predictors, we first trained SNAP (Korf 2004) from within MAKER. We first ran MAKER on the *Microcebus murinus* 3.0 assembly only using evidence-based annotations from our *de novo* transcriptome assembly and homologous proteins of related species. A single GFF file was merged from the resulting annotations and MAKER was run again only using SNAP with hints supplied from the previous GFF. A new GFF file was made from SNAP's predications and we repeated gene finding once more to refine the HMM used by SNAP.

**Enrichment of LINES in V1R clusters**

We assessed the molecular environment of vomeronasal receptors with the newly improved *Microcebus murinus* genome compared to mouse (Larsen et al. 2017). We used bedtools closest to determine average distance separating V1Rs. We assigned receptors within 500kb of other receptors to clusters then calculated repeat element density and analyzed surrounding annotations for the V1R-containing subset of the genome. We generated 100 randomly distributed subsets of each genome of similar size and base pair composition to the V1R-containing subsets for each species. We estimated repeat element density in the random and V1R-containing subsets using RepeatMasker version open-4.0.6 with the RepBase database (Chen 2004; Jurka et al. 2005).

#### Supplementary Results

##### Subfamily-Specific Patterns in Sites Under Pervasive Positive Selection

Sites under positive selection were biased towards transmembrane domains in subfamilies *Strep*/I, II, V, VI, and VIII, although not consistently across taxonomic filters (Supplementary Table S8). Increased numbers of sites under selection in transmembrane domains were only evident when analyzing Lemuriformes, Cheirogaleidae, or *Microcebus*, which suggests that selection on these structurally conserved domains may be occurring at more shallow time scales or specifically in dwarf and mouse lemurs. Subfamily III was enriched for sites under positive selection in loop domains (Supplementary Table S8). Some individual transmembrane domains showed decreased or increased levels of positive selection depending on the subfamily (Supplementary Figure S5; Table S9). For instance, transmembrane domain three was conserved, showing less evidence of positive selection with respect to all other transmembrane domains, across all taxonomic filters in subfamilies *Strep*/I and IX (Figure 5; Figure S5). Some loop domains appeared to experience more positive selection in specific subfamilies too; loop three was particularly enriched for sites under positive selection in subfamilies II and VII; loop five in subfamilies II, IV, V, and VI while loop seven was enriched for positive selection in subfamily IX.

##### Lemur Subfamily IX and Uncertainty in Detecting Episodic Positive Selection

Approximately 10% of sites are estimated to be under selection along the basal cheirogaleid branch and were identified in transmembrane domain three as well as in loops three and five (Supplementary Table S12). However, signatures of episodic positive selection in subfamily IX are typically downstream or upstream of domains comprising the ligand-binding pocket (Supplementary Table S12). Otherwise episodic positive selection was detected in 5% or less of sites among branches with many sites mapping downstream of transmembrane domain seven where alignment can be less reliable. However, incorrectly identified sites under positive selection due to alignment or topological errors were likely minimal. For each lemur subfamily, the topological relationships based on the ML analysis of the entire repertoire could not be rejected when compared to arguably more correct trees based on realigned data (Supplementary Table S3). Bootstrap trees from the entire repertoire and realigned subtree analyses found only one false branch, which occurred in subfamily IX (Supplementary Figure S6).

#### Supplementary Figures

**Figure S1 – Comparing Benchmarking Universal Single-Copy Orthologs (BUSCO) results across publicly available Strepsirrhine genomes.** Bars depict percentage of orthologs recovered from assemblies that were complete (Single), more than once but were still complete (Duplicate), recovered but were fragmented (Fragment), or that were not recovered from the genome (Missing). Orthologs were from the mammalian BUSCO gene set.

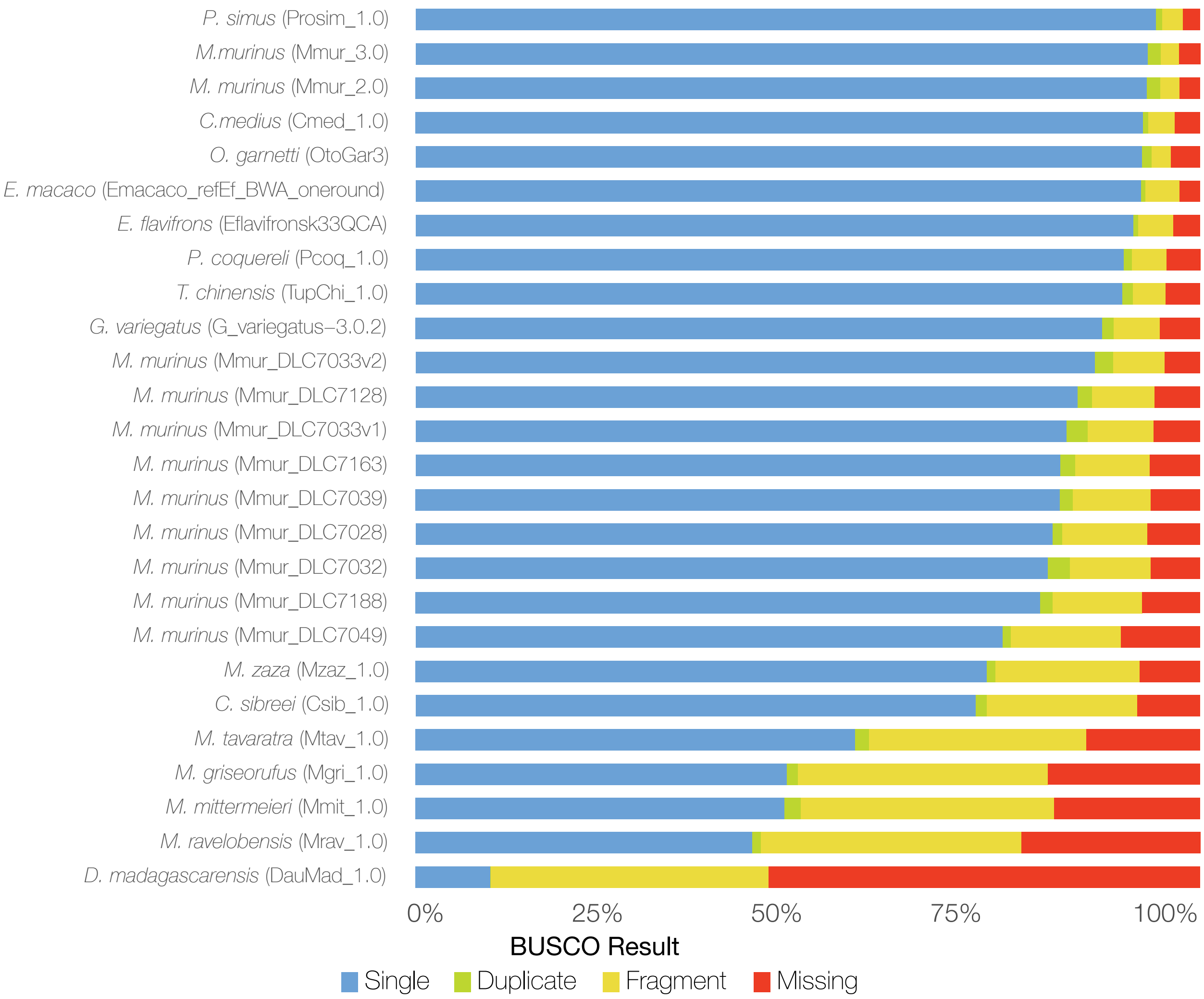

**Figure S2 – Changes in subfamily counts given increasing alignment stringency.** The number of gene copies per subfamily per species is largely insensitive to removal of sequences introducing that require multiple gap openings in transmembrane domains. Alignment version 1 (v1) represents the least conservative alignment while v17 represents the most conservative alignment.

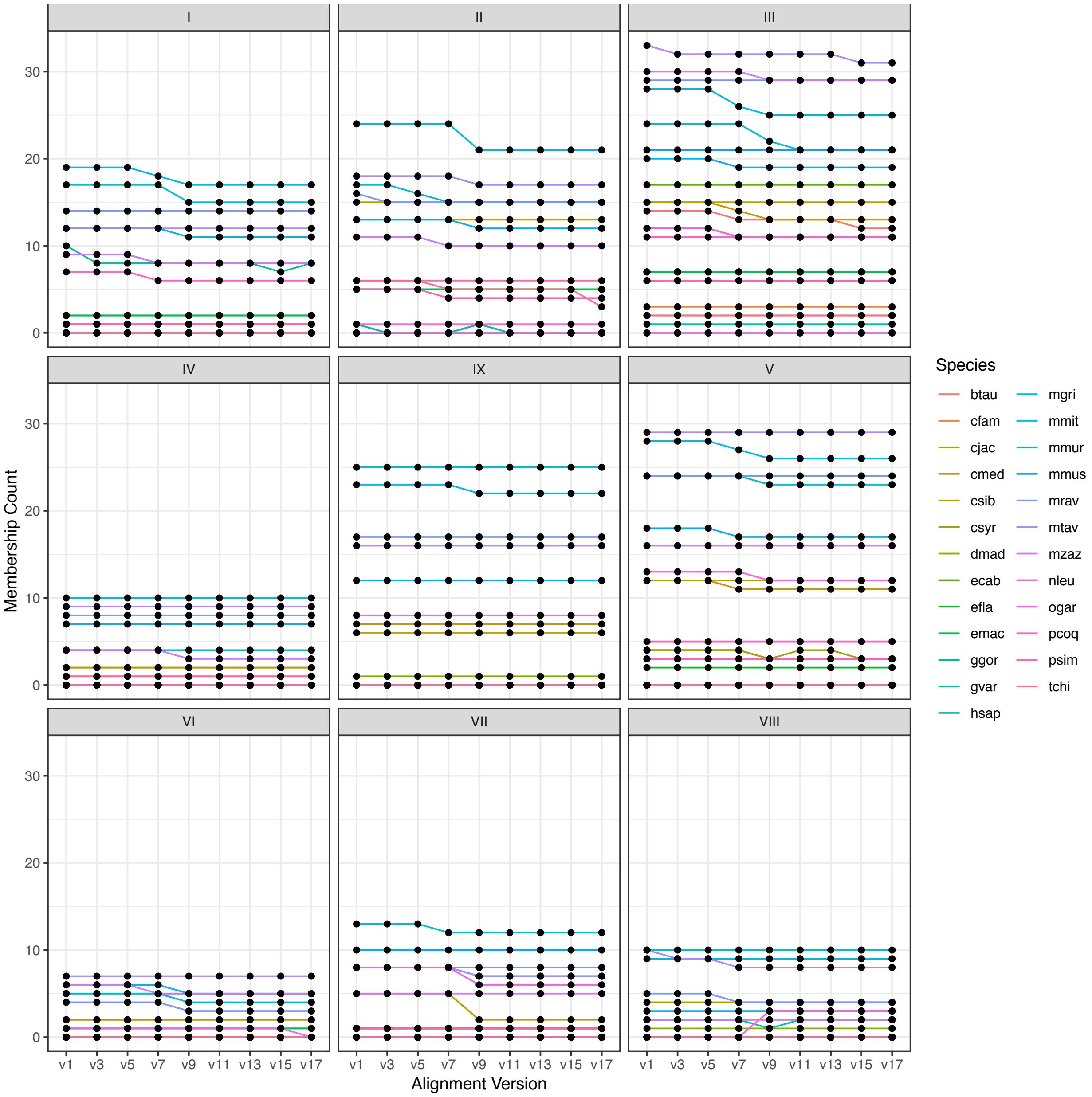

**Figure S3 – Sites under pervasive positive selection shared across subfamilies.** Although many sites under positive selection are unique to one subfamily, most sites are under selection in two or more subfamilies. Shared sites may be found in at least one taxonomic filter.

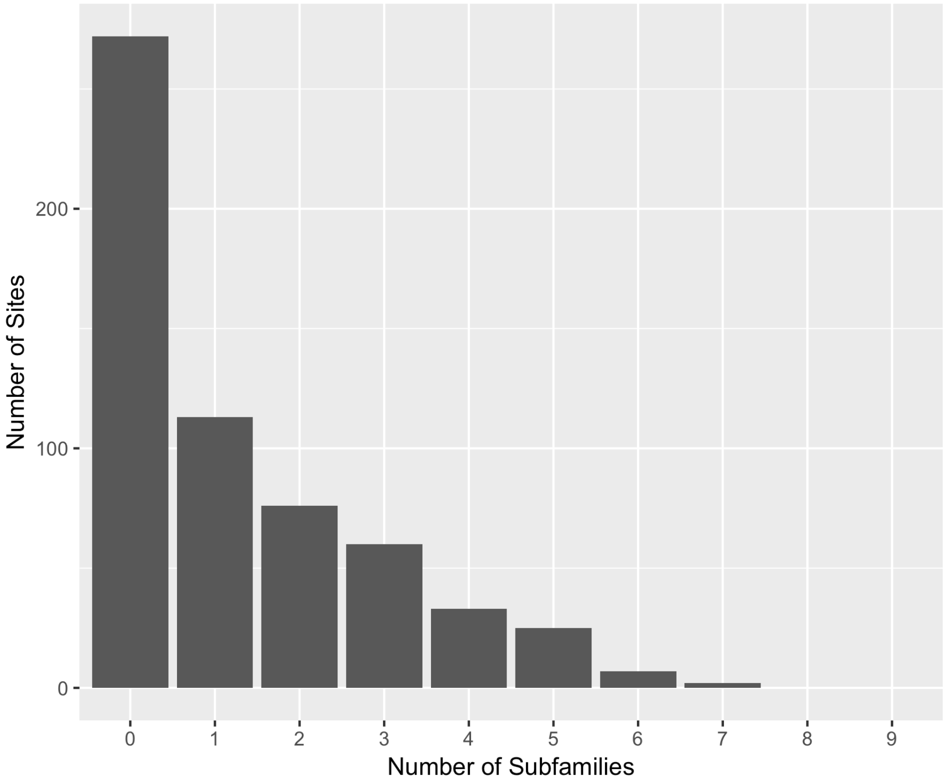

**Figure S4 – Transmembrane domain predictions in *Microcebus murinus* sequences from subfamilies I-IX.** Each line is a an aligned and putatively intact V1R sequence from the *M. murinus* genome. Pink lines are extracellular loops, blue lines are inside, and thickened gray lines represent predicted transmembrane domains. Black vertical dashed lines are the average start and stop positions for transmembrane domains one through seven when mapped to the codon alignment. Only models with seven predicted transmembrane domains were used for obtaining alignment-wide start and stop sites.

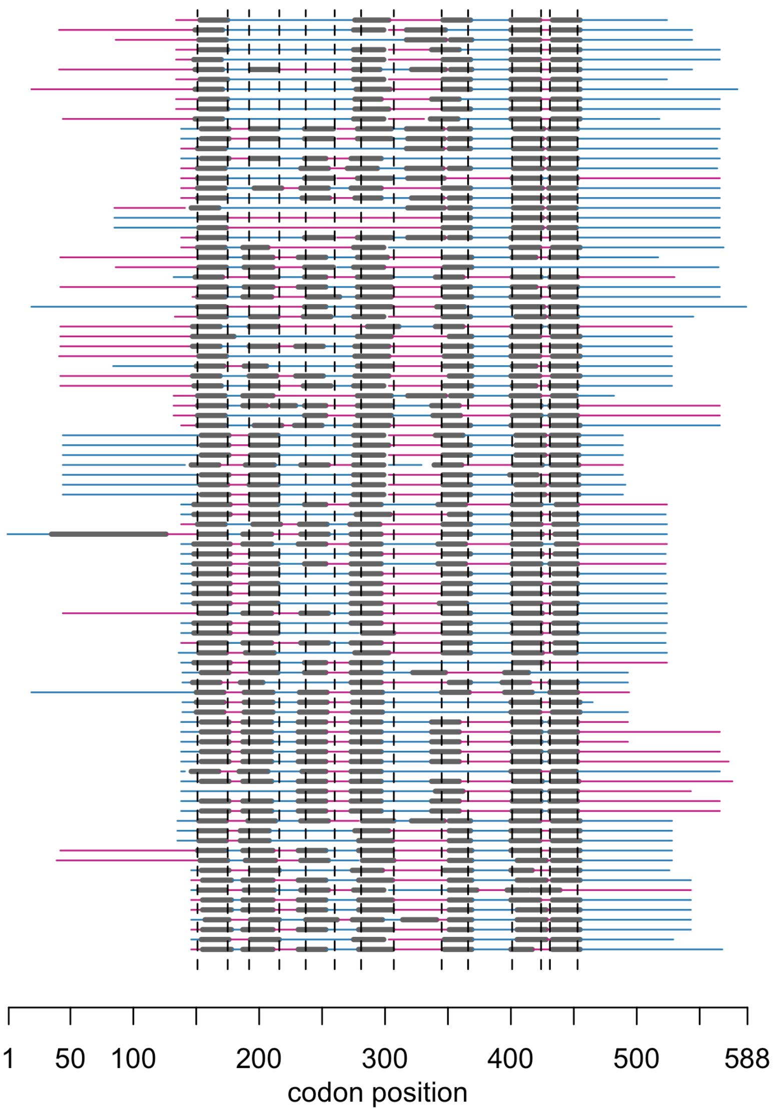

**Figure S5 – Over- or under-representation of sites under selection among V1R domains.** Red boxes represent more than expected and blue boxes represent fewer than expected sites under selection for loop (L) and transmembrane (T) domains. Subfamilies are given on the y-axis and multiple rows represent different taxonomic filters.

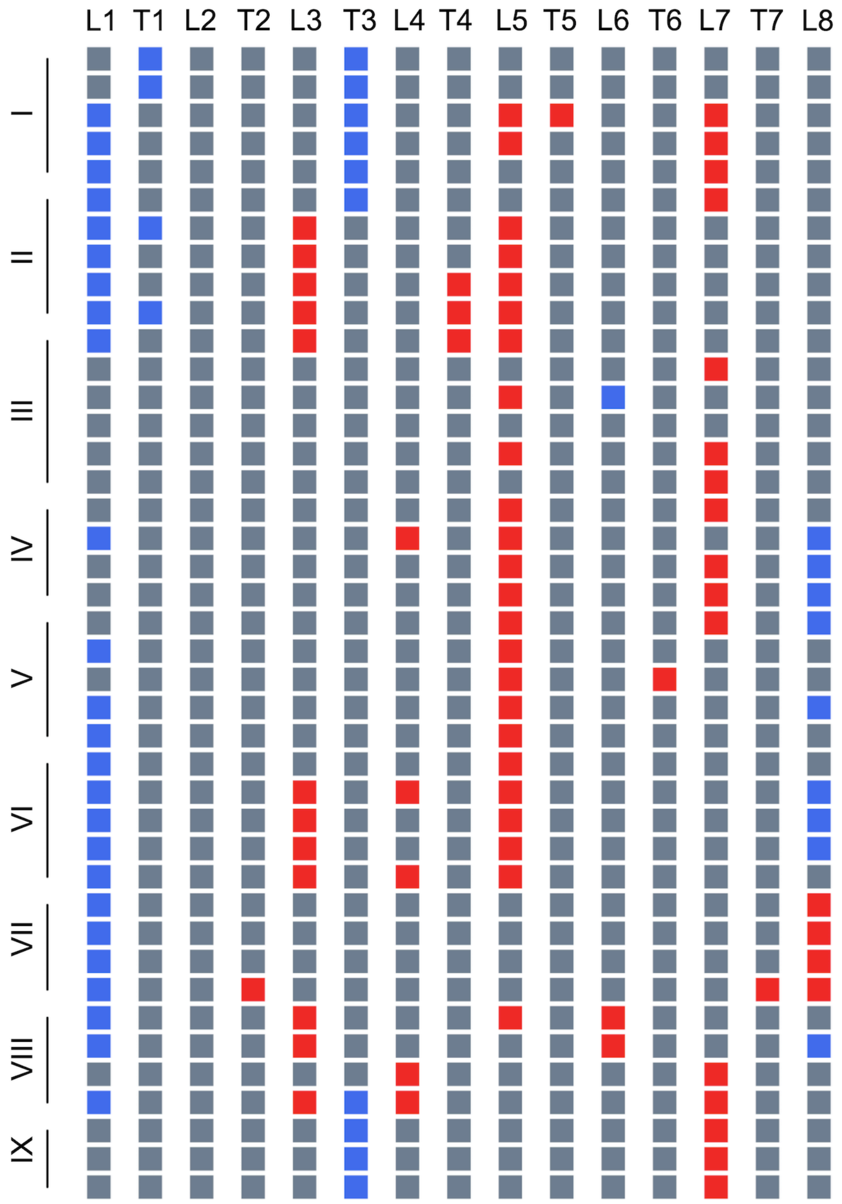

**Figure S6 – Branches with evidence of episodic positive selection across subfamilies.** The subtrees are parsed from the ML topology of the entire V1R repertoire. Blue diamonds at nodes represent evidence of episodic positive selection occurring during the subtending branch. Solid lines are taxa within Cheirogaleidae while dashed lines may represent other primates or *Tupaia*. Circles at internal nodes represent the ratio of bootstrap support between the realigned data and alignment from the entire repertoire. Red circles indicate bipartitions supported in the whole repertoire ML analysis that were not supported in the realigned ML trees. No episodic positive selection was detected in subfamily VIII; subfamily III was too large for computationally efficient estimation of likelihoods. Node numbers correspond to Supplementary Table S12 for identifying individual sites under selection along branches.

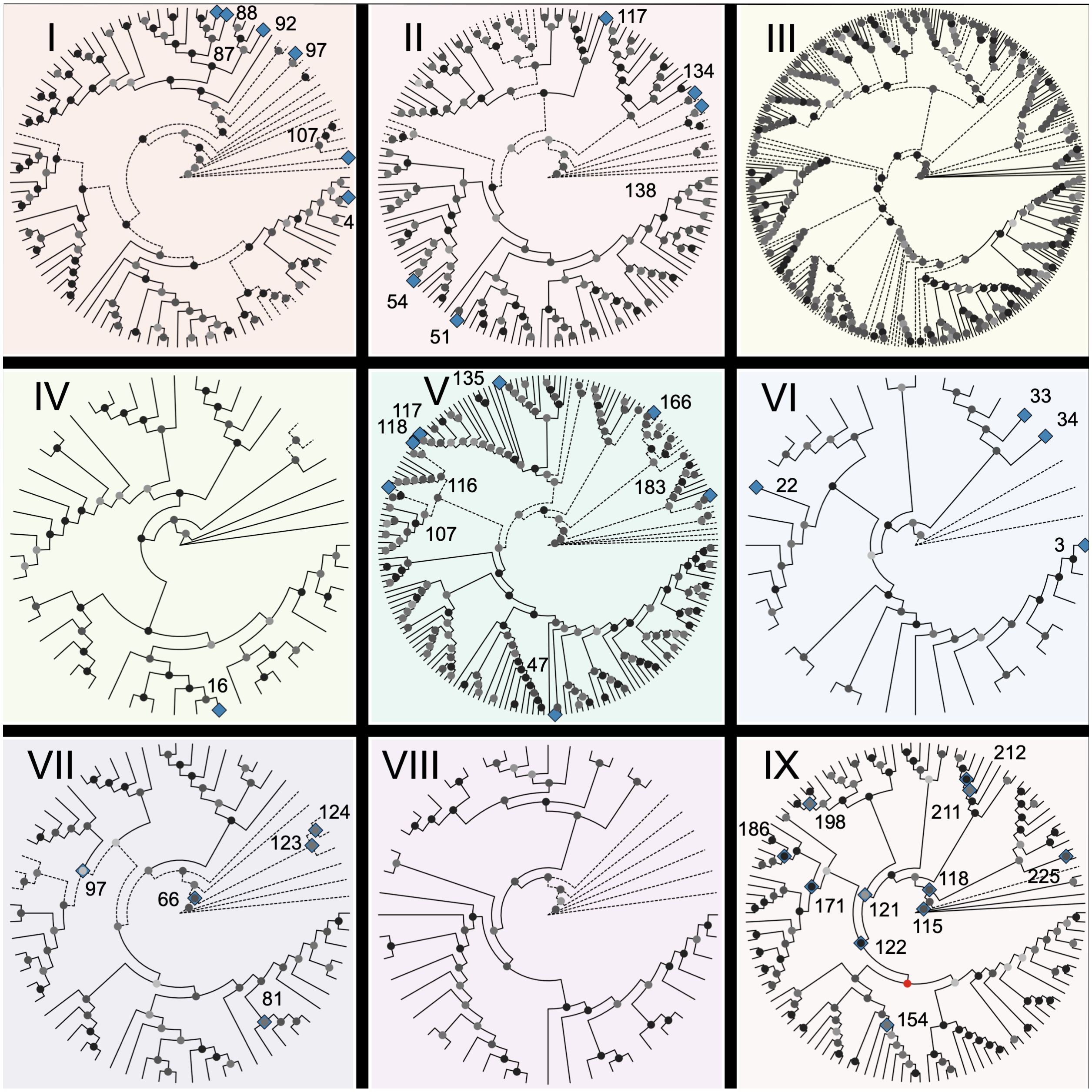

**Figure S7 – Repeat element density of V1R-containing and non-V1R-containing regions.** Regions of the genome containing V1Rs were analyzed for repeat element density and compared with random non-V1R containing regions to test for element enrichment in the cow, horse, mouse, and mouse lemur genomes. We estimated the percentage of the regions analyzed that are comprised of SINEs, LINEs, LTRs, and DNA transposons. Significance denoted by * for p<0.05 and *** for p<0.001.

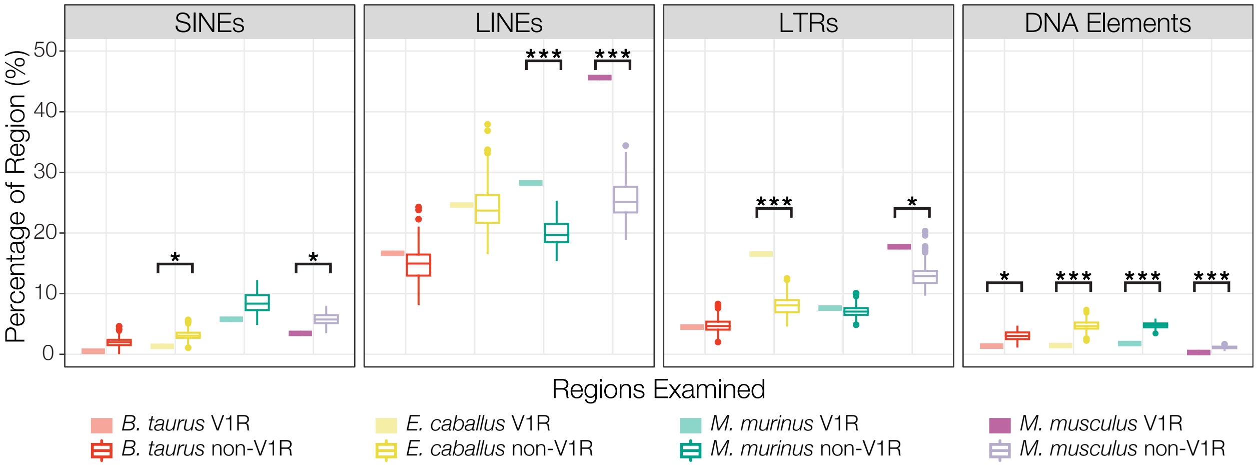

**Figure S8 – Syntenic relationships between mouse and dog.** Dog chromosomes are painted based on syntenic relationships with mouse. Locations of V1R genes are denoted in red.

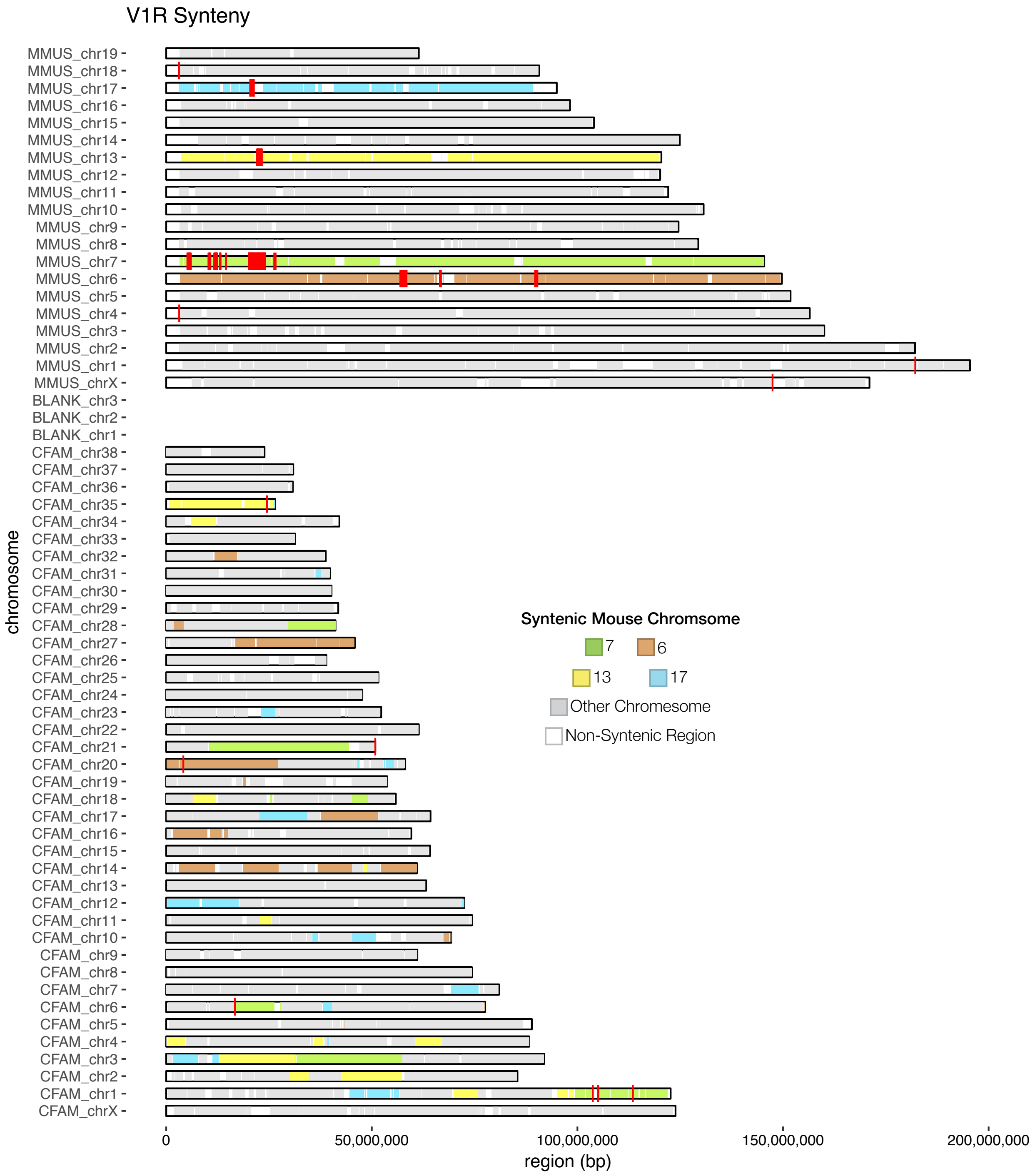

**Figure S9 – Syntenic relationships between mouse and horse.** Horse chromosomes are painted based on syntenic relationships with mouse. Locations of V1R genes are denoted in red.

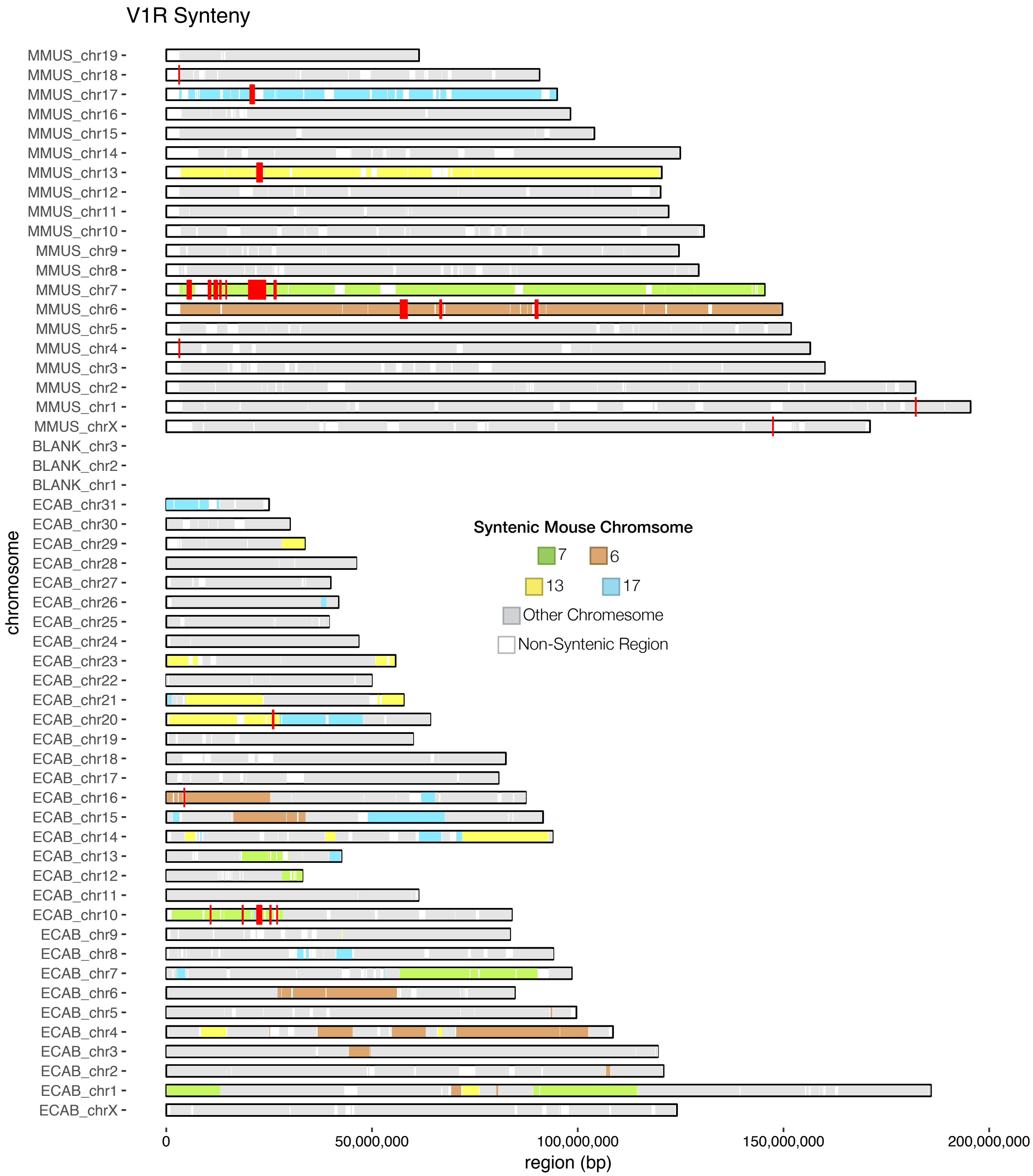

**Figure S10 – Syntenic relationships between mouse and human.** Human chromosomes are painted based on syntenic relationships with mouse. Locations of V1R genes are denoted in red.

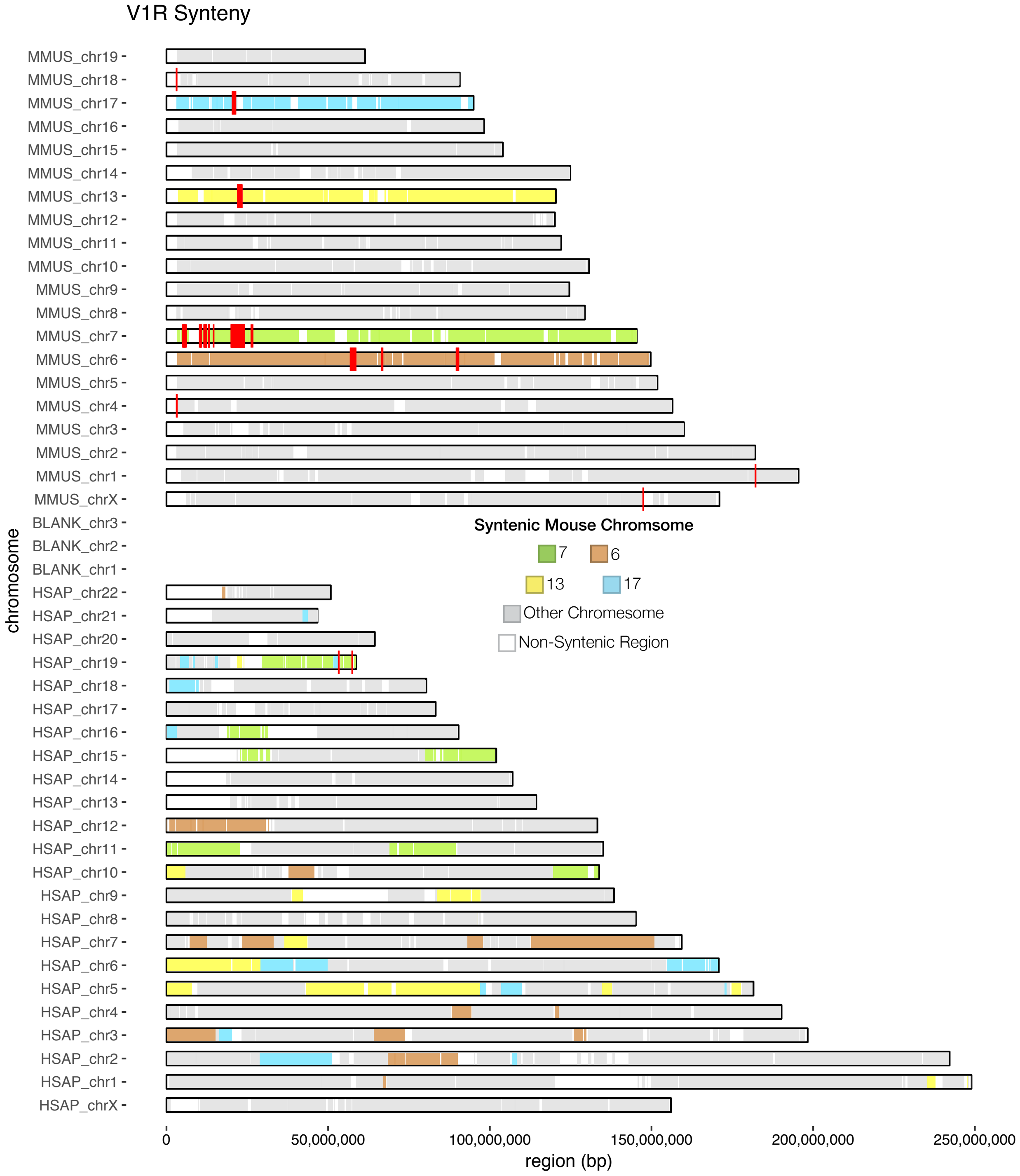

**Figure S11 – Syntenic relationships between mouse lemur and human.** Human chromosomes are painted based on syntenic relationships with mouse lemur. Locations of V1R genes are denoted in red. Putatively intact human V1Rs are labelled with gene names; VN1R1 and VN1R4 are the most likely V1Rs to have retained function in humans. Dashed box corresponds to "hotspot" chromosome in Figure 7.

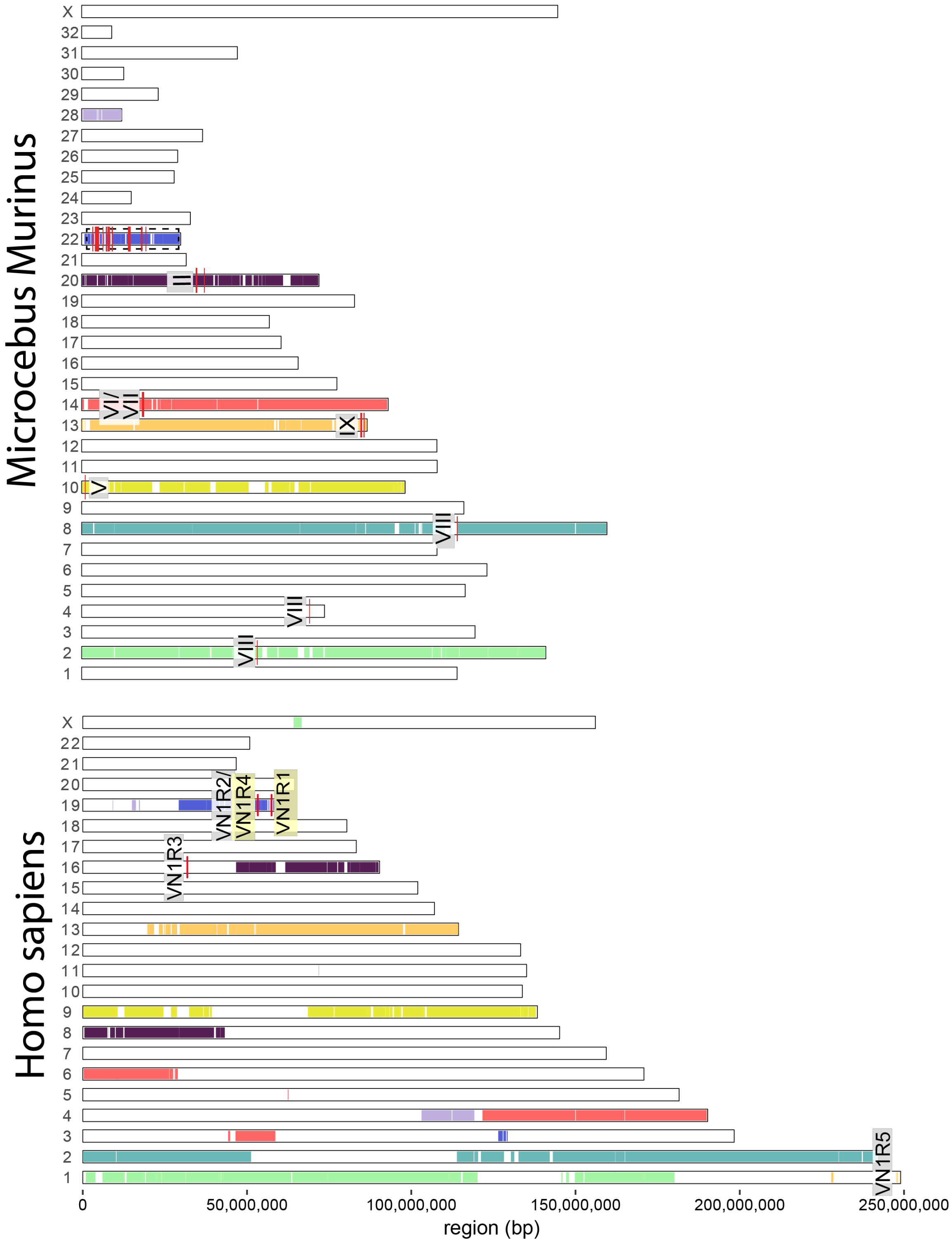

**Figure S12 – Intact and pseudogenized V1Rs in mouse and mouse lemur.** Mouse lemur chromosomes are painted based on syntenic relationships with mouse. Intact V1Rs indicated in red and pseudogenes in blue.

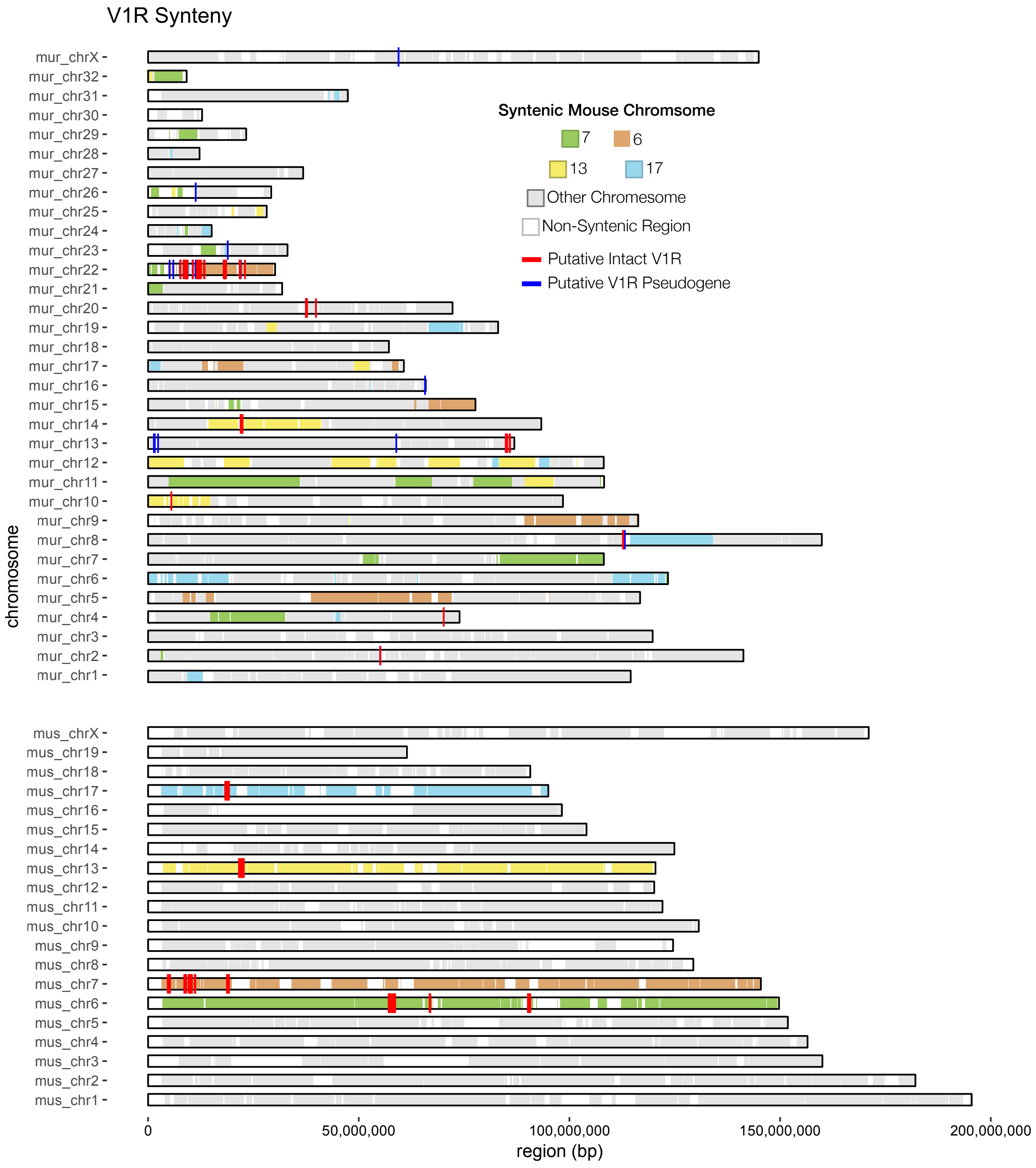

#### Supplementary Tables

**Table S1 –Genome assemblies used for analyses.** Gray rows indicate new dwarf or mouse lemur genomes.

| Species | Assembly Name | Assembly Accession | Total Sequence Length (Gb) | Contig N50 (bp) | Scaffold N50 (bp) | Coverage |
|---|---|---|---|---|---|---|
| *Bos taurus* | Bos_taurus_UMD_3.1.1 | GCF_000003055.6 | 2.67 | 98406 | 105708250* | 9x |
| *Callithrix jacchus* | Callithrix jacchus-3.2 | GCF_000004665.1 | 2.91 | 37523 | 132174527* | 6.6x |
| *Canis Familiaris* | CanFam3.1 | GCF_000002285.3 | 2.41 | 411364 | 63241923* | >7x |
| *Carlito syrichta* | Tarsius_syrichta_2.0.1 | GCF_164805.1 | 3.45 | 59907 | 401181 | 48x |
| *Cheirogaleus medius* | Cmed_1.0 | Pending | 2.22 | 36308 | 50625866 | 110x |
| *Cheirogaleus sibreei* | Csib_1.0 | Pending | 2.4 | 53909 | 54899 | 39x |
| *Daubentonia madagascariensis* | DauMad_1.0 | GCA_241425.1 | 2.86 | 3594 | 3653 | 38x |
| *Equus Caballus* | EquCab3.0 | GCF_002863925.1 | 2.51 | 4715787 | 87230776* | 88x |
| *Eulemur flavifrons* | Eflavifronsk33QCA | GCA_001262665.1 | 2.12 | 28371 | 413352 | 52.0x |
| *Eulemur macaco* | Emacaco_refEf_BWA_oneround | GCA_1262655.1 | 2.12 | 29783 | 405987 | 21.0x |
| *Galeopterus variegatus* | G_variegatus-3.0.2 | GCF_000696425.1 | 3.19 | 22275 | 245189 | 55x |
| *Gorilla gorilla* | gorGor4 | GCF_000151905.2 | 3.06 | 73808 | 146757320* | 80x |
| *Homo sapiens* | GRCh38.p12 | GCF_000001405.38 | 3.26 | 56413054 | 145138636* | NA |
| *Microcebus griseorufus* | Mgri_1.0 | Pending | 2.84 | 16706 | 16814 | 40x |
| *Microcebus mittermeieri* | Mmit_1.0 | Pending | 3.01 | 15313 | 15313 | 35x |
| *Microcebus murinus* | Mmur_2.0 | GCF_165445.1 | 2.44 | 182929 | 3711085 | 221.6x |
| *Microcebus murinus* | Mmur_3.0 | GCF_165445.2 | 2.49 | 234371 | 108171978* | 221.6x |
| *Microcebus murinus* | Mmur_DLC7028 | Pending | 2.48 | 40992 | 7199757 | 29x |
| *Microcebus murinus* | Mmur_DLC7032 | Pending | 2.4 | 52099 | 9443435 | 28x |
| *Microcebus murinus* | Mmur_DLC7033v1 | Pending | 2.4 | 43653 | 10213316 | 25x |
| *Microcebus murinus* | Mmur_DLC7033v2 | Pending | 2.4 | 50994 | 12158062 | 27x |
| *Microcebus murinus* | Mmur_DLC7039 | Pending | 2.34 | 51699 | 4172979 | 26x |
| *Microcebus murinus* | Mmur_DLC7049 | Pending | 2.26 | 24431 | 631600 | 21x |
| *Microcebus murinus* | Mmur_DLC7128 | Pending | 2.37 | 40965 | 7271319 | 24x |
| *Microcebus murinus* | Mmur_DLC7163 | Pending | 2.3 | 38087 | 1220679 | 24x |
| *Microcebus murinus* | Mmur_DLC7188 | Pending | 2.3 | 32375 | 973440 | 22x |
| *Microcebus ravelobensis* | Mrav_1.0 | Pending | 2.42 | 21470 | 21489 | 26x |
| *Microcebus tavaratra* | Mtav_1.0 | Pending | 2.58 | 29674 | 30035 | 41x |
| *Mirza zaza* | Mzaz_1.0 | Pending | 2.36 | 74231 | 75828 | 44x |
| *Mus musculus* | GRCm38.p6 | GCF_000001635.26 | 2.82 | 32273079 | 130694993* | NA |
| *Nomascus leucogenys* | Nleu_3.0 | GCA_000146795.3 | 2.96 | 35148 | 117381234 | 5.6x |
| *Otolemur garnetti* | OtoGar3 | GCF_181295.1 | 2.52 | 30487 | 13852661 | 137x |
| *Prolemur simus* | Prosim_1.0 | GCA_003258685.1 | 2.41 | 47760 | 2710671 | 153x |
| *Propithecus coquereli* | Pcoq_1.0 | GCF_000956105.1 | 2.8 | 28129 | 5604909 | 104.7x |
| *Tupaia chinensis* | TupChi_1.0 | GCF_000334495.1 | 2.85 | 30015 | 3670124 | 80x |

**Table S2 – Allele counts by lemur V1R subfamily.** Strepsirrhine primates shown in blue; haplorrhine primates in orange; outgroups in gray. Larsen et al. (2014) evaluated V1Rs for subfamily I and IX only.

| Species | I | II | III | IV | V | VI | VII | VIII | IX | Sum |
|---|---|---|---|---|---|---|---|---|---|---|
| *Otolemur garnetti* | 1 | 1 | 10 | 1 | 12 | 6 | 0 | 3 | 0 | 54 |
| *Daubentonia madagascariensis* | 0 | 0 | 6 | 0 | 0 | 1 | 0 | 1 | 1 | 17 |
| *Propithecus coquereli* | 6 | 4 | 10 | 0 | 3 | 1 | 0 | 0 | 0 | 27 |
| *Prolemur simus* | 1 | 6 | 6 | 0 | 5 | 1 | 0 | 0 | 0 | 22 |
| *Eulemur flavifrons* | 2 | 5 | 7 | 0 | 2 | 1 | 1 | 0 | 0 | 23 |
| *Eulemur macaco* | 2 | 5 | 7 | 0 | 3 | 1 | 1 | 0 | 0 | 25 |
| *Cheirogaleus medius* | 1 | 13 | 10 | 2 | 11 | 1 | 2 | 2 | 7 | 58 |
| *Cheirogaleus sibreei* | 8 | 15 | 14 | 2 | 12 | 2 | 2 | 4 | 6 | 70 |
| *Mirza zaza* | 8 | 10 | 26 | 3 | 16 | 5 | 5 | 2 | 8 | 95 |
| *Microcebus ravelobensis* | 14 | 15 | 25 | 8 | 24 | 8 | 3 | 4 | 17 | 129 |
| *Microcebus tavaratra* | 12 | 17 | 27 | 9 | 29 | 7 | 7 | 8 | 16 | 143 |
| *Microcebus griseus* | 15 | 15 | 17 | 4 | 23 | 12 | 5 | 10 | 25 | 137 |
| *Microcebus mittermeieri* | 17 | 21 | 22 | 10 | 26 | 7 | 4 | 9 | 22 | 146 |
| *Microcebus murinus* 3.0 (reference) | 11 | 12 | 17 | 7 | 17 | 10 | 5 | 3 | 12 | 102 |
| *Microcebus murinus* (Larsen et al., 2014) | 34-36 | - | - | - | - | - | - | - | 28-32 | - |
| *Microcebus murinus* (Young et al., 2010) | 29 | 7 | 13 | 4 | 14 | 9 | 4 | 5 | 19 | 105 |
| *Homo sapiens* | 0 | 0 | 1 | 0 | 0 | 0 | 0 | 0 | 0 | 2 |
| *Gorilla gorilla* | 0 | 0 | 0 | 0 | 0 | 0 | 0 | 0 | 0 | 2 |
| *Nomascus leucogenys* | 0 | 0 | 0 | 0 | 0 | 0 | 1 | 0 | 0 | 1 |
| *Callithrix jacchus* | 0 | 0 | 2 | 0 | 3 | 0 | 0 | 0 | 0 | 7 |
| *Carlito syrichta* | 1 | 0 | 10 | 0 | 4 | 0 | 0 | 0 | 0 | 28 |
| *Galeopterus variegatus* | 8 | 0 | 4 | 0 | 0 | 0 | 0 | 2 | 0 | 32 |
| *Tupaia chinensis* | 0 | 5 | 1 | 1 | 0 | 0 | 0 | 0 | 0 | 15 |

**Table S3 – AU p-values for three competing topological hypotheses for each lemur subfamily.** The three hypotheses are the topology estimated from the entire repertoire (Parsed Subtree), the topology estimated by ML from the original alignment when only analyzing taxa in each respective subfamily separately (MLE Subtree), and the MLE topology from the re-aligned data for each subfamily (Re-aligned Subtree). P-values < 0.0.5 should be interpreted as rejecting a topology from the plausible set.

| | Subfamily | Parsed Subtree | MLE Subtree | Re-aligned Subtree |
|---|---|---|---|---|
| **Original Alignment** | I | 0.194 | 0.688 | 0.524 |
| | II | 0.195 | 0.604 | 0.611 |
| | III | 0.398 | 0.622 | 0.428 |
| | IV | 0.433 | 0.525 | 0.511 |
| | V | 0.317 | 0.472 | 0.644 |
| | VI | 0.347 | 0.551 | 0.574 |
| | VII | 0.399 | 0.696 | 0.309 |
| | VIII | 0.465 | 0.447 | 0.535 |
| | IX | 0.217 | 0.569 | 0.605 |
| **Subfamily Re-Alignment** | I | 0.184 | 0.682 | 0.532 |
| | II | 0.149 | 0.614 | 0.618 |
| | III | 0.363 | 0.509 | 0.571 |
| | IV | 0.401 | 0.522 | 0.548 |
| | V | 0.347 | 0.171 | 0.783 |
| | VI | 0.374 | 0.533 | 0.562 |
| | VII | 0.269 | 0.45 | 0.667 |
| | VIII | 0.47 | 0.467 | 0.517 |
| | IX | 0.209 | 0.581 | 0.597 |

**Table S4 – Site model results for M2a and M1a comparisons.** Proportions of sites (p) and dN/dS for each proportion (ω) are given for each subfamily analysis across taxonomic filters. Missing rows mean that a taxonomic filter was equivalent the next-rank filter.

| | | Model M1a | | | | Model M2a | | | | | | | |
|---|---|---|---|---|---|---|---|---|---|---|---|---|---|
| **Subfamily** | **Filter** | $p_1$ | $p_2$ | $\omega_1$ | $\omega_2$ | $p_1$ | $p_2$ | $p_3$ | $\omega_1$ | $\omega_2$ | $\omega_3$ | **LRT** | **p-value** |
| I | Microcebus | 0.60 | 0.40 | 0.10 | 1 | 0.57 | 0.31 | 0.12 | 0.12 | 1 | 2.82 | 111.805934 | 3.9405E-26 |
| I | Cheirogaleidae | 0.57 | 0.43 | 0.11 | 1 | 0.54 | 0.33 | 0.12 | 0.12 | 1 | 2.67 | 137.943934 | 7.4964E-32 |
| I | Lemuriformes | 0.58 | 0.42 | 0.13 | 1 | 0.56 | 0.29 | 0.15 | 0.15 | 1 | 2.45 | 160.95441 | 7.0004E-37 |
| I | Strepsirrhini | 0.58 | 0.42 | 0.13 | 1 | 0.55 | 0.31 | 0.14 | 0.15 | 1 | 2.48 | 165.751974 | 6.2668E-38 |
| I | Primates | 0.59 | 0.41 | 0.14 | 1 | 0.56 | 0.30 | 0.14 | 0.16 | 1 | 2.42 | 157.485958 | 4.0083E-36 |
| I | Euarchontoglires | 0.58 | 0.42 | 0.17 | 1 | 0.53 | 0.33 | 0.13 | 0.18 | 1 | 2.41 | 165.310042 | 7.8268E-38 |
| II | Microcebus | 0.51 | 0.49 | 0.12 | 1 | 0.46 | 0.39 | 0.15 | 0.13 | 1 | 2.55 | 116.820924 | 3.1419E-27 |
| II | Cheirogaleidae | 0.55 | 0.45 | 0.17 | 1 | 0.46 | 0.41 | 0.14 | 0.16 | 1 | 2.65 | 242.259782 | 1.2647E-54 |
| II | Lemuriformes | 0.54 | 0.46 | 0.17 | 1 | 0.49 | 0.35 | 0.16 | 0.19 | 1 | 2.38 | 255.74758 | 1.4503E-57 |
| II | Strepsirrhini | 0.52 | 0.48 | 0.16 | 1 | 0.46 | 0.38 | 0.16 | 0.17 | 1 | 2.41 | 262.997026 | 3.8129E-59 |
| II | Primates | - | - | - | - | - | - | - | - | - | - | - | - |
| II | Euarchontoglires | 0.50 | 0.50 | 0.17 | 1 | 0.45 | 0.38 | 0.16 | 0.18 | 1 | 2.33 | 261.758322 | 7.0998E-59 |
| III | Microcebus | 0.47 | 0.53 | 0.27 | 1 | 0.45 | 0.46 | 0.09 | 0.29 | 1 | 2.09 | 30.298634 | 3.7039E-08 |
| III | Cheirogaleidae | 0.47 | 0.53 | 0.29 | 1 | 0.44 | 0.47 | 0.08 | 0.30 | 1 | 2.05 | 36.328468 | 1.6671E-09 |
| III | Lemuriformes | 0.42 | 0.58 | 0.29 | 1 | 0.38 | 0.51 | 0.11 | 0.29 | 1 | 2.21 | 86.534884 | 1.3729E-20 |
| III | Strepsirrhini | 0.44 | 0.56 | 0.31 | 1 | 0.39 | 0.50 | 0.11 | 0.31 | 1 | 2.27 | 136.735624 | 1.3776E-31 |
| III | Primates | 0.42 | 0.58 | 0.31 | 1 | 0.38 | 0.51 | 0.11 | 0.32 | 1 | 2.18 | 156.325702 | 7.1861E-36 |
| III | Euarchontoglires | 0.41 | 0.59 | 0.31 | 1 | 0.36 | 0.48 | 0.16 | 0.31 | 1 | 1.94 | 176.839356 | 2.3744E-40 |
| IV | Microcebus | 0.59 | 0.41 | 0.10 | 1 | 0.56 | 0.40 | 0.03 | 0.10 | 1 | 4.95 | 53.482136 | 2.6096E-13 |
| IV | Cheirogaleidae | 0.61 | 0.39 | 0.10 | 1 | 0.59 | 0.36 | 0.06 | 0.10 | 1 | 3.32 | 47.768638 | 4.796E-12 |
| IV | Lemuriformes | - | - | - | - | - | - | - | - | - | - | - | - |
| IV | Strepsirrhini | 0.62 | 0.38 | 0.14 | 1 | 0.60 | 0.35 | 0.06 | 0.15 | 1 | 3.38 | 59.238676 | 1.3966E-14 |
| IV | Primates | - | - | - | - | - | - | - | - | - | - | - | - |
| IV | Euarchontoglires | 0.60 | 0.40 | 0.15 | 1 | 0.58 | 0.37 | 0.05 | 0.16 | 1 | 3.26 | 54.28628 | 1.7331E-13 |
| V | Microcebus | 0.53 | 0.47 | 0.18 | 1 | 0.49 | 0.41 | 0.10 | 0.19 | 1 | 2.46 | 90.854152 | 1.5466E-21 |
| V | Cheirogaleidae | 0.56 | 0.44 | 0.21 | 1 | 0.52 | 0.36 | 0.12 | 0.22 | 1 | 2.32 | 158.032934 | 3.044E-36 |
| V | Lemuriformes | 0.56 | 0.44 | 0.23 | 1 | 0.53 | 0.35 | 0.12 | 0.25 | 1 | 2.24 | 171.736126 | 3.0902E-39 |
| V | Strepsirrhini | 0.54 | 0.46 | 0.24 | 1 | 0.50 | 0.34 | 0.16 | 0.27 | 1 | 2.20 | 233.18608 | 1.2037E-52 |
| V | Primates | 0.53 | 0.47 | 0.25 | 1 | 0.48 | 0.37 | 0.16 | 0.27 | 1 | 2.24 | 274.17196 | 1.3985E-61 |
| V | Euarchontoglires | - | - | - | - | - | - | - | - | - | - | - | - |
| VI | Microcebus | 0.41 | 0.59 | 0.00 | 1 | 0.39 | 0.50 | 0.11 | 0.00 | 1 | 3.51 | 19.380746 | 0.000010708 |
| VI | Cheirogaleidae | 0.49 | 0.51 | 0.12 | 1 | 0.44 | 0.51 | 0.04 | 0.12 | 1 | 4.61 | 42.972056 | 5.5527E-11 |
| VI | Lemuriformes | 0.47 | 0.53 | 0.14 | 1 | 0.42 | 0.54 | 0.04 | 0.14 | 1 | 4.70 | 39.72317 | 2.9263E-10 |
| VI | Strepsirrhini | - | - | - | - | - | - | - | - | - | - | - | - |
| VI | Primates | 0.48 | 0.52 | 0.16 | 1 | 0.43 | 0.54 | 0.03 | 0.16 | 1 | 5.45 | 46.81382 | 7.8061E-12 |
| VI | Euarchontoglires | - | - | - | - | - | - | - | - | - | - | - | - |
| VII | Microcebus | 0.64 | 0.36 | 0.13 | 1 | 0.70 | 0.17 | 0.13 | 0.20 | 1 | 2.50 | 29.510204 | 5.5623E-08 |
| VII | Cheirogaleidae | 0.65 | 0.35 | 0.16 | 1 | 0.66 | 0.28 | 0.06 | 0.19 | 1 | 2.85 | 28.2485 | 1.067E-07 |
| VII | Lemuriformes | 0.63 | 0.37 | 0.15 | 1 | 0.62 | 0.35 | 0.04 | 0.16 | 1 | 3.21 | 24.418524 | 7.752E-07 |
| VII | Strepsirrhini | 0.49 | 0.51 | 0.16 | 1 | 0.47 | 0.45 | 0.08 | 0.16 | 1 | 2.54 | 33.064448 | 8.9154E-09 |
| VII | Primates | - | - | - | - | - | - | - | - | - | - | - | - |
| VII | Euarchontoglires | - | - | - | - | - | - | - | - | - | - | - | - |
| VIII | Microcebus | 0.51 | 0.49 | 0.02 | 1 | 0.62 | 0.19 | 0.19 | 0.13 | 1 | 2.87 | 34.796766 | 3.6598E-09 |
| VIII | Cheirogaleidae | 0.51 | 0.49 | 0.11 | 1 | 0.51 | 0.35 | 0.14 | 0.16 | 1 | 2.91 | 40.058136 | 2.4652E-10 |
| VIII | Lemuriformes | - | - | - | - | - | - | - | - | - | - | - | - |
| VIII | Strepsirrhini | 0.52 | 0.48 | 0.20 | 1 | 0.49 | 0.37 | 0.14 | 0.26 | 1 | 2.99 | 64.793112 | 8.3189E-16 |
| VIII | Primates | - | - | - | - | - | - | - | - | - | - | - | - |
| VIII | Euarchontoglires | 0.51 | 0.49 | 0.25 | 1 | 0.49 | 0.35 | 0.15 | 0.31 | 1 | 2.62 | 57.357012 | 3.6347E-14 |
| IX | Microcebus | 0.61 | 0.39 | 0.11 | 1 | 0.57 | 0.26 | 0.16 | 0.12 | 1 | 2.99 | 245.664162 | 2.2894E-55 |
| IX | Cheirogaleidae | 0.60 | 0.40 | 0.13 | 1 | 0.54 | 0.29 | 0.17 | 0.15 | 1 | 2.94 | 329.572516 | 1.191E-73 |
| IX | Lemuriformes | - | - | - | - | - | - | - | - | - | - | - | - |
| IX | Strepsirrhini | 0.60 | 0.40 | 0.14 | 1 | 0.55 | 0.28 | 0.18 | 0.16 | 1 | 2.87 | 329.832496 | 1.0454E-73 |
| IX | Primates | - | - | - | - | - | - | - | - | - | - | - | - |
| IX | Euarchontoglires | - | - | - | - | - | - | - | - | - | - | - | - |

**Table S5 – Site model results for M8 and M7 comparisons.** Proportions of sites (p) and dN/dS for each proportion (ω) are given for each subfamily analysis across taxonomic filters. Missing rows mean that a taxonomic filter was equivalent the next-rank filter.

| | | Model M7 | | Model M8 | | | | | | |
|---|---|---|---|---|---|---|---|---|---|---|
| Subfamily | Filter | α | β | $p_{<=1}$ | $p_{>1}$ | α | β | ω> 1 | LRT | p-value |
| I | Microcebus | 0.26 | 0.31 | 0.84 | 0.16 | 0.4 | 0.71 | 2.47 | 137.392636 | 9.90E-32 |
| I | Cheirogaleidae | 0.27 | 0.31 | 0.85 | 0.15 | 0.36 | 0.57 | 2.37 | 157.565802 | 3.85E-36 |
| I | Lemuriformes | 0.33 | 0.37 | 0.81 | 0.19 | 0.51 | 0.9 | 2.17 | 197.675886 | 6.71E-45 |
| I | Strepsirrhini | 0.34 | 0.38 | 0.82 | 0.18 | 0.5 | 0.84 | 2.18 | 196.47499 | 1.23E-44 |
| I | Primates | 0.37 | 0.41 | 0.82 | 0.18 | 0.56 | 0.97 | 2.1 | 192.406904 | 9.48E-44 |
| I | Euarchontoglires | 0.45 | 0.47 | 0.83 | 0.17 | 0.64 | 0.96 | 2.05 | 193.734876 | 4.87E-44 |
| II | Microcebus | 0.3 | 0.27 | 0.82 | 0.18 | 0.43 | 0.55 | 2.28 | 137.184348 | 1.10E-31 |
| II | Cheirogaleidae | 0.38 | 0.37 | 0.83 | 0.17 | 0.51 | 0.64 | 2.27 | 251.543584 | 1.20E-56 |
| II | Lemuriformes | 0.42 | 0.37 | 0.8 | 0.2 | 0.59 | 0.72 | 2.14 | 287.17727 | 2.05E-64 |
| II | Strepsirrhini | 0.43 | 0.36 | 0.81 | 0.19 | 0.58 | 0.68 | 2.13 | 280.149164 | 6.97E-63 |
| II | Primates | - | - | - | - | - | - | - | - | - |
| II | Euarchontoglires | 0.47 | 0.38 | 0.81 | 0.19 | 0.62 | 0.69 | 2.07 | 274.870922 | 9.85E-62 |
| III | Microcebus | 0.82 | 0.52 | 0.81 | 0.19 | 1.15 | 1.04 | 1.63 | 53.054524 | 3.24E-13 |
| III | Cheirogaleidae | 0.87 | 0.56 | 0.84 | 0.16 | 1.1 | 0.93 | 1.6 | 56.213286 | 6.50E-14 |
| III | Lemuriformes | 0.81 | 0.48 | 0.86 | 0.14 | 0.96 | 0.7 | 1.79 | 95.212948 | 1.71E-22 |
| III | Strepsirrhini | 0.81 | 0.48 | 0.86 | 0.14 | 0.95 | 0.67 | 1.84 | 138.378504 | 6.023E-32 |
| III | Primates | 0.84 | 0.49 | 0.84 | 0.16 | 1 | 0.71 | 1.73 | 152.993582 | 3.84E-35 |
| III | Euarchontoglires | 0.82 | 0.49 | 0.82 | 0.18 | 1.01 | 0.76 | 1.66 | 172.539424 | 2.06E-39 |
| IV | Microcebus | 0.18 | 0.2 | 0.92 | 0.08 | 0.31 | 0.44 | 3.29 | 60.529864 | 7.25E-15 |
| IV | Cheirogaleidae | 0.22 | 0.28 | 0.92 | 0.08 | 0.31 | 0.5 | 2.8 | 58.369812 | 2.17E-14 |
| IV | Lemuriformes | - | - | - | - | - | - | - | - | - |
| IV | Strepsirrhini | 0.34 | 0.4 | 0.91 | 0.09 | 0.52 | 0.77 | 2.74 | 76.188962 | 2.58E-18 |
| IV | Primates | - | - | - | - | - | - | - | - | - |
| IV | Euarchontoglires | 0.41 | 0.45 | 0.91 | 0.09 | 0.58 | 0.81 | 2.59 | 71.923442 | 2.24E-17 |
| V | Microcebus | 0.47 | 0.4 | 0.81 | 0.19 | 0.7 | 0.87 | 1.93 | 113.537168 | 1.65E-26 |
| V | Cheirogaleidae | 0.57 | 0.51 | 0.85 | 0.15 | 0.78 | 0.91 | 2.04 | 186.14484 | 2.21E-42 |
| V | Lemuriformes | 0.64 | 0.54 | 0.84 | 0.16 | 0.93 | 1.03 | 1.96 | 210.413496 | 1.12E-47 |
| V | Strepsirrhini | 0.66 | 0.51 | 0.81 | 0.19 | 0.94 | 0.99 | 1.96 | 269.719446 | 1.31E-60 |
| V | Primates | 0.71 | 0.54 | 0.82 | 0.18 | 1 | 1 | 1.96 | 295.790652 | 2.72E-66 |
| V | Euarchontoglires | - | - | - | - | - | - | - | - | - |
| VI | Microcebus | 0.01 | 0.01 | 0.87 | 0.13 | 0.03 | 0.03 | 3.32 | 19.33141 | 0.000010988 |
| VI | Cheirogaleidae | 0.21 | 0.15 | 0.94 | 0.06 | 0.27 | 0.2 | 4.21 | 47.128876 | 6.65E-12 |
| VI | Lemuriformes | 0.25 | 0.17 | 0.95 | 0.05 | 0.32 | 0.21 | 4.17 | 43.464826 | 4.32E-11 |
| VI | Strepsirrhini | - | - | - | - | - | - | - | - | - |
| VI | Primates | 0.33 | 0.21 | 0.96 | 0.04 | 0.35 | 0.22 | 5.08 | 51.081888 | 8.86E-13 |
| VI | Euarchontoglires | - | - | - | - | - | - | - | - | - |
| VII | Microcebus | 0.26 | 0.29 | 0.81 | 0.19 | 2.7 | 7.43 | 2.18 | 43.343834 | 4.59E-11 |
| VII | Cheirogaleidae | 0.42 | 0.48 | 0.88 | 0.12 | 1.08 | 1.98 | 2.26 | 45.243112 | 1.74E-11 |
| VII | Lemuriformes | 0.39 | 0.46 | 0.9 | 0.1 | 0.68 | 1.07 | 2.24 | 36.053678 | 1.92E-09 |
| VII | Strepsirrhini | 0.43 | 0.35 | 0.86 | 0.14 | 0.62 | 0.7 | 2.07 | 47.219908 | 6.35E-12 |
| VII | Primates | - | - | - | - | - | - | - | - | - |
| VII | Euarchontoglires | - | - | - | - | - | - | - | - | - |
| VIII | Microcebus | 0.02 | 0.02 | 0.76 | 0.24 | 0.75 | 2.24 | 2.67 | 34.656634 | 3.93E-09 |
| VIII | Cheirogaleidae | 0.16 | 0.11 | 0.82 | 0.18 | 0.56 | 0.69 | 2.65 | 48.01802 | 4.22E-12 |
| VIII | Lemuriformes | - | - | - | - | - | - | - | - | - |
| VIII | Strepsirrhini | 0.54 | 0.36 | 0.83 | 0.17 | 1.09 | 1.04 | 2.7 | 89.677224 | 2.80E-21 |
| VIII | Primates | - | - | - | - | - | - | - | - | - |
| VIII | Euarchontoglires | 0.63 | 0.37 | 0.82 | 0.18 | 1.2 | 1 | 2.43 | 80.584932 | 2.78E-19 |
| IX | Microcebus | 0.29 | 0.37 | 0.82 | 0.18 | 0.45 | 0.93 | 2.65 | 280.7036 | 5.28E-63 |
| IX | Cheirogaleidae | 0.32 | 0.36 | 0.81 | 0.19 | 0.46 | 0.76 | 2.63 | 356.988518 | 1.27E-79 |
| IX | Lemuriformes | - | - | - | - | - | - | - | - | - |
| IX | Strepsirrhini | 0.35 | 0.38 | 0.81 | 0.19 | 0.51 | 0.83 | 2.6 | 365.349952 | 1.93E-81 |
| IX | Primates | - | - | - | - | - | - | - | - | - |
| IX | Euarchontoglires | - | - | - | - | - | - | - | - | - |

**Table S6 – Site model results for M2a and M1a comparisons with realigned subfamilies.** Proportions of sites (p) and dN/dS for each proportion (ω) are given for each subfamily analysis across taxonomic filters. Missing rows mean that a taxonomic filter was equivalent the next-rank filter.

| | | Model M1a | | | | Model M2a | | | | | | | |
|---|---|---|---|---|---|---|---|---|---|---|---|---|---|
| **Subfamily** | **Filter** | **$p_1$** | **$p_2$** | **$\omega_1$** | **$\omega_2$** | **$p_1$** | **$p_2$** | **$p_3$** | **$\omega_1$** | **$\omega_2$** | **$\omega_3$** | **LRT** | **p-value** |
| I | Microcebus | 0.60 | 0.40 | 0.10 | 1.00 | 0.57 | 0.31 | 0.12 | 0.12 | 1.00 | 2.86 | 122.50463 | 1.79E-28 |
| I | Cheirogaleidae | 0.57 | 0.43 | 0.12 | 1.00 | 0.54 | 0.33 | 0.13 | 0.13 | 1.00 | 2.68 | 147.000558 | 7.85E-34 |
| I | Lemuriformes | 0.59 | 0.41 | 0.13 | 1.00 | 0.56 | 0.29 | 0.16 | 0.15 | 1.00 | 2.50 | 173.567412 | 1.23E-39 |
| I | Strepsirrhini | 0.58 | 0.42 | 0.13 | 1.00 | 0.54 | 0.31 | 0.15 | 0.15 | 1.00 | 2.54 | 179.88805 | 5.13E-41 |
| I | Primates | 0.59 | 0.41 | 0.14 | 1.00 | 0.55 | 0.30 | 0.14 | 0.16 | 1.00 | 2.47 | 170.458462 | 5.88E-39 |
| I | Euarchontoglires | 0.58 | 0.42 | 0.17 | 1.00 | 0.53 | 0.33 | 0.14 | 0.18 | 1.00 | 2.44 | 173.354098 | 1.37E-39 |
| II | Microcebus | 0.51 | 0.49 | 0.13 | 1.00 | 0.46 | 0.39 | 0.15 | 0.13 | 1.00 | 2.54 | 116.795128 | 3.18E-27 |
| II | Cheirogaleidae | 0.55 | 0.45 | 0.17 | 1.00 | 0.46 | 0.41 | 0.13 | 0.16 | 1.00 | 2.66 | 247.423316 | 9.47E-56 |
| II | Lemuriformes | 0.54 | 0.46 | 0.17 | 1.00 | 0.49 | 0.34 | 0.17 | 0.19 | 1.00 | 2.41 | 267.714142 | 3.57E-60 |
| II | Strepsirrhini | 0.52 | 0.48 | 0.17 | 1.00 | 0.47 | 0.37 | 0.16 | 0.18 | 1.00 | 2.44 | 274.394682 | 1.25E-61 |
| II | Primates | - | - | - | - | - | - | - | - | - | - | - | - |
| II | Euarchontoglires | 0.50 | 0.50 | 0.17 | 1.00 | 0.46 | 0.38 | 0.16 | 0.18 | 1.00 | 2.36 | 274.093382 | 1.45E-61 |
| III | Microcebus | 0.48 | 0.52 | 0.27 | 1.00 | 0.45 | 0.47 | 0.08 | 0.28 | 1.00 | 2.26 | 39.516662 | 3.25E-10 |
| III | Cheirogaleidae | 0.48 | 0.52 | 0.29 | 1.00 | 0.46 | 0.50 | 0.04 | 0.29 | 1.00 | 3.27 | 55.03314 | 1.19E-13 |
| III | Lemuriformes | 0.44 | 0.56 | 0.29 | 1.00 | 0.40 | 0.50 | 0.10 | 0.29 | 1.00 | 2.23 | 105.810392 | 8.11E-25 |
| III | Strepsirrhini | 0.46 | 0.54 | 0.31 | 1.00 | 0.42 | 0.48 | 0.10 | 0.31 | 1.00 | 2.29 | 154.07496 | 2.23E-35 |
| III | Primates | 0.45 | 0.55 | 0.32 | 1.00 | 0.40 | 0.49 | 0.11 | 0.32 | 1.00 | 2.21 | 179.411586 | 6.51E-41 |
| III | Euarchontoglires | 0.43 | 0.57 | 0.31 | 1.00 | 0.38 | 0.46 | 0.16 | 0.32 | 1.00 | 1.96 | 198.090052 | 5.45E-45 |
| IV | Microcebus | 0.60 | 0.40 | 0.10 | 1.00 | 0.58 | 0.38 | 0.04 | 0.11 | 1.00 | 4.32 | 53.61382 | 2.44E-13 |
| IV | Cheirogaleidae | 0.61 | 0.39 | 0.10 | 1.00 | 0.59 | 0.35 | 0.06 | 0.11 | 1.00 | 3.42 | 54.955884 | 1.23E-13 |
| IV | Lemuriformes | - | - | - | - | - | - | - | - | - | - | - | - |
| IV | Strepsirrhini | 0.62 | 0.38 | 0.14 | 1.00 | 0.59 | 0.35 | 0.06 | 0.15 | 1.00 | 3.52 | 64.523152 | 9.54E-16 |
| IV | Primates | - | - | - | - | - | - | - | - | - | - | - | - |
| IV | Euarchontoglires | 0.60 | 0.40 | 0.15 | 1.00 | 0.58 | 0.36 | 0.06 | 0.16 | 1.00 | 3.41 | 63.486246 | 1.61E-15 |
| V | Microcebus | 0.52 | 0.48 | 0.18 | 1.00 | 0.49 | 0.42 | 0.09 | 0.19 | 1.00 | 2.43 | 84.930738 | 3.09E-20 |
| V | Cheirogaleidae | 0.54 | 0.46 | 0.20 | 1.00 | 0.50 | 0.37 | 0.13 | 0.22 | 1.00 | 2.30 | 151.910276 | 6.63E-35 |
| V | Lemuriformes | 0.55 | 0.45 | 0.23 | 1.00 | 0.51 | 0.36 | 0.13 | 0.25 | 1.00 | 2.24 | 167.123622 | 3.14E-38 |
| V | Strepsirrhini | 0.53 | 0.47 | 0.24 | 1.00 | 0.50 | 0.35 | 0.15 | 0.27 | 1.00 | 2.21 | 230.308924 | 5.10E-52 |
| V | Primates | 0.51 | 0.49 | 0.25 | 1.00 | 0.47 | 0.37 | 0.15 | 0.27 | 1.00 | 2.27 | 280.939722 | 4.69E-63 |
| V | Euarchontoglires | - | - | - | - | - | - | - | - | - | - | - | - |
| VI | Microcebus | 0.40 | 0.60 | 0.00 | 1.00 | 0.39 | 0.52 | 0.10 | 0.00 | 1.00 | 3.53 | 16.811742 | 0.000041277 |
| VI | Cheirogaleidae | 0.48 | 0.52 | 0.12 | 1.00 | 0.44 | 0.51 | 0.05 | 0.12 | 1.00 | 4.23 | 32.927292 | 9.57E-09 |
| VI | Lemuriformes | 0.46 | 0.54 | 0.13 | 1.00 | 0.42 | 0.53 | 0.04 | 0.13 | 1.00 | 4.25 | 32.307594 | 1.32E-08 |
| VI | Strepsirrhini | - | - | - | - | - | - | - | - | - | - | - | - |
| VI | Primates | 0.46 | 0.54 | 0.15 | 1.00 | 0.42 | 0.55 | 0.03 | 0.15 | 1.00 | 4.84 | 37.918052 | 7.38E-10 |
| VI | Euarchontoglires | - | - | - | - | - | - | - | - | - | - | - | - |
| VII | Microcebus | 0.65 | 0.35 | 0.12 | 1.00 | 0.75 | 0.07 | 0.18 | 0.21 | 1.00 | 2.29 | 34.254512 | 4.84E-09 |
| VII | Cheirogaleidae | 0.65 | 0.35 | 0.15 | 1.00 | 0.68 | 0.23 | 0.10 | 0.19 | 1.00 | 2.36 | 23.969488 | 9.79E-07 |
| VII | Lemuriformes | 0.63 | 0.37 | 0.14 | 1.00 | 0.63 | 0.33 | 0.04 | 0.16 | 1.00 | 2.86 | 19.910532 | 8.12E-06 |
| VII | Strepsirrhini | 0.50 | 0.50 | 0.15 | 1.00 | 0.48 | 0.44 | 0.08 | 0.16 | 1.00 | 2.54 | 30.027 | 4.26E-08 |
| VII | Primates | - | - | - | - | - | - | - | - | - | - | - | - |
| VII | Euarchontoglires | - | - | - | - | - | - | - | - | - | - | - | - |
| VIII | Microcebus | 0.51 | 0.49 | 0.02 | 1.00 | 0.62 | 0.19 | 0.19 | 0.13 | 1.00 | 2.87 | 34.796754 | 3.66E-09 |
| VIII | Cheirogaleidae | 0.51 | 0.49 | 0.11 | 1.00 | 0.51 | 0.35 | 0.14 | 0.16 | 1.00 | 2.91 | 40.058122 | 2.47E-10 |
| VIII | Lemuriformes | - | - | - | - | - | - | - | - | - | - | - | - |
| VIII | Strepsirrhini | 0.53 | 0.47 | 0.21 | 1.00 | 0.50 | 0.37 | 0.13 | 0.26 | 1.00 | 2.96 | 63.007258 | 2.06E-15 |
| VIII | Primates | - | - | - | - | - | - | - | - | - | - | - | - |
| VIII | Euarchontoglires | 0.51 | 0.49 | 0.25 | 1.00 | 0.50 | 0.35 | 0.15 | 0.31 | 1.00 | 2.59 | 55.772076 | 8.14E-14 |
| IX | Microcebus | 0.61 | 0.39 | 0.10 | 1.00 | 0.57 | 0.26 | 0.16 | 0.12 | 1.00 | 3.06 | 255.83814 | 1.39E-57 |
| IX | Cheirogaleidae | 0.60 | 0.40 | 0.13 | 1.00 | 0.55 | 0.28 | 0.17 | 0.15 | 1.00 | 2.95 | 333.087086 | 2.04E-74 |
| IX | Lemuriformes | - | - | - | - | - | - | - | - | - | - | - | - |
| IX | Strepsirrhini | 0.60 | 0.40 | 0.14 | 1.00 | 0.55 | 0.27 | 0.18 | 0.16 | 1.00 | 2.91 | 342.395736 | 1.92E-76 |
| IX | Primates | - | - | - | - | - | - | - | - | - | - | - | - |
| IX | Euarchontoglires | - | - | - | - | - | - | - | - | - | - | - | - |

**Table S7 – Site model results for M8 and M7 comparisons with realigned subfamilies.** Proportions of sites (p) and dN/dS for each proportion (ω) are given for each subfamily analysis across taxonomic filters. Missing rows mean that a taxonomic filter was equivalent the next-rank filter.

| | | Model M7 | | Model M8 | | | | | | |
|---|---|---|---|---|---|---|---|---|---|---|
| Subfamily | Filter | α | β | $p_{<=1}$ | $p_{>1}$ | α | β | ω> 1 | LRT | p-value |
| I | Microcebus | 0.26 | 0.31 | 0.83 | 0.17 | 0.41 | 0.74 | 2.53 | 150.170338 | 1.59E-34 |
| I | Cheirogaleidae | 0.27 | 0.31 | 0.84 | 0.16 | 0.37 | 0.60 | 2.39 | 167.935146 | 2.09E-38 |
| I | Lemuriformes | 0.33 | 0.37 | 0.81 | 0.19 | 0.52 | 0.94 | 2.22 | 212.262326 | 4.41E-48 |
| I | Strepsirrhini | 0.34 | 0.38 | 0.82 | 0.18 | 0.51 | 0.84 | 2.24 | 210.760468 | 9.37E-48 |
| I | Primates | 0.37 | 0.42 | 0.82 | 0.18 | 0.57 | 0.98 | 2.15 | 205.612676 | 1.24E-46 |
| I | Euarchontoglires | 0.45 | 0.47 | 0.82 | 0.18 | 0.66 | 1.00 | 2.06 | 202.948876 | 4.75E-46 |
| II | Microcebus | 0.30 | 0.27 | 0.82 | 0.18 | 0.44 | 0.56 | 2.28 | 137.529682 | 9.24E-32 |
| II | Cheirogaleidae | 0.38 | 0.37 | 0.82 | 0.18 | 0.53 | 0.66 | 2.27 | 259.714792 | 1.98E-58 |
| II | Lemuriformes | 0.42 | 0.36 | 0.80 | 0.20 | 0.60 | 0.72 | 2.17 | 300.977164 | 2.02E-67 |
| II | Strepsirrhini | 0.43 | 0.36 | 0.80 | 0.20 | 0.59 | 0.69 | 2.15 | 294.108412 | 6.33E-66 |
| II | Primates | - | - | - | - | - | - | - | - | - |
| II | Euarchontoglires | 0.47 | 0.38 | 0.81 | 0.19 | 0.63 | 0.69 | 2.11 | 288.757602 | 9.27E-65 |
| III | Microcebus | 0.81 | 0.52 | 0.84 | 0.16 | 1.09 | 0.94 | 1.74 | 60.595118 | 7.01E-15 |
| III | Cheirogaleidae | 0.86 | 0.57 | 0.87 | 0.13 | 1.06 | 0.87 | 1.72 | 69.257938 | 8.64E-17 |
| III | Lemuriformes | 0.81 | 0.50 | 0.87 | 0.13 | 0.96 | 0.71 | 1.85 | 113.20616 | 1.94E-26 |
| III | Strepsirrhini | 0.82 | 0.50 | 0.86 | 0.14 | 0.96 | 0.70 | 1.84 | 157.63547 | 3.7179E-36 |
| III | Primates | 0.84 | 0.50 | 0.85 | 0.15 | 0.99 | 0.72 | 1.79 | 176.906024 | 2.30E-40 |
| III | Euarchontoglires | 0.81 | 0.50 | 0.82 | 0.18 | 1.01 | 0.78 | 1.69 | 193.914718 | 4.45E-44 |
| IV | Microcebus | 0.19 | 0.22 | 0.92 | 0.08 | 0.34 | 0.50 | 3.27 | 61.911974 | 3.59E-15 |
| IV | Cheirogaleidae | 0.22 | 0.29 | 0.91 | 0.09 | 0.33 | 0.54 | 2.88 | 66.474776 | 3.54E-16 |
| IV | Lemuriformes | - | - | - | - | - | - | - | - | - |
| IV | Strepsirrhini | 0.35 | 0.41 | 0.91 | 0.09 | 0.52 | 0.76 | 2.91 | 81.651228 | 1.62E-19 |
| IV | Primates | - | - | - | - | - | - | - | - | - |
| IV | Euarchontoglires | 0.39 | 0.44 | 0.90 | 0.10 | 0.55 | 0.77 | 2.79 | 82.09013 | 1.30E-19 |
| V | Microcebus | 0.47 | 0.39 | 0.81 | 0.19 | 0.68 | 0.82 | 1.90 | 105.453038 | 9.72E-25 |
| V | Cheirogaleidae | 0.56 | 0.49 | 0.84 | 0.16 | 0.76 | 0.86 | 2.03 | 177.625352 | 1.60E-40 |
| V | Lemuriformes | 0.64 | 0.52 | 0.83 | 0.17 | 0.91 | 0.99 | 1.95 | 202.804436 | 5.10E-46 |
| V | Strepsirrhini | 0.66 | 0.50 | 0.81 | 0.19 | 0.93 | 0.97 | 1.95 | 263.943558 | 2.37E-59 |
| V | Primates | 0.71 | 0.53 | 0.82 | 0.18 | 0.97 | 0.93 | 2.00 | 296.61835 | 1.80E-66 |
| V | Euarchontoglires | - | - | - | - | - | - | - | - | - |
| VI | Microcebus | 0.01 | 0.01 | 0.89 | 0.11 | 0.02 | 0.02 | 3.37 | 16.584434 | 0.000046531 |
| VI | Cheirogaleidae | 0.21 | 0.15 | 0.94 | 0.06 | 0.28 | 0.20 | 3.88 | 36.764068 | 1.33E-09 |
| VI | Lemuriformes | 0.25 | 0.16 | 0.95 | 0.05 | 0.32 | 0.22 | 3.86 | 35.786758 | 2.20E-09 |
| VI | Strepsirrhini | - | - | - | - | - | - | - | - | - |
| VI | Primates | 0.32 | 0.21 | 0.96 | 0.04 | 0.36 | 0.23 | 4.42 | 41.867654 | 9.77E-11 |
| VI | Euarchontoglires | - | - | - | - | - | - | - | - | - |
| VII | Microcebus | 0.23 | 0.27 | 0.79 | 0.21 | 29.05 | 99.00 | 2.14 | 51.540866 | 7.01E-13 |
| VII | Cheirogaleidae | 0.38 | 0.45 | 0.78 | 0.22 | 3.50 | 10.37 | 1.78 | 44.981404 | 1.99E-11 |
| VII | Lemuriformes | 0.37 | 0.43 | 0.76 | 0.24 | 1.90 | 5.96 | 1.59 | 35.581718 | 2.45E-09 |
| VII | Strepsirrhini | 0.41 | 0.35 | 0.85 | 0.15 | 0.62 | 0.75 | 2.00 | 44.861122 | 2.12E-11 |
| VII | Primates | - | - | - | - | - | - | - | - | - |
| VII | Euarchontoglires | - | - | - | - | - | - | - | - | - |
| VIII | Microcebus | 0.02 | 0.02 | 0.76 | 0.24 | 0.75 | 2.24 | 2.67 | 34.657132 | 3.93E-09 |
| VIII | Cheirogaleidae | 0.04 | 0.02 | 0.82 | 0.18 | 0.56 | 0.69 | 2.65 | 47.580534 | 5.28E-12 |
| VIII | Lemuriformes | - | - | - | - | - | - | - | - | - |
| VIII | Strepsirrhini | 0.53 | 0.37 | 0.83 | 0.17 | 1.09 | 1.03 | 2.67 | 87.56152 | 8.17E-21 |
| VIII | Primates | - | - | - | - | - | - | - | - | - |
| VIII | Euarchontoglires | 0.63 | 0.38 | 0.82 | 0.18 | 1.20 | 1.03 | 2.39 | 79.258242 | 5.45E-19 |
| IX | Microcebus | 0.28 | 0.36 | 0.82 | 0.18 | 0.43 | 0.87 | 2.73 | 289.51926 | 6.33E-65 |
| IX | Cheirogaleidae | 0.32 | 0.36 | 0.81 | 0.19 | 0.46 | 0.79 | 2.63 | 360.772864 | 1.91E-80 |
| IX | Lemuriformes | - | - | - | - | - | - | - | - | - |
| IX | Strepsirrhini | 0.34 | 0.38 | 0.80 | 0.20 | 0.51 | 0.85 | 2.62 | 376.86275 | 6.00E-84 |
| IX | Primates | - | - | - | - | - | - | - | - | - |
| IX | Euarchontoglires | - | - | - | - | - | - | - | - | - |

**Table S8 – P-values from chi-square tests for biases of pervasive selection occurring in transmembrane or loop domains across subfamilies.** Asterisks (*) indicate a p-value less than 0.05 given a null distribution of $X^2_1$.

| Subfamily | Filter | Chi Square | p-value | significant | Proportion TM | Proportion Loop | Expected Proportion |
|---|---|---|---|---|---|---|---|
| I | Microcebus | 1.61 | 0.20455 | NS | 0.129 | 0.094 | 0.104 |
| I | Cheirogaleidae | 7.03 | 0.0080016 | * | 0.158 | 0.084 | 0.105 |
| I | Lemuriformes | 4.59 | 0.032105 | * | 0.164 | 0.101 | 0.119 |
| I | Strepsirrhini | 2.20 | 0.13794 | NS | 0.146 | 0.103 | 0.116 |
| I | Primates | 3.83 | 0.050241 | NS | 0.152 | 0.096 | 0.112 |
| I | Euarchontoglires | 2.78 | 0.095474 | NS | 0.140 | 0.094 | 0.107 |
| II | Microcebus | 4.28 | 0.038654 | * | 0.140 | 0.084 | 0.100 |
| II | Cheirogaleidae | 4.59 | 0.032105 | * | 0.164 | 0.101 | 0.119 |
| II | Lemuriformes | 0.75 | 0.38538 | NS | 0.146 | 0.120 | 0.128 |
| II | Strepsirrhini | 0.38 | 0.53519 | NS | 0.146 | 0.127 | 0.133 |
| II | Euarchontoglires | 1.76 | 0.18491 | NS | 0.158 | 0.118 | 0.129 |
| III | Microcebus | 0.00 | 0.99873 | NS | 0.094 | 0.094 | 0.094 |
| III | Cheirogaleidae | 1.96 | 0.16169 | NS | 0.064 | 0.101 | 0.090 |
| III | Lemuriformes | 1.36 | 0.24414 | NS | 0.070 | 0.101 | 0.092 |
| III | Strepsirrhini | 2.21 | 0.13691 | NS | 0.076 | 0.118 | 0.105 |
| III | Primates | 5.49 | 0.01911 | * | 0.070 | 0.139 | 0.119 |
| III | Euarchontoglires | 3.44 | 0.063462 | NS | 0.070 | 0.122 | 0.107 |
| IV | Microcebus | 0.30 | 0.58325 | NS | 0.088 | 0.074 | 0.078 |
| IV | Cheirogaleidae | 1.73 | 0.18881 | NS | 0.041 | 0.070 | 0.061 |
| IV | Strepsirrhini | 1.07 | 0.30198 | NS | 0.047 | 0.070 | 0.063 |
| IV | Euarchontoglires | 2.87 | 0.090478 | NS | 0.035 | 0.072 | 0.061 |
| V | Microcebus | 8.55 | 0.0034553 | * | 0.170 | 0.086 | 0.111 |
| V | Cheirogaleidae | 2.14 | 0.14336 | NS | 0.129 | 0.089 | 0.100 |
| V | Lemuriformes | 3.12 | 0.077566 | NS | 0.140 | 0.091 | 0.105 |
| V | Strepsirrhini | 3.48 | 0.062259 | NS | 0.164 | 0.108 | 0.124 |
| V | Primates | 2.25 | 0.13356 | NS | 0.158 | 0.113 | 0.126 |
| VI | Microcebus | 12.37 | 0.00043689 | * | 0.199 | 0.094 | 0.124 |
| VI | Cheirogaleidae | 10.74 | 0.0010499 | * | 0.164 | 0.074 | 0.100 |
| VI | Lemuriformes | 6.49 | 0.010834 | * | 0.158 | 0.086 | 0.107 |
| VI | Primates | 0.79 | 0.37518 | NS | 0.094 | 0.072 | 0.078 |
| VII | Microcebus | 0.91 | 0.33983 | NS | 0.082 | 0.108 | 0.100 |
| VII | Cheirogaleidae | 2.74 | 0.097887 | NS | 0.047 | 0.086 | 0.075 |
| VII | Lemuriformes | 1.26 | 0.26123 | NS | 0.041 | 0.065 | 0.058 |
| VII | Strepsirrhini | 1.27 | 0.2601 | NS | 0.105 | 0.077 | 0.085 |
| VIII | Microcebus | 18.49 | 0.000017052 | * | 0.240 | 0.103 | 0.143 |
| VIII | Cheirogaleidae | 3.48 | 0.062278 | NS | 0.140 | 0.089 | 0.104 |
| VIII | Strepsirrhini | 0.06 | 0.79916 | NS | 0.105 | 0.098 | 0.100 |
| VIII | Euarchontoglires | 3.83 | 0.050361 | NS | 0.170 | 0.110 | 0.128 |
| IX | Microcebus | 1.23 | 0.26704 | NS | 0.140 | 0.108 | 0.117 |
| IX | Cheirogaleidae | 1.54 | 0.21492 | NS | 0.158 | 0.120 | 0.131 |
| IX | Strepsirrhini | 2.55 | 0.11038 | NS | 0.164 | 0.115 | 0.129 |

**Table S9 – P-values from Fisher exact tests for biases of pervasive selection occurring within individual transmembrane domains with respect to the other six.** Asterisks (*) indicate a p-value less than 0.05 given a null distribution of $X^2_1$.

| Subfamily | Filter | TM Domain | p-value | Odds Ratio | Significant |
|---|---|---|---|---|---|
| I | Microcebus | TM1 | 0.047081412 | 0 | * |
| I | Microcebus | TM2 | 0.327206822 | 1.888790677 | NS |
| I | Microcebus | TM3 | 0.046370661 | 0 | * |
| I | Microcebus | TM4 | 0.350786439 | 1.691888237 | NS |
| I | Microcebus | TM5 | 0.168022974 | 2.270072172 | NS |
| I | Microcebus | TM6 | 1 | 0.962604963 | NS |
| I | Microcebus | TM7 | 0.518523455 | 1.430020846 | NS |
| I | Cheirogaleidae | TM1 | 0.015437577 | 0 | * |
| I | Cheirogaleidae | TM2 | 0.238366712 | 1.871747847 | NS |
| I | Cheirogaleidae | TM3 | 0.015853999 | 0 | * |
| I | Cheirogaleidae | TM4 | 0.773305955 | 1.258532828 | NS |
| I | Cheirogaleidae | TM5 | 0.052979029 | 2.985011333 | NS |
| I | Cheirogaleidae | TM6 | 0.770446263 | 0.733389151 | NS |
| I | Cheirogaleidae | TM7 | 0.224019595 | 1.990512011 | NS |
| I | Lemuriformes | TM1 | 0.37819732 | 0.403052967 | NS |
| I | Lemuriformes | TM2 | 0.566455676 | 1.334528628 | NS |
| I | Lemuriformes | TM3 | 0.015394726 | 0 | * |
| I | Lemuriformes | TM4 | 1 | 0.870255842 | NS |
| I | Lemuriformes | TM5 | 0.012312593 | 3.648209397 | * |
| I | Lemuriformes | TM6 | 0.769383514 | 0.698505446 | NS |
| I | Lemuriformes | TM7 | 0.236434174 | 1.885818042 | NS |
| I | Strepsirrhini | TM1 | 0.538616846 | 0.466706747 | NS |
| I | Strepsirrhini | TM2 | 0.373753578 | 1.57033329 | NS |
| I | Strepsirrhini | TM3 | 0.026937586 | 0 | * |
| I | Strepsirrhini | TM4 | 1 | 1.018523926 | NS |
| I | Strepsirrhini | TM5 | 0.100137263 | 2.547872991 | NS |
| I | Strepsirrhini | TM6 | 0.534978296 | 0.491779836 | NS |
| I | Strepsirrhini | TM7 | 0.127287641 | 2.232754188 | NS |
| I | Primates | TM1 | 0.375548307 | 0.443717441 | NS |
| I | Primates | TM2 | 0.545287361 | 1.484294378 | NS |
| I | Primates | TM3 | 0.027425054 | 0 | * |
| I | Primates | TM4 | 1 | 0.964610109 | NS |
| I | Primates | TM5 | 0.11070173 | 2.403325351 | NS |
| I | Primates | TM6 | 1 | 0.771294249 | NS |
| I | Primates | TM7 | 0.214487747 | 2.105615452 | NS |
| I | Euarchontoglires | TM1 | 0.534978296 | 0.491779836 | NS |
| I | Euarchontoglires | TM2 | 0.35517097 | 1.665369573 | NS |
| I | Euarchontoglires | TM3 | 0.027573458 | 0 | * |
| I | Euarchontoglires | TM4 | 0.770446263 | 0.733389151 | NS |
| I | Euarchontoglires | TM5 | 0.201724482 | 2.002593945 | NS |
| I | Euarchontoglires | TM6 | 1 | 0.857886831 | NS |
| I | Euarchontoglires | TM7 | 0.113050798 | 2.37389745 | NS |
| II | Microcebus | TM1 | 0.026937586 | 0 | * |
| II | Microcebus | TM2 | 0.127287641 | 2.232754188 | NS |
| II | Microcebus | TM3 | 0.204987066 | 0.235719989 | NS |
| II | Microcebus | TM4 | 0.068761972 | 2.596272684 | NS |
| II | Microcebus | TM5 | 1 | 0.962604963 | NS |

| | | | | | |
|---|---|---|---|---|---|
| II | Microcebus | TM6 | 0.534672633 | 0.518225846 | NS |
| II | Microcebus | TM7 | 0.340135491 | 1.765947801 | NS |
| II | Cheirogaleidae | TM1 | 0.082470726 | 0.184794963 | NS |
| II | Cheirogaleidae | TM2 | 0.566455676 | 1.334528628 | NS |
| II | Cheirogaleidae | TM3 | 0.132403821 | 0.194423813 | NS |
| II | Cheirogaleidae | TM4 | 0.051369144 | 2.592435357 | NS |
| II | Cheirogaleidae | TM5 | 0.762246155 | 1.156418332 | NS |
| II | Cheirogaleidae | TM6 | 0.374721126 | 0.424771163 | NS |
| II | Cheirogaleidae | TM7 | 0.078514219 | 2.454600228 | NS |
| II | Lemuriformes | TM1 | 0.131505024 | 0.213053137 | NS |
| II | Lemuriformes | TM2 | 0.373753578 | 1.57033329 | NS |
| II | Lemuriformes | TM3 | 0.208070643 | 0.224107739 | NS |
| II | Lemuriformes | TM4 | 0.032325234 | 3.118420015 | * |
| II | Lemuriformes | TM5 | 1 | 0.911965769 | NS |
| II | Lemuriformes | TM6 | 1 | 0.812630849 | NS |
| II | Lemuriformes | TM7 | 0.757379981 | 1.198660987 | NS |
| II | Strepsirrhini | TM1 | 0.027496606 | 0 | * |
| II | Strepsirrhini | TM2 | 0.373753578 | 1.57033329 | NS |
| II | Strepsirrhini | TM3 | 0.208070643 | 0.224107739 | NS |
| II | Strepsirrhini | TM4 | 0.032325234 | 3.118420015 | * |
| II | Strepsirrhini | TM5 | 0.534897443 | 1.351899921 | NS |
| II | Strepsirrhini | TM6 | 1 | 0.812630849 | NS |
| II | Strepsirrhini | TM7 | 0.757379981 | 1.198660987 | NS |
| II | Euarchontoglires | TM1 | 0.133011436 | 0.193490944 | NS |
| II | Euarchontoglires | TM2 | 1 | 1.018523926 | NS |
| II | Euarchontoglires | TM3 | 0.130604584 | 0.203558588 | NS |
| II | Euarchontoglires | TM4 | 0.017241668 | 3.466662637 | * |
| II | Euarchontoglires | TM5 | 0.755453188 | 1.215920634 | NS |
| II | Euarchontoglires | TM6 | 0.770446263 | 0.733389151 | NS |
| II | Euarchontoglires | TM7 | 0.544536236 | 1.491346538 | NS |
| III | Microcebus | TM1 | 0.132623631 | 0 | NS |
| III | Microcebus | TM2 | 0.062506624 | 3.041280203 | NS |
| III | Microcebus | TM3 | 0.703131472 | 0.384202726 | NS |
| III | Microcebus | TM4 | 0.28649934 | 1.904519216 | NS |
| III | Microcebus | TM5 | 0.43750096 | 1.646211521 | NS |
| III | Microcebus | TM6 | 0.131800652 | 0 | NS |
| III | Microcebus | TM7 | 0.474241535 | 1.468808022 | NS |
| III | Cheirogaleidae | TM1 | 0.3706493 | 0 | NS |
| III | Cheirogaleidae | TM2 | 0.057803898 | 3.740405573 | NS |
| III | Cheirogaleidae | TM3 | 0.366437303 | 0 | NS |
| III | Cheirogaleidae | TM4 | 0.383166655 | 2.113482238 | NS |
| III | Cheirogaleidae | TM5 | 0.362688509 | 0 | NS |
| III | Cheirogaleidae | TM6 | 0.366437303 | 0 | NS |
| III | Cheirogaleidae | TM7 | 0.050362257 | 3.952920818 | NS |
| III | Lemuriformes | TM1 | 0.688908855 | 1.181355125 | NS |
| III | Lemuriformes | TM2 | 0.386967449 | 2.064863153 | NS |
| III | Lemuriformes | TM3 | 0.220744063 | 0 | NS |
| III | Lemuriformes | TM4 | 1 | 1.071568415 | NS |
| III | Lemuriformes | TM5 | 0.368068411 | 0 | NS |
| III | Lemuriformes | TM6 | 1 | 0.539156958 | NS |
| III | Lemuriformes | TM7 | 0.068309402 | 3.439421615 | NS |
| III | Strepsirrhini | TM1 | 1 | 1.066792429 | NS |
| III | Strepsirrhini | TM2 | 0.408633341 | 1.846472576 | NS |
| III | Strepsirrhini | TM3 | 0.696484148 | 0.490770557 | NS |

| | | | | | |
|---|---|---|---|---|---|
| III | Strepsirrhini | TM4 | 1 | 0.967451106 | NS |
| III | Strepsirrhini | TM5 | 0.222857902 | 0 | NS |
| III | Strepsirrhini | TM6 | 0.696484148 | 0.490770557 | NS |
| III | Strepsirrhini | TM7 | 0.089228925 | 3.039311542 | NS |
| III | Primates | TM1 | 0.688908855 | 1.181355125 | NS |
| III | Primates | TM2 | 0.386967449 | 2.064863153 | NS |
| III | Primates | TM3 | 0.220744063 | 0 | NS |
| III | Primates | TM4 | 1 | 1.071568415 | NS |
| III | Primates | TM5 | 0.368068411 | 0 | NS |
| III | Primates | TM6 | 1 | 0.539156958 | NS |
| III | Primates | TM7 | 0.068309402 | 3.439421615 | NS |
| III | Euarchontoglires | TM1 | 0.688908855 | 1.181355125 | NS |
| III | Euarchontoglires | TM2 | 0.688908855 | 1.181355125 | NS |
| III | Euarchontoglires | TM3 | 0.220744063 | 0 | NS |
| III | Euarchontoglires | TM4 | 1 | 1.071568415 | NS |
| III | Euarchontoglires | TM5 | 1 | 0.598927814 | NS |
| III | Euarchontoglires | TM6 | 1 | 0.539156958 | NS |
| III | Euarchontoglires | TM7 | 0.068309402 | 3.439421615 | NS |
| IV | Microcebus | TM1 | 1 | 0.8902204 | NS |
| IV | Microcebus | TM2 | 0.463155063 | 1.518511301 | NS |
| IV | Microcebus | TM3 | 0.133423622 | 0 | NS |
| IV | Microcebus | TM4 | 0.261417097 | 2.091674871 | NS |
| IV | Microcebus | TM5 | 0.414752837 | 1.795198147 | NS |
| IV | Microcebus | TM6 | 0.445899998 | 1.602035114 | NS |
| IV | Microcebus | TM7 | 0.133423622 | 0 | NS |
| IV | Cheirogaleidae | TM1 | 0.595409788 | 0 | NS |
| IV | Cheirogaleidae | TM2 | 0.271737619 | 2.435450173 | NS |
| IV | Cheirogaleidae | TM3 | 0.595065456 | 0 | NS |
| IV | Cheirogaleidae | TM4 | 0.079667159 | 4.319936432 | NS |
| IV | Cheirogaleidae | TM5 | 0.596893438 | 0 | NS |
| IV | Cheirogaleidae | TM6 | 0.255209678 | 2.562652437 | NS |
| IV | Cheirogaleidae | TM7 | 0.595065456 | 0 | NS |
| IV | Strepsirrhini | TM1 | 0.605277842 | 0 | NS |
| IV | Strepsirrhini | TM2 | 0.330945942 | 2.018641851 | NS |
| IV | Strepsirrhini | TM3 | 0.602120065 | 0 | NS |
| IV | Strepsirrhini | TM4 | 0.114025886 | 3.439722901 | NS |
| IV | Strepsirrhini | TM5 | 1 | 0.966175688 | NS |
| IV | Strepsirrhini | TM6 | 0.311999279 | 2.124420529 | NS |
| IV | Strepsirrhini | TM7 | 0.602120065 | 0 | NS |
| IV | Euarchontoglires | TM1 | 0.594180031 | 0 | NS |
| IV | Euarchontoglires | TM2 | 0.212709259 | 3.058710484 | NS |
| IV | Euarchontoglires | TM3 | 0.596694971 | 0 | NS |
| IV | Euarchontoglires | TM4 | 0.240662338 | 2.777649491 | NS |
| IV | Euarchontoglires | TM5 | 1 | 0 | NS |
| IV | Euarchontoglires | TM6 | 0.198991401 | 3.218257231 | NS |
| IV | Euarchontoglires | TM7 | 0.596694971 | 0 | NS |
| V | Microcebus | TM1 | 0.081779029 | 0.176719194 | NS |
| V | Microcebus | TM2 | 0.772657504 | 1.268952262 | NS |
| V | Microcebus | TM3 | 0.769899579 | 0.666295903 | NS |
| V | Microcebus | TM4 | 0.576357431 | 0.568945774 | NS |
| V | Microcebus | TM5 | 1 | 0.748158743 | NS |
| V | Microcebus | TM6 | 0.136830992 | 2.325507614 | NS |
| V | Microcebus | TM7 | 0.136830992 | 2.325507614 | NS |
| V | Cheirogaleidae | TM1 | 0.745603833 | 0.549484408 | NS |

| | | | | | |
|---|---|---|---|---|---|
| V | Cheirogaleidae | TM2 | 0.745603833 | 0.549484408 | NS |
| V | Cheirogaleidae | TM3 | 0.319720061 | 0.262255596 | NS |
| V | Cheirogaleidae | TM4 | 1 | 0.823266298 | NS |
| V | Cheirogaleidae | TM5 | 1 | 1.079848882 | NS |
| V | Cheirogaleidae | TM6 | 0.018212625 | 3.586773086 | * |
| V | Cheirogaleidae | TM7 | 0.518523455 | 1.430020846 | NS |
| V | Lemuriformes | TM1 | 0.534978296 | 0.491779836 | NS |
| V | Lemuriformes | TM2 | 0.534978296 | 0.491779836 | NS |
| V | Lemuriformes | TM3 | 0.204987066 | 0.235719989 | NS |
| V | Lemuriformes | TM4 | 1 | 1.077788285 | NS |
| V | Lemuriformes | TM5 | 0.518523455 | 1.430020846 | NS |
| V | Lemuriformes | TM6 | 0.113050798 | 2.37389745 | NS |
| V | Lemuriformes | TM7 | 0.340135491 | 1.765947801 | NS |
| V | Strepsirrhini | TM1 | 0.37819732 | 0.403052967 | NS |
| V | Strepsirrhini | TM2 | 0.770279982 | 0.661473655 | NS |
| V | Strepsirrhini | TM3 | 0.132403821 | 0.194423813 | NS |
| V | Strepsirrhini | TM4 | 0.397214238 | 1.579696842 | NS |
| V | Strepsirrhini | TM5 | 0.366294559 | 1.606216512 | NS |
| V | Strepsirrhini | TM6 | 0.236434174 | 1.885818042 | NS |
| V | Strepsirrhini | TM7 | 0.553262922 | 1.415615117 | NS |
| V | Primates | TM1 | 0.375093575 | 0.422562566 | NS |
| V | Primates | TM2 | 0.769479564 | 0.694556309 | NS |
| V | Primates | TM3 | 0.375584685 | 0.445325224 | NS |
| V | Primates | TM4 | 0.386227939 | 1.667814299 | NS |
| V | Primates | TM5 | 0.350786439 | 1.691888237 | NS |
| V | Primates | TM6 | 1 | 1.077788285 | NS |
| V | Primates | TM7 | 0.544536236 | 1.491346538 | NS |
| VI | Microcebus | TM1 | 0.283220107 | 1.707936693 | NS |
| VI | Microcebus | TM2 | 1 | 1.008570113 | NS |
| VI | Microcebus | TM3 | 0.788624778 | 0.781086935 | NS |
| VI | Microcebus | TM4 | 0.295855607 | 0.457404965 | NS |
| VI | Microcebus | TM5 | 1 | 0.882115778 | NS |
| VI | Microcebus | TM6 | 0.580772477 | 1.413564331 | NS |
| VI | Microcebus | TM7 | 1 | 1.070353165 | NS |
| VI | Cheirogaleidae | TM1 | 1 | 0.96841978 | NS |
| VI | Cheirogaleidae | TM2 | 0.254471194 | 1.772990215 | NS |
| VI | Cheirogaleidae | TM3 | 0.374721126 | 0.424771163 | NS |
| VI | Cheirogaleidae | TM4 | 0.257051389 | 0.364763604 | NS |
| VI | Cheirogaleidae | TM5 | 0.211414911 | 2.153083432 | NS |
| VI | Cheirogaleidae | TM6 | 0.374721126 | 0.424771163 | NS |
| VI | Cheirogaleidae | TM7 | 0.236434174 | 1.885818042 | NS |
| VI | Lemuriformes | TM1 | 1 | 1.018523926 | NS |
| VI | Lemuriformes | TM2 | 0.238366712 | 1.871747847 | NS |
| VI | Lemuriformes | TM3 | 0.375584685 | 0.445325224 | NS |
| VI | Lemuriformes | TM4 | 0.082072831 | 0.175675494 | NS |
| VI | Lemuriformes | TM5 | 0.123226923 | 2.272306493 | NS |
| VI | Lemuriformes | TM6 | 0.375584685 | 0.445325224 | NS |
| VI | Lemuriformes | TM7 | 0.224019595 | 1.990512011 | NS |
| VI | Primates | TM1 | 1 | 0.820776155 | NS |
| VI | Primates | TM2 | 0.708048549 | 1.392122726 | NS |
| VI | Primates | TM3 | 0.131800652 | 0 | NS |
| VI | Primates | TM4 | 0.472506477 | 0.332303194 | NS |
| VI | Primates | TM5 | 0.43750096 | 1.646211521 | NS |
| VI | Primates | TM6 | 0.474241535 | 1.468808022 | NS |

| | | | | | |
|---|---|---|---|---|---|
| VI | Primates | TM7 | 0.246790026 | 2.236695006 | NS |
| VII | Microcebus | TM1 | 0.434396622 | 1.667722458 | NS |
| VII | Microcebus | TM2 | 0.127739906 | 2.571683981 | NS |
| VII | Microcebus | TM3 | 0.695203491 | 0.449806551 | NS |
| VII | Microcebus | TM4 | 0.129942405 | 0 | NS |
| VII | Microcebus | TM5 | 1 | 0.499799387 | NS |
| VII | Microcebus | TM6 | 0.695203491 | 0.449806551 | NS |
| VII | Microcebus | TM7 | 0.112981852 | 2.718694361 | NS |
| VII | Cheirogaleidae | TM1 | 0.330945942 | 2.018641851 | NS |
| VII | Cheirogaleidae | TM2 | 0.330945942 | 2.018641851 | NS |
| VII | Cheirogaleidae | TM3 | 1 | 0.870244336 | NS |
| VII | Cheirogaleidae | TM4 | 0.359019693 | 0 | NS |
| VII | Cheirogaleidae | TM5 | 0.59829799 | 0 | NS |
| VII | Cheirogaleidae | TM6 | 0.602120065 | 0 | NS |
| VII | Cheirogaleidae | TM7 | 0.084840876 | 4.008202584 | NS |
| VII | Lemuriformes | TM1 | 1 | 0.972377558 | NS |
| VII | Lemuriformes | TM2 | 0.271737619 | 2.435450173 | NS |
| VII | Lemuriformes | TM3 | 1 | 1.021611706 | NS |
| VII | Lemuriformes | TM4 | 0.598337668 | 0 | NS |
| VII | Lemuriformes | TM5 | 0.596893438 | 0 | NS |
| VII | Lemuriformes | TM6 | 0.595065456 | 0 | NS |
| VII | Lemuriformes | TM7 | 0.05844571 | 5.03106327 | NS |
| VII | Strepsirrhini | TM1 | 1 | 0.707828145 | NS |
| VII | Strepsirrhini | TM2 | 0.029149157 | 3.490816447 | * |
| VII | Strepsirrhini | TM3 | 0.47415965 | 0.334012056 | NS |
| VII | Strepsirrhini | TM4 | 0.7416784 | 0.641487918 | NS |
| VII | Strepsirrhini | TM5 | 0.133344799 | 0 | NS |
| VII | Strepsirrhini | TM6 | 0.47415965 | 0.334012056 | NS |
| VII | Strepsirrhini | TM7 | 0.023734098 | 3.709789809 | * |
| VIII | Microcebus | TM1 | 0.616684102 | 1.279123458 | NS |
| VIII | Microcebus | TM2 | 0.616684102 | 1.279123458 | NS |
| VIII | Microcebus | TM3 | 0.069566163 | 0.253252595 | NS |
| VIII | Microcebus | TM4 | 0.624807196 | 0.683241207 | NS |
| VIII | Microcebus | TM5 | 0.180113971 | 1.999439189 | NS |
| VIII | Microcebus | TM6 | 0.448031274 | 0.596225729 | NS |
| VIII | Microcebus | TM7 | 0.301674295 | 1.721145364 | NS |
| VIII | Cheirogaleidae | TM1 | 0.208070643 | 0.224107739 | NS |
| VIII | Cheirogaleidae | TM2 | 0.054503626 | 2.926994208 | NS |
| VIII | Cheirogaleidae | TM3 | 0.204987066 | 0.235719989 | NS |
| VIII | Cheirogaleidae | TM4 | 0.224019595 | 1.990512011 | NS |
| VIII | Cheirogaleidae | TM5 | 1 | 0.962604963 | NS |
| VIII | Cheirogaleidae | TM6 | 1 | 0.857886831 | NS |
| VIII | Cheirogaleidae | TM7 | 1 | 0.857886831 | NS |
| VIII | Strepsirrhini | TM1 | 0.730341801 | 1.18963246 | NS |
| VIII | Strepsirrhini | TM2 | 0.730341801 | 1.18963246 | NS |
| VIII | Strepsirrhini | TM3 | 0.079868828 | 0 | NS |
| VIII | Strepsirrhini | TM4 | 1 | 1.074548729 | NS |
| VIII | Strepsirrhini | TM5 | 1 | 0.832101809 | NS |
| VIII | Strepsirrhini | TM6 | 1 | 0.745502601 | NS |
| VIII | Strepsirrhini | TM7 | 0.141100642 | 2.692030388 | NS |
| VIII | Euarchontoglires | TM1 | 0.145849317 | 2.179996348 | NS |
| VIII | Euarchontoglires | TM2 | 0.772657504 | 1.268952262 | NS |
| VIII | Euarchontoglires | TM3 | 0.01561596 | 0 | * |
| VIII | Euarchontoglires | TM4 | 1 | 0.828708138 | NS |

| | | | | | |
|---|---|---|---|---|---|
| VIII | Euarchontoglires | TM5 | 1 | 0.748158743 | NS |
| VIII | Euarchontoglires | TM6 | 0.56424206 | 1.346156689 | NS |
| VIII | Euarchontoglires | TM7 | 0.56424206 | 1.346156689 | NS |
| IX | Microcebus | TM1 | 0.757379981 | 1.198660987 | NS |
| IX | Microcebus | TM2 | 0.757379981 | 1.198660987 | NS |
| IX | Microcebus | TM3 | 0.027573458 | 0 | * |
| IX | Microcebus | TM4 | 0.544536236 | 1.491346538 | NS |
| IX | Microcebus | TM5 | 0.518523455 | 1.430020846 | NS |
| IX | Microcebus | TM6 | 1 | 0.857886831 | NS |
| IX | Microcebus | TM7 | 0.750910114 | 1.268114233 | NS |
| IX | Cheirogaleidae | TM1 | 1 | 1.018523926 | NS |
| IX | Cheirogaleidae | TM2 | 0.554676431 | 1.406028791 | NS |
| IX | Cheirogaleidae | TM3 | 0.015853999 | 0 | * |
| IX | Cheirogaleidae | TM4 | 0.386227939 | 1.667814299 | NS |
| IX | Cheirogaleidae | TM5 | 0.755453188 | 1.215920634 | NS |
| IX | Cheirogaleidae | TM6 | 0.375584685 | 0.445325224 | NS |
| IX | Cheirogaleidae | TM7 | 0.224019595 | 1.990512011 | NS |
| IX | Strepsirrhini | TM1 | 1 | 0.96841978 | NS |
| IX | Strepsirrhini | TM2 | 0.566455676 | 1.334528628 | NS |
| IX | Strepsirrhini | TM3 | 0.015394726 | 0 | * |
| IX | Strepsirrhini | TM4 | 0.397214238 | 1.579696842 | NS |
| IX | Strepsirrhini | TM5 | 0.762246155 | 1.156418332 | NS |
| IX | Strepsirrhini | TM6 | 0.374721126 | 0.424771163 | NS |
| IX | Strepsirrhini | TM7 | 0.078514219 | 2.454600228 | NS |

**Table S10 – P-values from Fisher exact tests for biases of pervasive selection occurring within individual loop domains with respect to the other seven.** Asterisks (*) indicate a p-value less than 0.05 given a null distribution of $X^2_1$.

| Subfamily | Filter | Loop Domain | p-value | Odds Ratio | Significant |
|---|---|---|---|---|---|
| I | Microcebus | L1 | 0.082484061 | 0.505010614 | NS |
| I | Microcebus | L2 | 0.65347923 | 1.404098116 | NS |
| I | Microcebus | L3 | 1 | 1.08090091 | NS |
| I | Microcebus | L4 | 0.418383639 | 1.766631716 | NS |
| I | Microcebus | L5 | 0.066938401 | 2.529891805 | NS |
| I | Microcebus | L6 | 0.349431592 | 0.275664583 | NS |
| I | Microcebus | L7 | 0.100528381 | 5.019019417 | NS |
| I | Microcebus | L8 | 0.858584504 | 1.061828028 | NS |
| I | Cheirogaleidae | L1 | 0.100309834 | 0.501536589 | NS |
| I | Cheirogaleidae | L2 | 0.142262217 | 2.652063918 | NS |
| I | Cheirogaleidae | L3 | 0.680115251 | 1.224943211 | NS |
| I | Cheirogaleidae | L4 | 0.230378385 | 2.008510523 | NS |
| I | Cheirogaleidae | L5 | 0.220770676 | 1.819613375 | NS |
| I | Cheirogaleidae | L6 | 0.340393596 | 0.311628939 | NS |
| I | Cheirogaleidae | L7 | 0.082901559 | 5.681672485 | NS |
| I | Cheirogaleidae | L8 | 1 | 0.965188606 | NS |
| I | Lemuriformes | L1 | 0.006175581 | 0.32522281 | * |
| I | Lemuriformes | L2 | 0.669618776 | 1.288421608 | NS |
| I | Lemuriformes | L3 | 0.440017021 | 1.617679908 | NS |
| I | Lemuriformes | L4 | 0.440017021 | 1.617679908 | NS |
| I | Lemuriformes | L5 | 0.001689232 | 4.007518158 | * |
| I | Lemuriformes | L6 | 0.23189318 | 0.253293195 | NS |
| I | Lemuriformes | L7 | 0.01537091 | 9.434022575 | * |
| I | Lemuriformes | L8 | 1 | 0.941140228 | NS |
| I | Strepsirrhini | L1 | 0.0040765 | 0.314843236 | * |
| I | Strepsirrhini | L2 | 0.675282455 | 1.253614376 | NS |
| I | Strepsirrhini | L3 | 0.447988558 | 1.572978068 | NS |
| I | Strepsirrhini | L4 | 0.139524078 | 2.288784672 | NS |
| I | Strepsirrhini | L5 | 0.007894057 | 3.257729612 | * |
| I | Strepsirrhini | L6 | 0.233936619 | 0.246545767 | NS |
| I | Strepsirrhini | L7 | 0.016428172 | 9.176206534 | * |
| I | Strepsirrhini | L8 | 1 | 1.021801219 | NS |
| I | Primates | L1 | 0.009464965 | 0.347866743 | * |
| I | Primates | L2 | 0.658697491 | 1.363516019 | NS |
| I | Primates | L3 | 0.42517886 | 1.71430691 | NS |
| I | Primates | L4 | 0.42517886 | 1.71430691 | NS |
| I | Primates | L5 | 0.070709689 | 2.446528569 | NS |

| | | | | | |
|---|---|---|---|---|---|
| I | Primates | L6 | 0.231070034 | 0.267825327 | NS |
| I | Primates | L7 | 0.013382585 | 9.990971184 | * |
| I | Primates | L8 | 0.722743734 | 1.152792367 | NS |
| I | Euarchontoglires | L1 | 0.036145742 | 0.429669761 | * |
| I | Euarchontoglires | L2 | 0.180583096 | 2.33319195 | NS |
| I | Euarchontoglires | L3 | 1 | 1.08090091 | NS |
| I | Euarchontoglires | L4 | 0.418383639 | 1.766631716 | NS |
| I | Euarchontoglires | L5 | 0.066938401 | 2.529891805 | NS |
| I | Euarchontoglires | L6 | 0.349431592 | 0.275664583 | NS |
| I | Euarchontoglires | L7 | 0.012450545 | 10.29238498 | * |
| I | Euarchontoglires | L8 | 1 | 0.932753556 | NS |
| II | Microcebus | L1 | 0.000181375 | 0.150359727 | * |
| II | Microcebus | L2 | 1 | 0.720111652 | NS |
| II | Microcebus | L3 | 0.019180467 | 4.055442254 | * |
| II | Microcebus | L4 | 0.680115251 | 1.224943211 | NS |
| II | Microcebus | L5 | 7.91E-06 | 7.384573173 | * |
| II | Microcebus | L6 | 1 | 1.061353078 | NS |
| II | Microcebus | L7 | 0.082901559 | 5.681672485 | NS |
| II | Microcebus | L8 | 0.131093132 | 0.502705274 | NS |
| II | Cheirogaleidae | L1 | 6.96E-05 | 0.165629049 | * |
| II | Cheirogaleidae | L2 | 0.669618776 | 1.288421608 | NS |
| II | Cheirogaleidae | L3 | 0.001834361 | 5.529342217 | * |
| II | Cheirogaleidae | L4 | 0.130644377 | 2.355228794 | NS |
| II | Cheirogaleidae | L5 | 1.09E-05 | 6.505272671 | * |
| II | Cheirogaleidae | L6 | 1 | 0.853905509 | NS |
| II | Cheirogaleidae | L7 | 0.473249224 | 1.801677176 | NS |
| II | Cheirogaleidae | L8 | 0.057213699 | 0.465738282 | NS |
| II | Lemuriformes | L1 | 0.000118096 | 0.211798163 | * |
| II | Lemuriformes | L2 | 1 | 1.050479644 | NS |
| II | Lemuriformes | L3 | 0.005340828 | 4.408163821 | * |
| II | Lemuriformes | L4 | 0.281223854 | 1.90403355 | NS |
| II | Lemuriformes | L5 | 2.62E-06 | 6.668636879 | * |
| II | Lemuriformes | L6 | 1 | 0.976863521 | NS |
| II | Lemuriformes | L7 | 0.15436895 | 3.762584595 | NS |
| II | Lemuriformes | L8 | 0.053579218 | 0.490670961 | NS |
| II | Strepsirrhini | L1 | 0.000112927 | 0.235844523 | * |
| II | Strepsirrhini | L2 | 1 | 0.980437182 | NS |
| II | Strepsirrhini | L3 | 0.007525726 | 4.087777714 | * |
| II | Strepsirrhini | L4 | 0.156884606 | 2.41678507 | NS |
| II | Strepsirrhini | L5 | 6.12E-06 | 6.093738222 | * |
| II | Strepsirrhini | L6 | 1 | 0.909065701 | NS |
| II | Strepsirrhini | L7 | 0.170168609 | 3.513114885 | NS |
| II | Strepsirrhini | L8 | 0.060302556 | 0.513906941 | NS |

| | | | | | |
|---|---|---|---|---|---|
| II | Euarchontoglires | L1 | 0.002315255 | 0.311293343 | * |
| II | Euarchontoglires | L2 | 1 | 1.075815916 | NS |
| II | Euarchontoglires | L3 | 0.02068534 | 3.512397464 | * |
| II | Euarchontoglires | L4 | 0.072019705 | 2.665570157 | NS |
| II | Euarchontoglires | L5 | 7.34E-05 | 5.143626624 | * |
| II | Euarchontoglires | L6 | 0.783075702 | 0.709501172 | NS |
| II | Euarchontoglires | L7 | 0.149196055 | 3.852265931 | NS |
| II | Euarchontoglires | L8 | 0.072997084 | 0.505294397 | NS |
| III | Microcebus | L1 | 0.599557949 | 0.773409702 | NS |
| III | Microcebus | L2 | 0.65347923 | 1.404098116 | NS |
| III | Microcebus | L3 | 0.239300654 | 0 | NS |
| III | Microcebus | L4 | 1 | 1.08090091 | NS |
| III | Microcebus | L5 | 0.139391688 | 2.03083807 | NS |
| III | Microcebus | L6 | 0.05966636 | 0 | NS |
| III | Microcebus | L7 | 0.000863416 | 21.12196876 | * |
| III | Microcebus | L8 | 0.858584504 | 1.061828028 | NS |
| III | Cheirogaleidae | L1 | 0.738438854 | 1.106498797 | NS |
| III | Cheirogaleidae | L2 | 1 | 0.58598294 | NS |
| III | Cheirogaleidae | L3 | 0.244141262 | 0 | NS |
| III | Cheirogaleidae | L4 | 1 | 0.991686356 | NS |
| III | Cheirogaleidae | L5 | 0.022459394 | 2.797769052 | * |
| III | Cheirogaleidae | L6 | 0.036446006 | 0 | * |
| III | Cheirogaleidae | L7 | 0.473249224 | 1.801677176 | NS |
| III | Cheirogaleidae | L8 | 0.862845602 | 1.06234323 | NS |
| III | Lemuriformes | L1 | 0.126126715 | 1.711041637 | NS |
| III | Lemuriformes | L2 | 1 | 0.58598294 | NS |
| III | Lemuriformes | L3 | 0.244141262 | 0 | NS |
| III | Lemuriformes | L4 | 0.70781866 | 0.45764076 | NS |
| III | Lemuriformes | L5 | 0.244961876 | 1.846231928 | NS |
| III | Lemuriformes | L6 | 0.557916229 | 0.536588812 | NS |
| III | Lemuriformes | L7 | 0.473249224 | 1.801677176 | NS |
| III | Lemuriformes | L8 | 0.486336483 | 0.727516184 | NS |
| III | Strepsirrhini | L1 | 0.341736294 | 1.388352318 | NS |
| III | Strepsirrhini | L2 | 0.235629522 | 0 | NS |
| III | Strepsirrhini | L3 | 0.149105947 | 0 | NS |
| III | Strepsirrhini | L4 | 1 | 0.827775343 | NS |
| III | Strepsirrhini | L5 | 0.027039494 | 2.723303958 | * |
| III | Strepsirrhini | L6 | 0.158318732 | 0.211961531 | NS |
| III | Strepsirrhini | L7 | 0.023682301 | 7.862134352 | * |
| III | Strepsirrhini | L8 | 0.418724128 | 0.737667599 | NS |
| III | Primates | L1 | 0.378036921 | 1.304434046 | NS |
| III | Primates | L2 | 1 | 0.880372724 | NS |
| III | Primates | L3 | 0.33400158 | 0.314531596 | NS |

| | | | | | |
|---|---|---|---|---|---|
| III | Primates | L4 | 1 | 0.677146167 | NS |
| III | Primates | L5 | 0.208412308 | 1.817510061 | NS |
| III | Primates | L6 | 0.603064578 | 0.577774581 | NS |
| III | Primates | L7 | 0.037609801 | 6.424672696 | * |
| III | Primates | L8 | 0.292864474 | 0.70410309 | NS |
| III | Euarchontoglires | L1 | 0.43778144 | 1.285222393 | NS |
| III | Euarchontoglires | L2 | 1 | 1.026178014 | NS |
| III | Euarchontoglires | L3 | 0.490502195 | 0.365883155 | NS |
| III | Euarchontoglires | L4 | 1 | 0.789529233 | NS |
| III | Euarchontoglires | L5 | 0.031199926 | 2.578969404 | * |
| III | Euarchontoglires | L6 | 0.784212934 | 0.675966425 | NS |
| III | Euarchontoglires | L7 | 0.02645609 | 7.496712005 | * |
| III | Euarchontoglires | L8 | 0.05373124 | 0.476759901 | NS |
| IV | Microcebus | L1 | 0.018666674 | 0.320356871 | * |
| IV | Microcebus | L2 | 0.620663774 | 0 | NS |
| IV | Microcebus | L3 | 0.178872667 | 2.319271744 | NS |
| IV | Microcebus | L4 | 0.011397323 | 4.725183056 | * |
| IV | Microcebus | L5 | 0.003146602 | 4.258066949 | * |
| IV | Microcebus | L6 | 0.160297667 | 2.360822469 | NS |
| IV | Microcebus | L7 | 0.066532153 | 6.525217598 | NS |
| IV | Microcebus | L8 | 0.004502107 | 0.209129139 | * |
| IV | Cheirogaleidae | L1 | 0.423509347 | 0.660581046 | NS |
| IV | Cheirogaleidae | L2 | 0.615982034 | 0 | NS |
| IV | Cheirogaleidae | L3 | 0.641054404 | 1.520817335 | NS |
| IV | Cheirogaleidae | L4 | 0.15463796 | 2.510173763 | NS |
| IV | Cheirogaleidae | L5 | 0.008929601 | 3.778456346 | * |
| IV | Cheirogaleidae | L6 | 0.279891927 | 1.905659608 | NS |
| IV | Cheirogaleidae | L7 | 0.005271123 | 14.55792248 | * |
| IV | Cheirogaleidae | L8 | 0.00159493 | 0.144078471 | * |
| IV | Strepsirrhini | L1 | 0.423509347 | 0.660581046 | NS |
| IV | Strepsirrhini | L2 | 0.615982034 | 0 | NS |
| IV | Strepsirrhini | L3 | 0.15463796 | 2.510173763 | NS |
| IV | Strepsirrhini | L4 | 0.641054404 | 1.520817335 | NS |
| IV | Strepsirrhini | L5 | 0.008929601 | 3.778456346 | * |
| IV | Strepsirrhini | L6 | 0.279891927 | 1.905659608 | NS |
| IV | Strepsirrhini | L7 | 0.005271123 | 14.55792248 | * |
| IV | Strepsirrhini | L8 | 0.00159493 | 0.144078471 | * |
| IV | Euarchontoglires | L1 | 0.32639514 | 0.628058498 | NS |
| IV | Euarchontoglires | L2 | 0.618098024 | 0 | NS |
| IV | Euarchontoglires | L3 | 0.166609481 | 2.411150949 | NS |
| IV | Euarchontoglires | L4 | 0.646860314 | 1.462726106 | NS |
| IV | Euarchontoglires | L5 | 0.010887625 | 3.604343068 | * |
| IV | Euarchontoglires | L6 | 0.291108229 | 1.827590779 | NS |

| | | | | | |
|---|---|---|---|---|---|
| IV | Euarchontoglires | L7 | 0.005823634 | 13.99471981 | * |
| IV | Euarchontoglires | L8 | 0.007244832 | 0.217713337 | * |
| V | Microcebus | L1 | 0.030378763 | 0.402505233 | * |
| V | Microcebus | L2 | 0.639056177 | 1.540117577 | NS |
| V | Microcebus | L3 | 0.68734077 | 1.185755987 | NS |
| V | Microcebus | L4 | 0.400951806 | 1.942592244 | NS |
| V | Microcebus | L5 | 7.52E-05 | 5.963954903 | * |
| V | Microcebus | L6 | 0.755274443 | 0.642130101 | NS |
| V | Microcebus | L7 | 0.087195972 | 5.50148473 | NS |
| V | Microcebus | L8 | 0.095727037 | 0.483502296 | NS |
| V | Cheirogaleidae | L1 | 0.071895308 | 0.463128373 | NS |
| V | Cheirogaleidae | L2 | 0.643635079 | 1.492197245 | NS |
| V | Cheirogaleidae | L3 | 1 | 0.528415862 | NS |
| V | Cheirogaleidae | L4 | 0.090411571 | 2.747924154 | NS |
| V | Cheirogaleidae | L5 | 0.000100487 | 5.718697511 | * |
| V | Cheirogaleidae | L6 | 0.755350011 | 0.622030845 | NS |
| V | Cheirogaleidae | L7 | 0.091566708 | 5.331956428 | NS |
| V | Cheirogaleidae | L8 | 0.095702154 | 0.465577812 | NS |
| V | Lemuriformes | L1 | 0.020443303 | 0.373423421 | * |
| V | Lemuriformes | L2 | 0.648450822 | 1.446927775 | NS |
| V | Lemuriformes | L3 | 0.412054303 | 1.8218735 | NS |
| V | Lemuriformes | L4 | 0.412054303 | 1.8218735 | NS |
| V | Lemuriformes | L5 | 0.000132729 | 5.493199451 | * |
| V | Lemuriformes | L6 | 0.534968427 | 1.367479445 | NS |
| V | Lemuriformes | L7 | 0.096011568 | 5.170981275 | NS |
| V | Lemuriformes | L8 | 0.02774386 | 0.368415675 | * |
| V | Strepsirrhini | L1 | 2.40E-05 | 0.151516509 | * |
| V | Strepsirrhini | L2 | 0.686926106 | 1.188834544 | NS |
| V | Strepsirrhini | L3 | 0.464868639 | 1.489947223 | NS |
| V | Strepsirrhini | L4 | 0.052462442 | 2.963783274 | NS |
| V | Strepsirrhini | L5 | 0.00071577 | 4.282442251 | * |
| V | Strepsirrhini | L6 | 1 | 0.786134599 | NS |
| V | Strepsirrhini | L7 | 0.129022172 | 4.254865199 | NS |
| V | Strepsirrhini | L8 | 0.866690743 | 1.062867091 | NS |
| V | Primates | L1 | 8.50E-06 | 0.143201367 | * |
| V | Primates | L2 | 0.698884187 | 1.129784443 | NS |
| V | Primates | L3 | 0.48279041 | 1.414440894 | NS |
| V | Primates | L4 | 0.06177636 | 2.80763919 | NS |
| V | Primates | L5 | 7.27E-06 | 6.352177702 | * |
| V | Primates | L6 | 0.783685542 | 0.746079242 | NS |
| V | Primates | L7 | 0.139001791 | 4.045163694 | NS |
| V | Primates | L8 | 0.868435204 | 0.884079144 | NS |
| VI | Microcebus | L1 | 5.15E-06 | 0.084303374 | * |

| | | | | | |
|---|---|---|---|---|---|
| VI | Microcebus | L2 | 1 | 0.637424015 | NS |
| VI | Microcebus | L3 | 0.001143493 | 6.092897665 | * |
| VI | Microcebus | L4 | 0.029950195 | 3.541918044 | * |
| VI | Microcebus | L5 | 2.90E-05 | 6.228133083 | * |
| VI | Microcebus | L6 | 0.114630719 | 2.266870568 | NS |
| VI | Microcebus | L7 | 0.447276913 | 1.959080183 | NS |
| VI | Microcebus | L8 | 0.006330668 | 0.28453686 | * |
| VI | Cheirogaleidae | L1 | 0.001414688 | 0.174726948 | * |
| VI | Cheirogaleidae | L2 | 0.620663774 | 0 | NS |
| VI | Cheirogaleidae | L3 | 0.011397323 | 4.725183056 | * |
| VI | Cheirogaleidae | L4 | 0.052153082 | 3.409992366 | NS |
| VI | Cheirogaleidae | L5 | 0.003146602 | 4.258066949 | * |
| VI | Cheirogaleidae | L6 | 0.303353589 | 1.755126534 | NS |
| VI | Cheirogaleidae | L7 | 0.066532153 | 6.525217598 | NS |
| VI | Cheirogaleidae | L8 | 0.047817828 | 0.383861252 | * |
| VI | Lemuriformes | L1 | 0.000850102 | 0.201785063 | * |
| VI | Lemuriformes | L2 | 1 | 0.697668509 | NS |
| VI | Lemuriformes | L3 | 0.021576989 | 3.914917839 | * |
| VI | Lemuriformes | L4 | 0.400951806 | 1.942592244 | NS |
| VI | Lemuriformes | L5 | 0.002100454 | 4.179814781 | * |
| VI | Lemuriformes | L6 | 0.100789267 | 2.513842659 | NS |
| VI | Lemuriformes | L7 | 0.087195972 | 5.50148473 | NS |
| VI | Lemuriformes | L8 | 0.040269255 | 0.396057625 | * |
| VI | Primates | L1 | 0.001365148 | 0.181947576 | * |
| VI | Primates | L2 | 0.618098024 | 0 | NS |
| VI | Primates | L3 | 0.046894274 | 3.549730176 | * |
| VI | Primates | L4 | 0.046894274 | 3.549730176 | * |
| VI | Primates | L5 | 7.52E-05 | 6.611554045 | * |
| VI | Primates | L6 | 1 | 0.792817611 | NS |
| VI | Primates | L7 | 0.362871152 | 2.625352524 | NS |
| VI | Primates | L8 | 0.159603153 | 0.506591385 | NS |
| VII | Microcebus | L1 | 2.55E-08 | 0.034156612 | * |
| VII | Microcebus | L2 | 1 | 0.541547011 | NS |
| VII | Microcebus | L3 | 0.464868639 | 1.489947223 | NS |
| VII | Microcebus | L4 | 0.254291734 | 2.165481451 | NS |
| VII | Microcebus | L5 | 0.266680742 | 1.689788833 | NS |
| VII | Microcebus | L6 | 0.775322759 | 1.111936426 | NS |
| VII | Microcebus | L7 | 0.129022172 | 4.254865199 | NS |
| VII | Microcebus | L8 | 0.002085463 | 2.714793637 | * |
| VII | Cheirogaleidae | L1 | 1.74E-06 | 0.044662352 | * |
| VII | Cheirogaleidae | L2 | 1 | 0.697668509 | NS |
| VII | Cheirogaleidae | L3 | 0.400951806 | 1.942592244 | NS |
| VII | Cheirogaleidae | L4 | 0.400951806 | 1.942592244 | NS |

| | | | | | |
|---|---|---|---|---|---|
| VII | Cheirogaleidae | L5 | 0.350375001 | 1.756181635 | NS |
| VII | Cheirogaleidae | L6 | 1 | 1.026331801 | NS |
| VII | Cheirogaleidae | L7 | 0.087195972 | 5.50148473 | NS |
| VII | Cheirogaleidae | L8 | 0.023859018 | 2.279526122 | * |
| VII | Lemuriformes | L1 | 4.33E-06 | 0 | * |
| VII | Lemuriformes | L2 | 1 | 0.961626474 | NS |
| VII | Lemuriformes | L3 | 1 | 0.751497909 | NS |
| VII | Lemuriformes | L4 | 1 | 0.751497909 | NS |
| VII | Lemuriformes | L5 | 0.283757836 | 1.877936792 | NS |
| VII | Lemuriformes | L6 | 1 | 0.89521013 | NS |
| VII | Lemuriformes | L7 | 0.051567475 | 7.634764362 | NS |
| VII | Lemuriformes | L8 | 0.004513415 | 3.341919594 | * |
| VII | Strepsirrhini | L1 | 3.75E-07 | 0 | * |
| VII | Strepsirrhini | L2 | 1 | 0.796068086 | NS |
| VII | Strepsirrhini | L3 | 0.191404253 | 2.233483489 | NS |
| VII | Strepsirrhini | L4 | 0.659442175 | 1.358129768 | NS |
| VII | Strepsirrhini | L5 | 0.052120082 | 2.626801306 | NS |
| VII | Strepsirrhini | L6 | 1 | 0.735902944 | NS |
| VII | Strepsirrhini | L7 | 0.382499133 | 2.444033741 | NS |
| VII | Strepsirrhini | L8 | 0.010547438 | 2.589423132 | * |
| VIII | Microcebus | L1 | 6.98E-06 | 0.116249456 | * |
| VIII | Microcebus | L2 | 0.221716232 | 2.077946597 | NS |
| VIII | Microcebus | L3 | 0.010883429 | 4.14732344 | * |
| VIII | Microcebus | L4 | 0.447988558 | 1.572978068 | NS |
| VIII | Microcebus | L5 | 0.002061373 | 3.875696121 | * |
| VIII | Microcebus | L6 | 0.015464893 | 3.047590269 | * |
| VIII | Microcebus | L7 | 0.119272702 | 4.484877146 | NS |
| VIII | Microcebus | L8 | 0.056814027 | 0.450641934 | NS |
| VIII | Cheirogaleidae | L1 | 9.65E-06 | 0.089889187 | * |
| VIII | Cheirogaleidae | L2 | 0.643635079 | 1.492197245 | NS |
| VIII | Cheirogaleidae | L3 | 0.000104452 | 8.372947586 | * |
| VIII | Cheirogaleidae | L4 | 0.090411571 | 2.747924154 | NS |
| VIII | Cheirogaleidae | L5 | 0.122752815 | 2.173741063 | NS |
| VIII | Cheirogaleidae | L6 | 0.021959502 | 3.039102979 | * |
| VIII | Cheirogaleidae | L7 | 0.091566708 | 5.331956428 | NS |
| VIII | Cheirogaleidae | L8 | 0.040902781 | 0.381790293 | * |
| VIII | Strepsirrhini | L1 | 0.394730787 | 0.714197389 | NS |
| VIII | Strepsirrhini | L2 | 1 | 0.602273896 | NS |
| VIII | Strepsirrhini | L3 | 0.432402125 | 1.664726416 | NS |
| VIII | Strepsirrhini | L4 | 0.001574896 | 5.707183279 | * |
| VIII | Strepsirrhini | L5 | 0.773874441 | 1.123306673 | NS |
| VIII | Strepsirrhini | L6 | 0.558552858 | 0.551926979 | NS |
| VIII | Strepsirrhini | L7 | 0.014355874 | 9.705251077 | * |

| | | | | | |
|---|---|---|---|---|---|
| VIII | Strepsirrhini | L8 | 0.16139343 | 0.56545591 | NS |
| VIII | Euarchontoglires | L1 | 0.014350159 | 0.39755861 | * |
| VIII | Euarchontoglires | L2 | 0.692873388 | 1.158641072 | NS |
| VIII | Euarchontoglires | L3 | 0.015224141 | 3.805621363 | * |
| VIII | Euarchontoglires | L4 | 0.00322861 | 4.911569644 | * |
| VIII | Euarchontoglires | L5 | 0.583854756 | 1.291059248 | NS |
| VIII | Euarchontoglires | L6 | 0.56436128 | 0.482199481 | NS |
| VIII | Euarchontoglires | L7 | 0.001663663 | 17.31974159 | * |
| VIII | Euarchontoglires | L8 | 0.24301651 | 0.634328767 | NS |
| IX | Microcebus | L1 | 0.514670006 | 0.784126754 | NS |
| IX | Microcebus | L2 | 0.686926106 | 1.188834544 | NS |
| IX | Microcebus | L3 | 0.254291734 | 2.165481451 | NS |
| IX | Microcebus | L4 | 0.464868639 | 1.489947223 | NS |
| IX | Microcebus | L5 | 0.266680742 | 1.689788833 | NS |
| IX | Microcebus | L6 | 0.56148521 | 0.494849356 | NS |
| IX | Microcebus | L7 | 0.018671166 | 8.698797503 | * |
| IX | Microcebus | L8 | 0.310699831 | 0.655573144 | NS |
| IX | Cheirogaleidae | L1 | 0.875414447 | 0.906550113 | NS |
| IX | Cheirogaleidae | L2 | 1 | 1.050479644 | NS |
| IX | Cheirogaleidae | L3 | 0.720601987 | 1.313196545 | NS |
| IX | Cheirogaleidae | L4 | 0.720601987 | 1.313196545 | NS |
| IX | Cheirogaleidae | L5 | 0.424454159 | 1.476443043 | NS |
| IX | Cheirogaleidae | L6 | 0.406230094 | 0.436854737 | NS |
| IX | Cheirogaleidae | L7 | 0.025046581 | 7.675606944 | * |
| IX | Cheirogaleidae | L8 | 0.6284663 | 0.800898826 | NS |
| IX | Strepsirrhini | L1 | 0.873283075 | 1.077077758 | NS |
| IX | Strepsirrhini | L2 | 0.704944597 | 1.102252841 | NS |
| IX | Strepsirrhini | L3 | 0.492069168 | 1.379197321 | NS |
| IX | Strepsirrhini | L4 | 0.492069168 | 1.379197321 | NS |
| IX | Strepsirrhini | L5 | 0.598160635 | 1.223929713 | NS |
| IX | Strepsirrhini | L6 | 0.403995976 | 0.458547776 | NS |
| IX | Strepsirrhini | L7 | 0.022362951 | 8.056757757 | * |
| IX | Strepsirrhini | L8 | 0.324700198 | 0.675463956 | NS |

**Table S11 – Subfamily branches with evidence of episodic positive selection.** Branch site model parameters for the null hypothesis (0) and alternative hypothesis (A).

| | | | | Null Hypothesis | | | | Alternative Hypothesis | | | | | | | | |
|---|---|---|---|---|---|---|---|---|---|---|---|---|---|---|---|---|
| **Subfamily** | **Node** | **$lnL_0$** | **$lnL_A$** | **$p_1$** | **$p_2$** | **$\omega_1$** | **$\omega_2$** | **$p_1$** | **$p_2$** | **$p_3$** | **$\omega_1$** | **$\omega_2$** | **$\omega_3$** | **LRT** | **p-value** | **q-value** |
| **I** | 92 | -18074.62 | -18047.69 | 0.58 | 0.42 | 0.17 | 1.00 | 0.57 | 0.41 | 0.02 | 0.17 | 1.00 | 999.00 | 53.85 | 2.1595E-13 | 0.000235849 |
| | 87 | -18073.86 | -18061.65 | 0.38 | 0.27 | 0.17 | 1.00 | 0.57 | 0.41 | 0.02 | 0.17 | 1.00 | 999.00 | 24.42 | 7.7274E-07 | 0.000471698 |
| | 88 | -18074.62 | -18066.56 | 0.58 | 0.42 | 0.17 | 1.00 | 0.58 | 0.41 | 0.02 | 0.17 | 1.00 | 999.00 | 16.11 | 0.000059892 | 0.000707547 |
| | 107 | -18065.84 | -18059.24 | 0.00 | 0.00 | 0.17 | 1.00 | 0.55 | 0.39 | 0.07 | 0.17 | 1.00 | 47.93 | 13.19 | 0.00028168 | 0.000943396 |
| | 97 | -18074.62 | -18068.61 | 0.58 | 0.42 | 0.17 | 1.00 | 0.57 | 0.41 | 0.02 | 0.17 | 1.00 | 454.04 | 12.02 | 0.00052634 | 0.001179245 |
| | 4 | -18074.62 | -18069.45 | 0.58 | 0.42 | 0.17 | 1.00 | 0.58 | 0.41 | 0.01 | 0.17 | 1.00 | 999.00 | 10.33 | 0.0013121 | 0.001415094 |
| **II** | 117 | -23372.89 | -23360.85 | 0.47 | 0.47 | 0.17 | 1.00 | 0.49 | 0.49 | 0.02 | 0.17 | 1.00 | 999.00 | 24.09 | 9.1947E-07 | 0.000175439 |
| | 51 | -23372.91 | -23362.29 | 0.50 | 0.50 | 0.17 | 1.00 | 0.48 | 0.48 | 0.03 | 0.17 | 1.00 | 999.00 | 21.23 | 4.0734E-06 | 0.000350877 |
| | 134 | -23372.88 | -23363.74 | 0.47 | 0.46 | 0.17 | 1.00 | 0.50 | 0.49 | 0.01 | 0.17 | 1.00 | 999.00 | 18.28 | 0.000019096 | 0.000526316 |
| | 138 | -23372.91 | -23365.56 | 0.50 | 0.50 | 0.17 | 1.00 | 0.49 | 0.49 | 0.02 | 0.17 | 1.00 | 842.87 | 14.70 | 0.00012603 | 0.000701754 |
| | 54 | -23372.91 | -23365.65 | 0.50 | 0.50 | 0.17 | 1.00 | 0.50 | 0.49 | 0.01 | 0.17 | 1.00 | 999.00 | 14.51 | 0.00013919 | 0.000877193 |
| **III** | **NA** | | | | | | | | | | | | | | | |
| **IV** | 16 | -7462.33 | -7441.97 | 0.60 | 0.40 | 0.15 | 1.00 | 0.59 | 0.39 | 0.01 | 0.15 | 1.00 | 999.00 | 40.73 | 1.7511E-10 | 0.000543478 |
| **V** | 118 | -30877.80 | -30850.07 | 0.37 | 0.32 | 0.25 | 1.00 | 0.52 | 0.45 | 0.04 | 0.25 | 1.00 | 999.00 | 55.46 | 9.5179E-14 | 0.000132979 |
| | 117 | -30878.67 | -30858.63 | 0.48 | 0.42 | 0.25 | 1.00 | 0.51 | 0.45 | 0.04 | 0.25 | 1.00 | 999.00 | 40.09 | 2.4267E-10 | 0.000265957 |
| | 116 | -30878.92 | -30866.04 | 0.53 | 0.47 | 0.25 | 1.00 | 0.52 | 0.46 | 0.02 | 0.25 | 1.00 | 999.00 | 25.75 | 3.8893E-07 | 0.000398936 |
| | 135 | -30878.36 | -30867.02 | 0.45 | 0.40 | 0.25 | 1.00 | 0.52 | 0.46 | 0.02 | 0.25 | 1.00 | 205.39 | 22.69 | 1.9072E-06 | 0.000531915 |
| | 47 | -30878.92 | -30872.57 | 0.53 | 0.47 | 0.25 | 1.00 | 0.53 | 0.47 | 0.00 | 0.25 | 1.00 | 999.00 | 12.71 | 0.00036468 | 0.000664894 |
| | 166 | -30875.48 | -30869.39 | 0.29 | 0.26 | 0.25 | 1.00 | 0.52 | 0.46 | 0.01 | 0.25 | 1.00 | 64.71 | 12.18 | 0.00048219 | 0.000797872 |
| | 107 | -30874.20 | -30868.45 | 0.22 | 0.20 | 0.25 | 1.00 | 0.49 | 0.44 | 0.07 | 0.25 | 1.00 | 20.96 | 11.50 | 0.00069764 | 0.000930851 |
| | 183 | -30877.89 | -30872.52 | 0.41 | 0.37 | 0.25 | 1.00 | 0.50 | 0.45 | 0.05 | 0.25 | 1.00 | 21.46 | 10.73 | 0.0010522 | 0.00106383 |
| **VI** | 34 | -5447.78 | -5424.80 | 0.47 | 0.52 | 0.16 | 1.00 | 0.45 | 0.51 | 0.05 | 0.16 | 1.00 | 332.48 | 45.95 | 1.2159E-11 | 0.000714286 |
| | 33 | -5447.78 | -5441.02 | 0.47 | 0.52 | 0.16 | 1.00 | 0.47 | 0.51 | 0.03 | 0.16 | 1.00 | 194.44 | 13.51 | 0.00023716 | 0.001428571 |
| | 22 | -5447.42 | -5441.43 | 0.39 | 0.42 | 0.16 | 1.00 | 0.48 | 0.51 | 0.01 | 0.16 | 1.00 | 152.05 | 11.98 | 0.0005379 | 0.002142857 |
| | 1 | -5447.78 | -5442.40 | 0.48 | 0.52 | 0.16 | 1.00 | 0.46 | 0.52 | 0.02 | 0.16 | 1.00 | 66.15 | 10.75 | 0.0010451 | 0.002857143 |
| **VII** | 97 | -8685.16 | -8657.49 | 0.34 | 0.33 | 0.16 | 1.00 | 0.50 | 0.48 | 0.01 | 0.16 | 1.00 | 999.00 | 55.34 | 1.0142E-13 | 0.000403226 |
| | 81 | -8685.66 | -8672.55 | 0.49 | 0.50 | 0.16 | 1.00 | 0.49 | 0.50 | 0.01 | 0.16 | 1.00 | 999.00 | 26.21 | 3.0595E-07 | 0.000806452 |
| | 123 | -8685.58 | -8676.83 | 0.44 | 0.45 | 0.16 | 1.00 | 0.49 | 0.50 | 0.01 | 0.16 | 1.00 | 573.41 | 17.51 | 0.000028589 | 0.001209677 |
| | 66 | -8685.05 | -8678.93 | 0.45 | 0.46 | 0.15 | 1.00 | 0.47 | 0.46 | 0.06 | 0.15 | 1.00 | 14.65 | 12.22 | 0.00047213 | 0.001612903 |
| | 124 | -8685.51 | -8679.65 | 0.43 | 0.44 | 0.15 | 1.00 | 0.49 | 0.50 | 0.01 | 0.16 | 1.00 | 703.40 | 11.73 | 0.00061624 | 0.002016129 |
| **IX** | 115 | -17195.12 | -17138.91 | 0.34 | 0.23 | 0.13 | 1.00 | 0.53 | 0.37 | 0.10 | 0.13 | 1.00 | 363.24 | 112.43 | 2.8734E-26 | 0.000223214 |
| | 171 | -17214.08 | -17185.27 | 0.60 | 0.40 | 0.14 | 1.00 | 0.59 | 0.40 | 0.02 | 0.14 | 1.00 | 575.62 | 57.64 | 3.1553E-14 | 0.000446429 |
| | 212 | -17211.79 | -17185.54 | 0.55 | 0.37 | 0.14 | 1.00 | 0.57 | 0.37 | 0.06 | 0.14 | 1.00 | 999.00 | 52.49 | 4.3149E-13 | 0.000669643 |
| | 225 | -17214.08 | -17195.79 | 0.60 | 0.40 | 0.14 | 1.00 | 0.58 | 0.39 | 0.02 | 0.14 | 1.00 | 999.00 | 36.59 | 1.4557E-09 | 0.000892857 |
| | 121 | -17212.86 | -17201.05 | 0.54 | 0.36 | 0.14 | 1.00 | 0.58 | 0.39 | 0.04 | 0.14 | 1.00 | 58.91 | 23.63 | 1.1699E-06 | 0.001116071 |
| | 186 | -17214.08 | -17202.97 | 0.60 | 0.40 | 0.14 | 1.00 | 0.60 | 0.40 | 0.01 | 0.14 | 1.00 | 999.00 | 22.23 | 2.4235E-06 | 0.001339286 |
| | 118 | -17207.46 | -17201.14 | 0.46 | 0.31 | 0.13 | 1.00 | 0.55 | 0.37 | 0.08 | 0.13 | 1.00 | 11.40 | 12.66 | 0.00037449 | 0.0015625 |
| | 154 | -17214.05 | -17208.70 | 0.57 | 0.38 | 0.14 | 1.00 | 0.59 | 0.40 | 0.00 | 0.14 | 1.00 | 697.21 | 10.71 | 0.001064 | 0.001785714 |
| | 211 | -17214.08 | -17208.81 | 0.60 | 0.40 | 0.14 | 1.00 | 0.59 | 0.38 | 0.03 | 0.14 | 1.00 | 428.70 | 10.55 | 0.0011615 | 0.002008929 |
| | 22 | -17214.08 | -17209.16 | 0.60 | 0.40 | 0.14 | 1.00 | 0.59 | 0.40 | 0.01 | 0.14 | 1.00 | 999.00 | 9.84 | 0.0017039 | 0.002232143 |
| | 198 | -17214.08 | -17209.38 | 0.60 | 0.40 | 0.14 | 1.00 | 0.59 | 0.40 | 0.02 | 0.14 | 1.00 | 999.00 | 9.42 | 0.0021487 | 0.002455357 |

**Table S12 – Individual sites under selection along specified branches.** Branch site model parameters were used to test sites for positive selection by way of Bayes empirical Bayes. The alignment column (sites) and domains where those sites are located – loop (L) or transmembrane (TM) are provided for sites with posterior probabilities greater than 0.95. NS implies no individual site was significant by Bayes empirical Bayes.

| Subfamily | Node | Sites | Domains |
|---|---|---|---|
| **I** | 92 | 317, 348 | L5,TM5 |
| | 87 | NS | - |
| | 88 | NS | - |
| | 107 | 464, 465 | L8, L8 |
| | 97 | 338 | L5 |
| | 4 | NS | - |
| **II** | 117 | NS | - |
| | 51 | 316, 326, 340 | L5, L5, L5 |
| | 134 | NS | - |
| | 138 | NS | - |
| | 54 | NS | - |
| **III** | **NA** | **-** | - |
| **IV** | 16 | 490 | L8 |
| **V** | 118 | 445, 457, 499, 518 | TM7, L8, L8, L8 |
| | 117 | 462 | L8 |
| | 116 | NS | - |
| | 135 | 273 | L4 |
| | 47 | NS | - |
| | 166 | 256 | T3 |
| | 107 | 242, 390, 400 | TM3, L6, L7 |
| | 183 | 363 | TM5 |
| **VI** | 34 | 438, 440, 443, 444 | TM7 TM7, TM7, TM7 |
| | 33 | NS | - |
| | 22 | NS | - |
| | 1 | NS | - |
| **VII** | 97 | 466, 467, 492 | L8,L8,L8 |
| | 81 | NS | - |
| | 123 | NS | - |
| | 66 | 216 | TM2 |
| | 124 | NS | - |
| **IX** | 115 | 172, 215 , 227 ,238, 322, 369, 373, 433, 434, 437, 438 , 439, 440, 441 ,447, 448, 449, 452, 453, 455, 456. 457, 462, 463, 464, 465, 467 | TM1, TM2, L3, TM3, L5, L6, L6, TM7, TM7, TM7, TM7, TM7, TM7, TM7 TM7, TM7, TM7, TM7, L8, L8, L8, L8, L8, L8, L8, L8, L8 |
| | 171 | 528 | L8 |
| | 212 | 147 | L1 |
| | 225 | 527, 528 | L8, L8 |
| | 121 | 415, 518 | TM6, L8 |
| | 186 | NS | - |
| | 118 | 363, 386 | TM3, L6 |

| | | |
|---|---|---|
| 154 | NS | - |
| 211 | NS | - |
| 22 | NS | - |
| 198 | NS | - |

**Table S13 – Sample information for individuals sequenced for this study.**

| Genus | Species | Sample ID | Sex | Tissue Type | Latitude | Longitude | Locality |
|---|---|---|---|---|---|---|---|
| *Cheirogaleus* | *medius* | DLC3619 | Female | Liver | - | - | Duke Lemur Center |
| *Cheirogaleus* | *sibreei* | 106186 | Female | Ear | -19.62 | 47.68 | Tsinjoarivo |
| *Microcebus* | *murinus* | DLC7028 | Female | Blood | - | - | Duke Lemur Center |
| *Microcebus* | *murinus* | DLC7049 | Female | Thigh Skin | - | - | Duke Lemur Center |
| *Microcebus* | *murinus* | DLC7163 | Female | Thigh Skin | - | - | Duke Lemur Center |
| *Microcebus* | *murinus* | DLC7188 | Female | Thigh Skin | - | - | Duke Lemur Center |
| *Microcebus* | *murinus* | DLC7128 | Male | Blood | - | - | Duke Lemur Center |
| *Microcebus* | *murinus* | DLC7033 | Male | Blood | - | - | Duke Lemur Center |
| *Microcebus* | *murinus* | DLC7032 | Male | Blood | - | - | Duke Lemur Center |
| *Microcebus* | *murinus* | DLC7039 | Male | Thigh Skin | - | - | Duke Lemur Center |
| *Microcebus* | *griseorufus* | RMR66 | Female | Kidney | -23.68 | 44.59 | Beza Mahafaly |
| *Microcebus* | *mittermeieri* | MBB011 | Female | Ear | -14.74 | 49.49 | Anjanaharibe Sud |
| *Microcebus* | *ravelobensis* | RMR55 | Male | Unknown | -16.333333 | 46.783333 | Ankarafantsika |
| *Microcebus* | *tavaratra* | RMR71 | Female | Unknown | -13.05 | 49.05 | Ankarana |
| *Mirza* | *zaza* | DLC2301 | Female | Heart | - | - | Duke Lemur Center |

**Table S14– Transcriptome sequences used for TRINITY assembly and SNAP training.**

| SRA ID | Tissue | Replicate | Read Pairs | Trimmed Read Pairs |
|---|---|---|---|---|
| SRR1758989 | cerebellum | 1 | 40282218 | 36203305 |
| SRR1758990 | cerebellum | 2 | 33701327 | 29725813 |
| SRR1758991 | frontal cortex | 1 | 22282358 | 19420162 |
| SRR1758992 | frontal cortex | 2 | 42671438 | 38211712 |
| SRR1758993 | temporal lobe | 1 | 97678744 | 85967715 |
| SRR1758994 | colon | 1 | 94987670 | 75049387 |
| SRR1758995 | kidney | 1 | 20357117 | 17355807 |
| SRR1758996 | kidney | 2 | 39391366 | 34870736 |
| SRR1758997 | liver | 1 | 68821579 | 61258808 |
| SRR1758998 | lung | 1 | 76923918 | 70589216 |
| SRR1758999 | skeletal muscle | 1 | 18183486 | 16943154 |
| SRR1759000 | skeletal muscle | 2 | 52445854 | 46545480 |
| SRR1759001 | skeletal muscle | 3 | 17366481 | 15493101 |
| SRR1759002 | spleen | 1 | 19307314 | 14823668 |
| SRR1759003 | spleen | 2 | 38808580 | 31952914 |

**Table S15 – NCBI records for new sequence data.**

| Genus | species | BioProject | Biosample | SRA | Genome |
|---|---|---|---|---|---|
| *Microcebus* | *griseorufus* | PRJNA512515 | SAMN10707780 | SRR8456524 | Pending |
| *Microcebus* | *mittermeieri* | PRJNA512515 | SAMN10707781 | SRR8456525 | Pending |
| *Microcebus* | *ravelobensis* | PRJNA512515 | SAMN10707782 | SRR8456522,SRR8456518, SRR8456519,SRR8456526, SRR8456527,SRR8456528, SRR8456529,SRR8456530 | Pending |
| *Microcebus* | *tavaratra* | PRJNA512515 | SAMN10707784 | SRR8456520 | Pending |
| *Mirza* | *zaza* | PRJNA512515 | SAMN10707785 | SRR8456521 | Pending |

**Table S16 – BUSCO statistics for all genomes analyzed.** New mouse and dwarf lemur genomes are highlighted in gray.

| Species | Assembly | Complete | Single | Duplicate | Fragment | Missing |
|---|---|---|---|---|---|---|
| *Cheirogaleus medius* | Cmed_1.0 | 93.40% | 92.70% | 0.70% | 3.4%% | 3.20% |
| *Cheirogaleus sibreei* | Csib_1.0 | 72.80% | 71.40% | 1.40% | 19.20% | 8.00% |
| *Daubentonia madagascarensis* | DauMad_1.0 | 9.60% | 9.50% | 0.10% | 35.40% | 55.00% |
| *Eulemur flavifrons* | Eflavifronsk33QCA | 92.10% | 91.50% | 0.60% | 4.50% | 3.40% |
| *Eulemur macaco* | Emacaco_refEf_BWA_oneround | 93.00% | 92.50% | 0.50% | 4.40% | 2.60% |
| *Galeopterus variegatus* | G_variegatus-3.0.2 | 89.00% | 87.50% | 1.50% | 5.90% | 5.10% |
| *Microcebus griseorufus* | Mgri_1.0 | 48.70% | 47.30% | 1.40% | 31.90% | 19.40% |
| *Microcebus mittermeieri* | Mmit_1.0 | 49.10% | 47.00% | 2.10% | 32.30% | 18.60% |
| Microcebus murinus | Mmur_2.0 | 94.90% | 93.20% | 1.70% | 2.50% | 2.60% |
| *Microcebus murinus* | Mmur_3.0 | 95.00% | 93.30% | 1.70% | 2.30% | 2.70% |
| *Microcebus murinus* | Mmur_DLC7028 | 82.40% | 81.20% | 1.20% | 10.90% | 6.70% |
| *Microcebus murinus* | Mmur_DLC7032 | 83.40% | 80.60% | 2.80% | 10.30% | 6.30% |
| *Microcebus murinus* | Mmur_DLC7033v1 | 85.70% | 83.00% | 2.70% | 8.40% | 5.90% |
| *Microcebus murinus* | Mmur_DLC7033v2 | 88.90% | 86.60% | 2.30% | 6.60% | 4.50% |
| *Microcebus murinus* | Mmur_DLC7039 | 83.80% | 82.10% | 1.70% | 9.90% | 6.30% |
| *Microcebus murinus* | Mmur_DLC7049 | 75.90% | 74.80% | 1.10% | 14.00% | 10.10% |
| *Microcebus murinus* | Mmur_DLC7128 | 86.20% | 84.40% | 1.80% | 8.00% | 5.80% |
| *Microcebus murinus* | Mmur_DLC7163 | 84.10% | 82.20% | 1.90% | 9.50% | 6.40% |
| *Microcebus murinus* | Mmur_DLC7188 | 81.20% | 79.60% | 1.60% | 11.40% | 7.40% |
| *Microcebus ravelobensis* | Mrav_1.0 | 44.00% | 42.90% | 1.10% | 33.20% | 22.80% |
| *Microcebus tavaratra* | Mtav_1.0 | 57.80% | 56.00% | 1.80% | 27.70% | 14.50% |
| *Mirza zaza* | Mzaz_1.0 | 73.90% | 72.80% | 1.10% | 18.40% | 7.70% |
| *Otolemur garnetti* | OtoGar3 | 93.80% | 92.60% | 1.20% | 2.50% | 3.70% |
| *Prolemur simus* | Prosim_1.0 | 95.20% | 94.40% | 0.80% | 2.60% | 2.20% |
| *Propithecus coquereli* | Pcoq_1.0 | 91.30% | 90.30% | 1.00% | 4.40% | 4.30% |
| *Tupaia chinensis* | TupChi_1.0 | 91.40% | 90.10% | 1.30% | 4.20% | 4.40% |
